# Disrupted development of sensory systems and the cerebellum in a zebrafish *ebf3a* mutant

**DOI:** 10.1101/2024.09.10.612287

**Authors:** Nghi D. P. Dang, Claire L. Conklin, Thinh Q. Truong, Michael D. Vivian, Jun Wang, Holly R. Thomas, John M. Parant, Nan Cher Yeo, Summer B. Thyme

## Abstract

Mutations in the transcription factor *EBF3* results in a neurodevelopmental disorder, and studies in animal models indicate that it has a critical role in neuronal differentiation. The molecular pathways and neuron types disrupted by its loss, however, have not been thoroughly investigated. Nor have the outcomes of these changes on behavior and brain activity. Here, we generated and characterized a zebrafish *ebf3a* loss-of-function mutant. We discovered morphological and neural phenotypes, including an overall smaller brain size, particularly in the hypothalamus, cerebellum, and hindbrain. Brain function was also compromised, with activity strongly increased in the cerebellum and abnormal behavior at baseline and in response to visual and acoustic stimuli. From RNA-sequencing of developing larvae, notable changes included significant downregulation of genes that mark olfactory sensory neurons, the lateral line, and cerebellar Purkinje neurons. This study sets the stage for determining which downstream pathways underlie the emergence of the observed phenotypes and establishes multiple strong phenotypes that could form the basis of a drug screen.

## Introduction

Mutations in the transcription factor *EBF3* cause a neurodevelopmental disorder characterized mainly by intellectual disability and hypotonia. This syndrome, also known as HADDS, was independently discovered in 2017 by three large sequencing projects, which together identified over twenty individuals (Chao et al. 2017; Harms et al. 2017; Sleven et al. 2017). Prior to the discovery of these coding variants, *EBF3* was hypothesized to contribute to intellectual phenotypes in individuals with terminal 10q deletions (Courtens et al. 2006; Faria et al. 2016), and many more cases have since been recognized (Blackburn et al. 2017; Tanaka et al. 2017; Jiménez de la Peña et al. 2021; Ciaccio et al. 2023). Sequence variation in *EBF3* includes heterozygous nonsense, splice, frameshift, and predicted deleterious missense mutations. While features of autism are present in only a subset of those with *EBF3* coding mutations, noncoding *de novo* variants in an *EBF3* enhancer were found in families with autism (Padhi et al. 2021). Other phenotypes found only in some individuals with *EBF3* mutations include motor coordination issues, ataxia, facial dysmorphism, pain insensitivity, ophthalmologic problems, short stature, restless sleep, and febrile seizures. Neuroimaging has uncovered cerebellar hypoplasia in some cases, although no other consistent structural abnormality has emerged (Harms et al. 2017; Tanaka et al. 2017; D’Arrigo et al. 2020).

Orthologs of *EBF3* are known to function in brain development in multiple species. While *Drosophila melanogaster* and *Caenorhabditis elegans* each only have a single Collier/Olf/EBF (COE) transcription factor, compared to four in most vertebrates, the protein has neurodevelopmental roles in both species (Crozatier et al. 1996; Prasad et al. 1998; Crozatier and Vincent 2008). In mice, homozygous disruption of *Ebf3* reduced survival, mating efficiency, and growth, and resulted in a failure of olfactory neurons to project to the olfactory bulb (Wang et al. 2004). Also in mice, *Ebf3* was shown to be a direct downstream target of the transcriptional repressor *Arx* (Fulp et al. 2008), which itself is involved in a neurodevelopmental syndrome consisting of intellectual disability, seizures, and dystonia (Strømme et al. 2002). In *Xenopus*, the *EBF3* ortholog regulates neuronal differentiation, inducing ectopic neurons when overexpressed in the developing embryo (Pozzoli et al. 2001). In most brain regions and at multiple stages of development, *ebf3* has a similar expression pattern to the *Xenopus* ortholog of mammalian *NEUROD1*, including the neural tube, neural crest cells, olfactory placodes, trigeminal placodes, spinal cord, and retina (Pozzoli et al. 2001; Green and Vetter 2011). Although *ebf3* was shown to be a direct downstream target of the NeuroD transcription factor, it also indirectly regulates *neurod1* expression, suggesting a more complex genetic interaction, possibly with feedback (Green and Vetter 2011).

Although *EBF3* orthologs have been studied in several species, a zebrafish mutant model has not been characterized. The genetically tractable zebrafish has proven useful for the study of the developmental and behavioral impacts of knocking out genes involved in neurodevelopmental disorders (Hoffman et al. 2016; Kozol et al. 2021; Zoodsma et al. 2022; Capps et al. 2024 Jan 20). Many features of zebrafish make it a powerful model to study these disorders, including optical transparency, external fertilization, and conservation of brain organization and many neuron types (Bae et al. 2009; Mueller 2012; Hashikawa et al. 2020; Porter and Mueller 2020; Pandey et al. 2023). Large-scale pharmacological screens for behavioral outputs or using reporter lines are also possible using larval zebrafish once a mutant phenotype has been established (Kokel et al. 2010; Rihel et al. 2010; Wang et al. 2015).

Here, we describe the generation and characterization of the zebrafish orthologs of *EBF3*. In zebrafish larvae, expression of the *ebf3a* ortholog of *EBF3* was found in the posterior tuberculum area of the diencephalon, including in dopamine neurons, the midbrain-hindbrain boundary, olfactory bulb, neuromast hair cells, and hindbrain, including in cerebellar Purkinje cells (Li et al. 2010; Takeuchi et al. 2017; Lush et al. 2019). Because of its extensive expression and importance to brain development of other species, we expected numerous neural and behavioral phenotypes. Using brain imaging and larval behavioral profiling, we found changes in brain size, brain activity, baseline behavior, and stimulus-driven responses. As *EBF3* is a transcription factor, we also expected gene expression differences to likely drive the observed phenotypes.

Previous work in *Xenopus* revealed a small number of direct and indirect *ebf3* target genes, but RNA-sequencing datasets only exist for human cell lines (Harms et al. 2017), not for developing vertebrate embryos with manipulated levels of their *EBF3* ortholog. Thus, we collected RNA-sequencing data from our *ebf3a* zebrafish mutant, which nominated disrupted pathways and impacted cell types. Both cerebellar Purkinje cells and the lateral line were strongly affected by *ebf3a* loss. This study provides a starting point for further in-depth investigations into the *ebf3a* downstream target genes that drive these phenotypes and drug screens to discover molecules that ameliorate them.

## Results

### Zebrafish *ebf3a* is the functional ortholog of human *EBF3*

There are two putative orthologs of human *EBF3* in the zebrafish genome: *ebf3a* and *ebf3b*. A common isoform of the Ebf3a protein (548 aa) has 88.8% identity with human EBF3 (596 aa) while Ebf3b (514 aa) has 62.7% identity. To investigate the function of these orthologs, we used CRISPR/Cas9 to establish loss-of-function mutant lines for *ebf3a* and *ebf3b* (Figure 1A, Supplementary Figure S1A). Both mutations are predicted to cause frameshift of the genes and a premature stop codon, leading to protein truncation at amino acids 128 and 124 for Ebf3a and Ebf3b, respectively (Figure 1B, Supplementary Figure S1B). Using RT-qPCR, we confirmed a significant reduction of the *ebf3a* transcript in the corresponding mutant (Figure 1C). For *ebf3b*, Quantification Cycle (Cq) values were consistently over 30 in wild-type and homozygous samples, indicating a low mRNA level (data not shown). This result is corroborated by a recent single-cell RNA-seq atlas, which revealed high expression of *ebf3a* in neurons across several tissue types (brain, eye, olfactory, and lateral line) and almost no expression of *ebf3b* in the whole organism from 0-5 days post-fertilization (dpf) (Supplementary Figure S1C) (Sur et al. 2023). Its lower expression level and protein identity indicate that *ebf3b* may not have retained function following zebrafish genome duplication.

**Figure 1.**
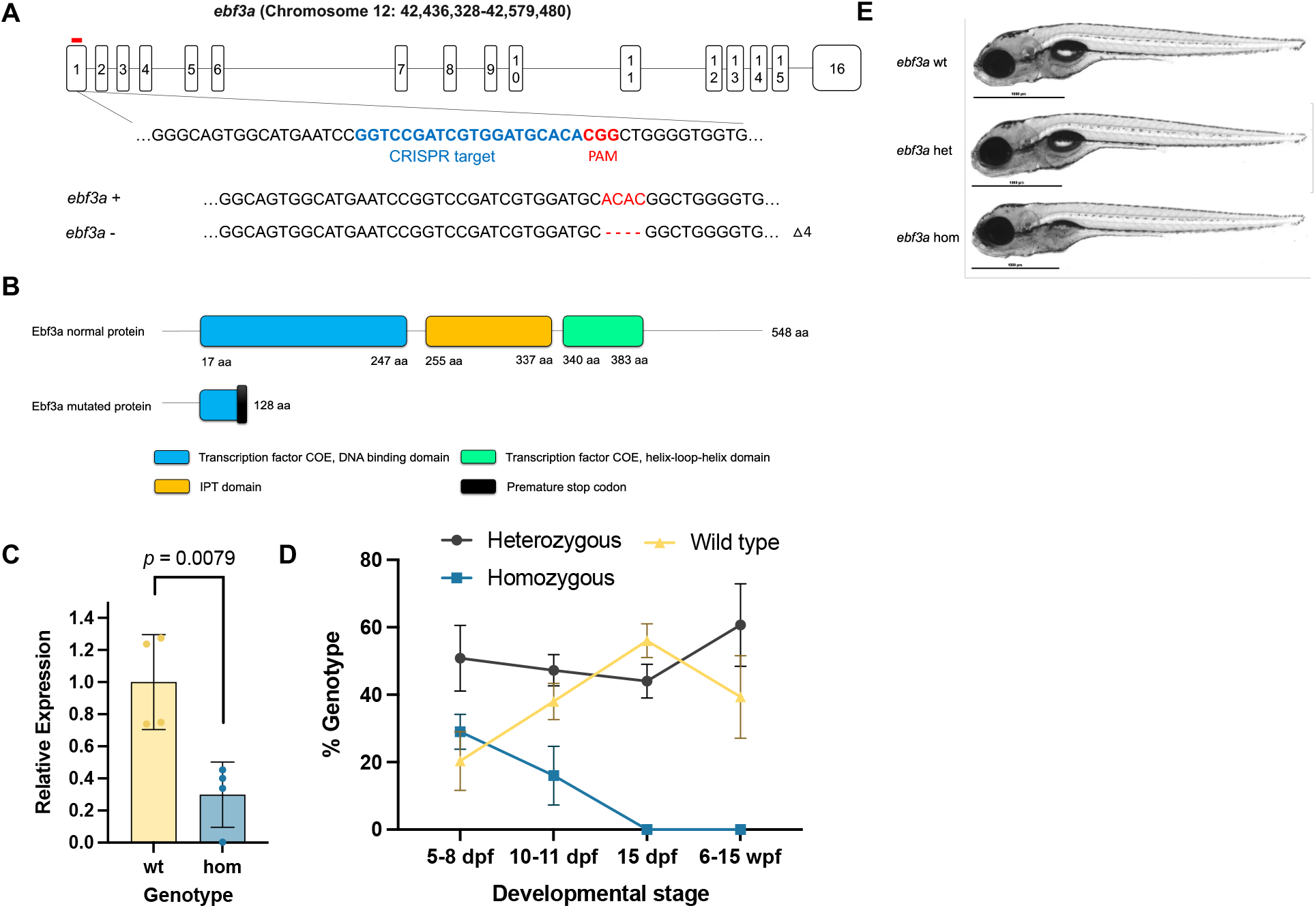
Generation and developmental characterization of zebrafish *ebf3a* mutants. (**A**) Schematic of zebrafish *ebf3a* gene transcript and CRISPR gRNA targeting sequence. (**B**) Schematic of zebrafish wild-type and mutant Ebf3a proteins. Domains are annotated based on ensemble Pfam database. (**C**) Expression level of *ebf3a* in wild-type and homozygous mutant animals at 5 dpf, normalized to the wt. (**D**) Survival of *ebf3a* homozygous mutants and siblings. (**E**) Representative photos of *ebf3a* homozygous mutants and respective siblings at 5 dpf.

First, we evaluated the morphology and survival of these two mutants. We failed to detect adult *ebf3a* F2 homozygous mutants and determined that lethality occurred as early as 10 dpf (Figure 1D, Supplementary Table S1). At larval stages, we found that the swim bladders of *ebf3a* homozygous mutants were not inflating (Figure 1E), while the *ebf3b* larvae were normal (Supplementary Figure S1D). In one clutch, the absent swim bladder was observed in 20 out of 35 homozygous larvae at 6 dpf and only 3 out of 133 heterozygous and wild-type siblings. In summary, we generated an *ebf3a* mutant model with developmental phenotypes, further supporting the designation of *ebf3a* as the functional ortholog.

### Zebrafish *ebf3a* mutants have altered brain morphology, brain activity, and behavior

Given the known role of its orthologs in the nervous system and neural expression, we next assessed the *ebf3a* mutant using phosphorylated Erk (pErk) brain activity mapping (Randlett et al. 2015; Thyme et al. 2019). In this approach, 6 dpf larvae are stained for pErk and total-Erk (tErk), image stacks registered to the Z-Brain atlas (Randlett et al. 2015), and the differences in pErk / tErk ratio between mutant and sibling groups are a proxy for activity. In addition, the registration process generates a deformation matrix that can be similarly compared to identify brain areas (Figure 2A) with increased or decreased size (Jefferis et al. 2007; Thyme et al. 2019). The *ebf3a* homozygous mutants showed altered brain activity and structure compared to their wild-type siblings, with replicable outcomes from two separate clutches (Figure 2B). The size of the brain was generally smaller, particularly in the hypothalamus, cerebellum, and hindbrain, although an area of the caudal rhombencephalon was increased in size. Increased brain activity was pronounced in the cerebellum, as well as in the optic tectum neuropil in the run with stronger signal, and reduced activity was observed in the telencephalon, habenula, mesencephalon, and caudal rhombencephalon. Examination of the raw image stacks confirmed the smaller size of the cerebellum (Figure 2, C and D). To ensure that these phenotypes were not caused by the lack of a swim bladder, we separately analyzed only those where it had fully inflated (Figure 2, E and F). Although the phenotype was milder, as would be expected from the smaller N and selection of least impacted animals, the reduced cerebellar size and increased caudal rhombencephalon remained, as did the increased activity in the cerebellum and reduced in the habenula. Although these strong phenotypes were absent in heterozygous siblings, mild, repeatable activity differences in the cerebellum and habenula remained (Figure 2G).

**Figure 2.**
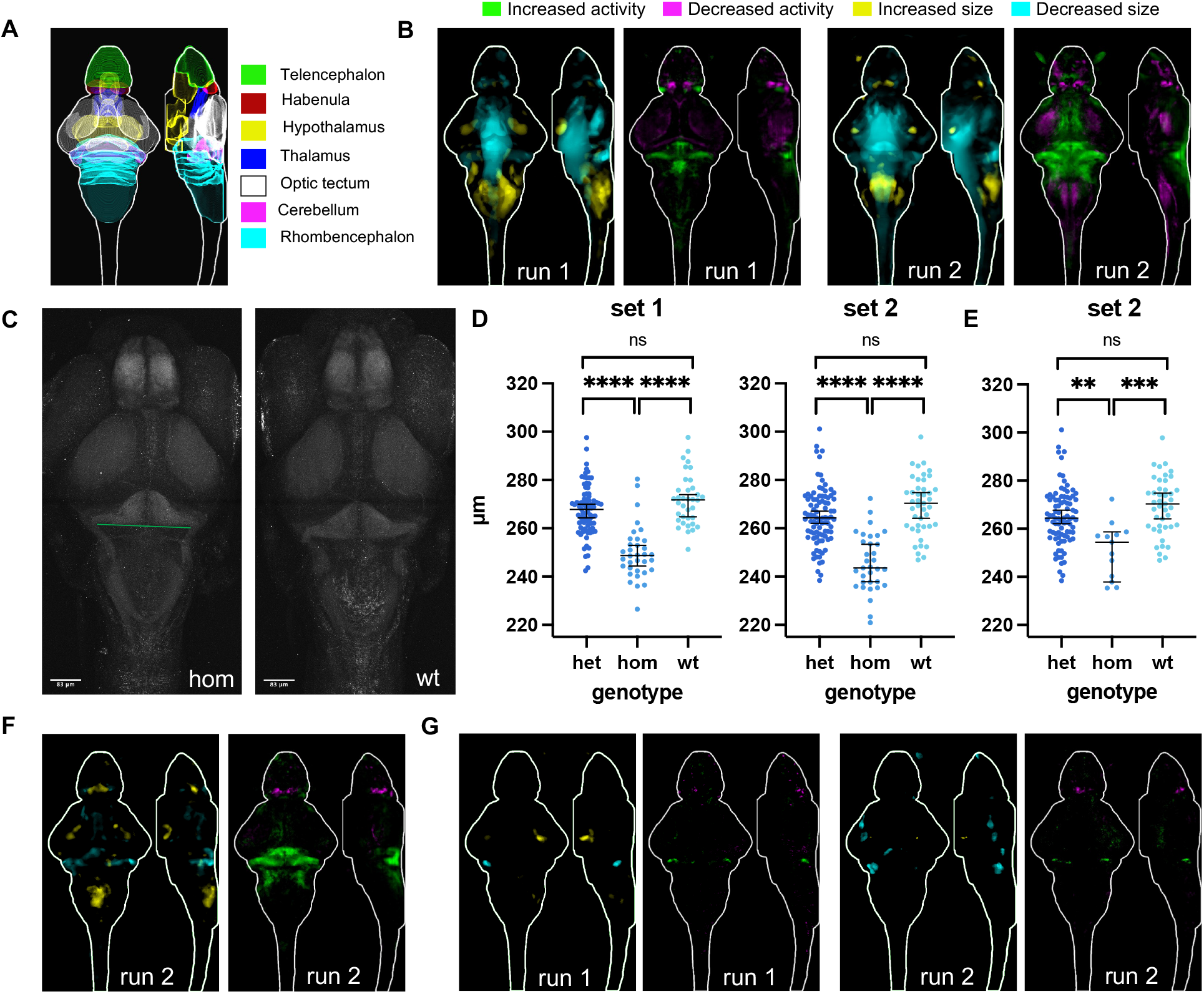
Brain structure and activity phenotypes of zebrafish *ebf3a* mutants. (**A**) Schematic of major zebrafish brain regions based on the Z-Brain atlas (Randlett et al. 2015). (**B**) Brain structure and activity maps for two runs from separate clutches, comparing homozygous mutants to wild-type siblings. A sum-of-slices intensity projection is shown (Z- and X-axes) of the activity signals inside the brain (white outline). Run 1 N = 34 homozygous compared to 36 wild type; Run 2 N = 36 homozygous compared to 46 wild type. (**C**) Example of measuring the width of the hindbrain in the cerebellum region on the raw confocal stacks. The green line represents the measurement that was made on maximum-intensity projections of the tErk stain. (**D**) Measurement of the hindbrain width in all animals from the two runs. The p-values for both runs are <0.0001 for comparisons of homozygous mutants to both wild-type and heterozygous siblings. P-values were calculated using the Brown-Forsythe and Welch ANOVA with Dunnett’s T3 multiple comparisons test (GraphPad Prism 10). (**E**) Measurement of the hindbrain width for only those with swim bladders from run 2. The p-values are 0.0002 for homozygous compared to wild type and 0.0024 for homozygous compared to heterozygous. (**F**) Brain structure and activity maps including only those animals with swim bladders from run 2. N = 15 homozygous compared to 45 wild type. (**G**) Brain structure and activity maps for both runs, comparing heterozygous mutants to wild-type siblings. Run 1 N = 83 heterozygous compared to 36 wild type; Run 2 N = 91 heterozygous compared to 46 wild type.

As they displayed significant changes to brain structure and function, we expected these larvae to have behavioral phenotypes. Thus, we assessed them using a multi-day battery that included visual and acoustic stimulation (Figure 3A). Homozygous mutants displayed repeatable differences in baseline and stimulus-driven measures compared to control siblings (Figure 3, B and C, Supplementary Figure S2, A-D). While mutants tended to have fewer daytime bouts compared to wild-type larvae (Figure 3D, Supplementary Figure S2E), stronger phenotypes were observed in their location preference (Figure 3E, Supplementary Figure S2F) and structure or magnitude of these bouts (Figure 3F, Supplementary Figure S2H). They preferred the center of the well, and the individual movements had altered velocity, time, and displacement, which are all related measures. These animals had a marked increase in the number of revolutions they made in the bout (Figure 3F, Supplementary Figure S2G), a measure that is closely linked to their increased number of seizure-like movements (Figure 3H, Supplementary Figure S2I), quantified based on high-speed circling behavior (Baraban et al. 2005). When exposed to visual or acoustic stimuli, the mutants had reduced responses (Figure 3, I and J, Supplementary Figure S2, J and K). The magnitude of the dark flash response was particularly impacted.

**Figure 3.**
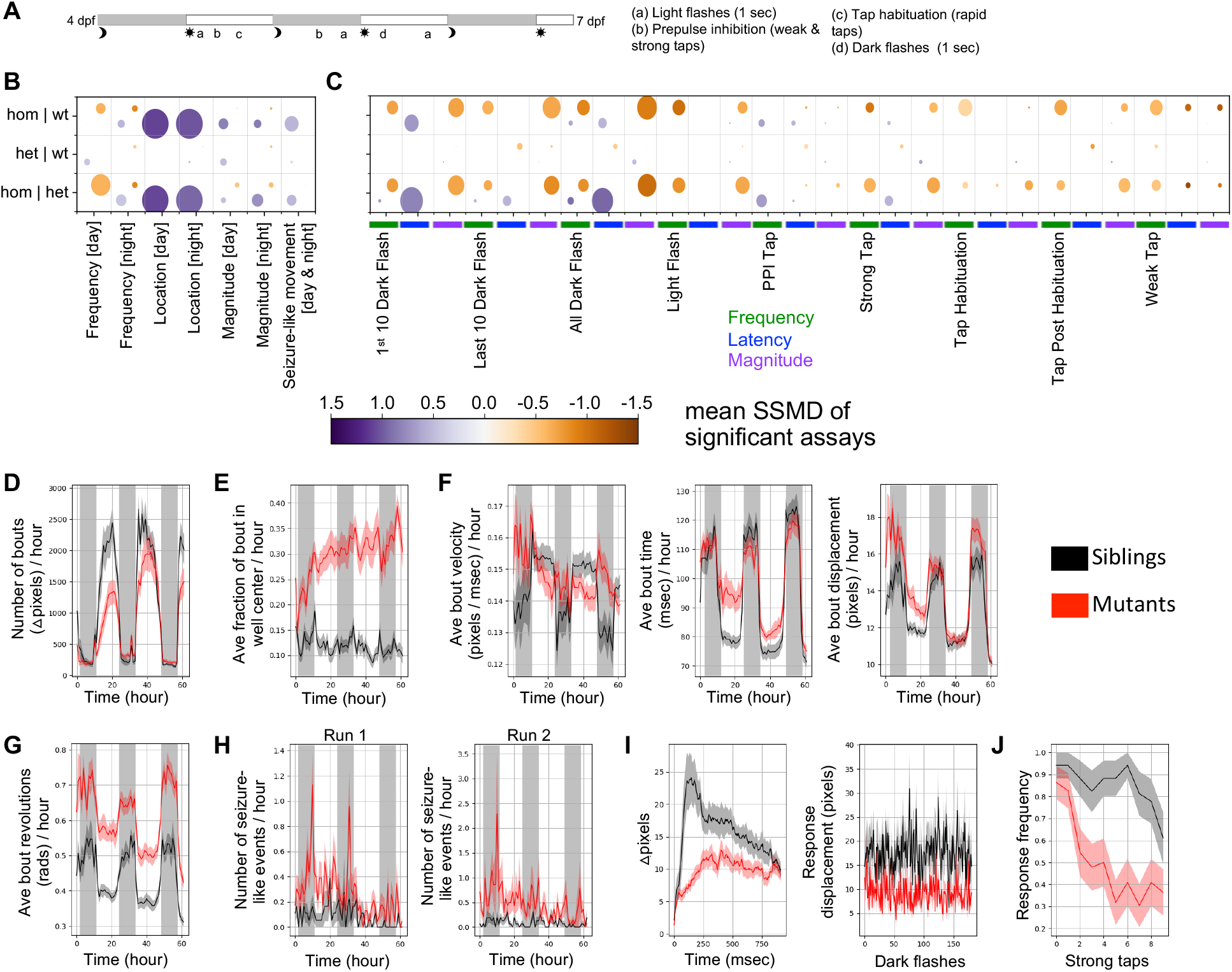
Behavioral phenotypes of zebrafish *ebf3a* mutants. (**A**) Multi-day behavioral pipeline with acoustic and visual stimulation. (**B**) Summary of baseline behavioral phenotypes in *ebf3a* mutants compared to siblings. The size of the bubble represents the percent of significant measurements in the summarized category, and the color represents the mean of the strictly standardized mean difference (SSMD) of the significant assays in that category. Run 1 N = 23 homozygous, 18 wild type, and 36 heterozygous. (**C**) Summary of stimulus-driven behavioral phenotypes in *ebf3a* mutants compared to siblings. (**D**) Example of a frequency of movement phenotype, shown for the entire duration of the experiment. This measure is the number of bouts, binned per hour, calculated using the change in pixels between each frame (linear mixed model p-value = 0.005). (**E**) Example of a location preference plot, shown for the direction of the experiment. This measure is the fraction of the frames of a bout that is spent in the center zone of the well, binned per hour, calculated based on the centroid positions of the fish (p-value = 0.001). Examples of bout magnitude measures, shown for the duration of the experiment, binned per hour, and calculated based on the centroid positions of the fish. The bout velocity is not significant when considering the experiment duration, but subregions do have significant p-values (e.g., the third night or day3nightall, binned per 10-minutes, p-value = 0.001). Bout time p-value for the experiment duration = 0.025, and bout displacement = 0.001. Bout revolutions for the duration of the experiment (p-value = 0.001). (**H**) Movements that resemble seizures (p-value = 0.001 for both sets). These are calculated based on the movement including more than 4 full revolutions, moving a distance of over 70 pixels, and having a speed of 0.3-1.3 pixels / msec. (**I**) Displacement for the dark flash response movement (right). Kruskal-Wallis ANOVA p-value = 1.7e-06. The response graph (right) is the average movement (pixel-based) of larvae in the mutant and control groups for events where a response was observed. (**J**) Frequency of responses to strong taps. The block shown is the strong taps completed prior to the first habituation block at 5 dpf (day5dpfhab1pre), with Kruskal-Wallis ANOVA p-value = 9.2e-05. Plots of mutant compared to control groups in panels D-J are mean ± s.e.m..

### Transcriptional differences in zebrafish *ebf3a* mutants indicate impacted development of multiple sensory systems and the cerebellum

To define the molecular basis for the observed neural phenotypes, we collected bulk RNA-sequencing data from *ebf3a* mutants at 2 dpf. This age was selected to uncover early molecular changes that could drive later phenotypes. Multiple differentially expressed genes were identified in all genotype comparisons (Figure 4A). As expected, *ebf3a* itself was significantly downregulated (Figure 4B). Many genes involved in eye development were upregulated (Figure 4B, Supplementary Figure S3A, Supplementary Data S1), which we grouped using Gene Ontology (GO) analysis. Other categories of dysregulated genes included lipid transport, cell-cell junction assembly, and vesicle organization. Surprisingly, no terms related to neuronal development emerged when the homozygous samples were compared to the combined data from heterozygous and wild-type siblings (Figure 4B), and only genes related to axon development were identified in the homozygous versus wild-type analysis (Supplementary Figure S3A). Among the genes with significantly reduced expression, however, were some with known involvement in brain development, such as *lhx1a* and *scn1lab* (Swanhart et al. 2010; Baraban et al. 2013; Lui et al. 2017; Symmank and Zimmer-Bensch 2019; Weinschutz Mendes et al. 2023). Among the most reduced genes in homozygous samples was *gpc1a*, the loss of which leads to reduced brain size in mice, possibly via reduced fibroblast growth factor (FGF) signaling (Jen et al. 2009). In zebrafish, *gpc1a* is expressed widely in the central nervous system, as well as also in the lateral line and some olfactory sensory neuron subtypes (Gupta and Brand 2013; Sur et al. 2023). Both the known Ebf3 target genes *pcdh8, neurod1*, and *prph* and the *ebf3* upstream repressor *arxa* were also differentially expressed (Fulp et al. 2008; Green and Vetter 2011) (Figure 4C). However, the target genes were unexpectedly upregulated in heterozygous and homozygous samples, opposite to the results from microinjection of mRNA and morpholinos in *Xenopus* (Green and Vetter 2011).

**Figure 4.**
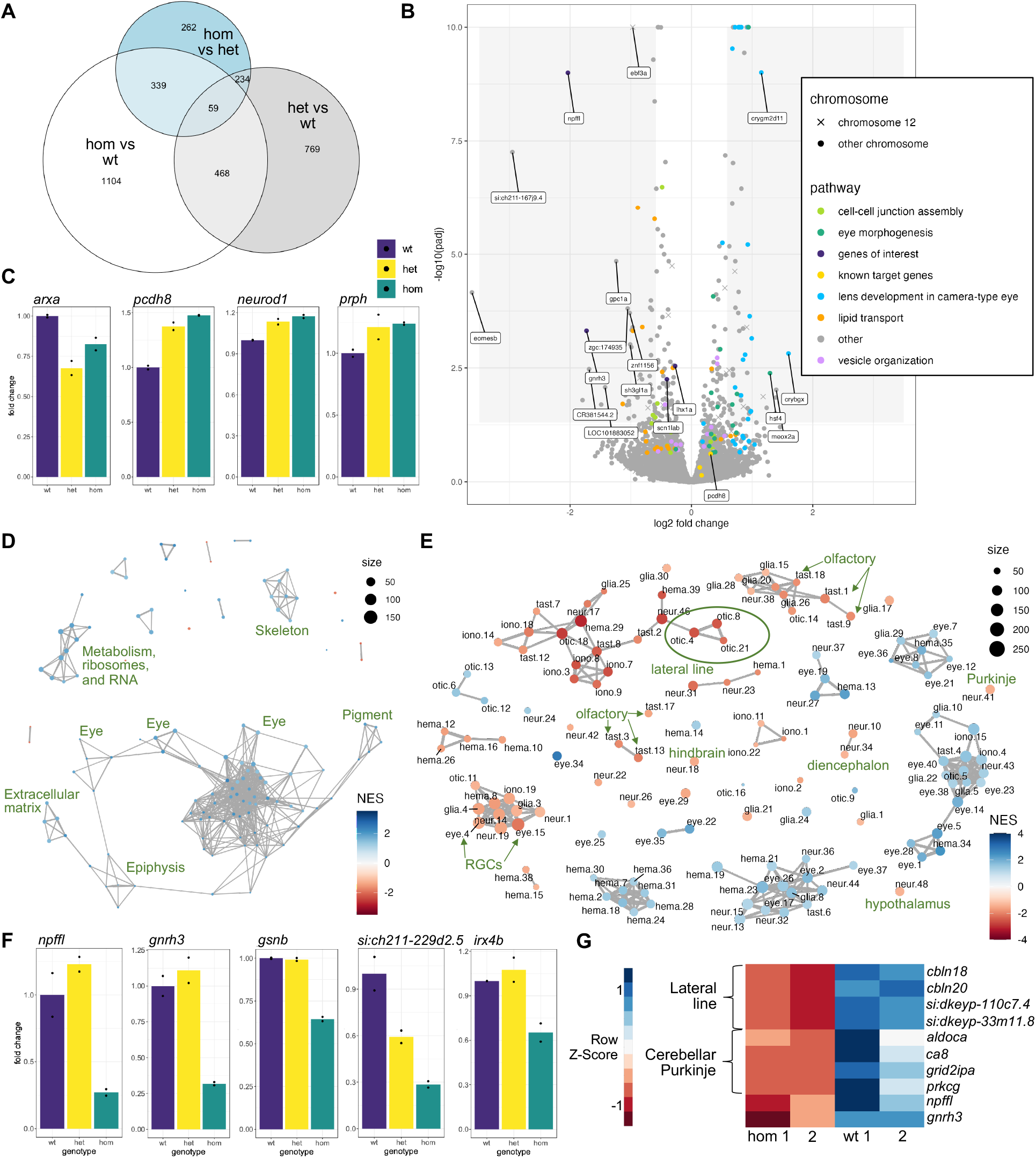
Transcriptional changes in zebrafish *ebf3a* mutants. (**A**) Shared DEGs between the genotype comparisons of 2 dpf RNA-sequencing data. The DEGs were filtered using a p-value of less than 0.05 and log2 fold change of greater than 0.2. (**B**) Volcano plot of the 2 dpf RNA-sequencing data for the comparison of homozygous mutants versus combined heterozygous and wild-type larvae. Genes involved in pathways identified by Gene Ontology (GO) analysis are identified. Genes with log2 fold changes of greater than 1.0 and adjusted p-values of less than 0.01 are labeled, as are several additional genes of interest. (**C**) Normalized counts data for wild-type, heterozygous, and homozygous samples for genes selected based on known involvement with *ebf3*. (**D**) Network plot of all GSEA C5 molecular signatures for the comparison of 2 dpf homozygous mutants versus combined heterozygous and wild-type larvae, with general groupings labeled. (**E**) GSEA network plot with single-cell types from the Daniocell larval zebrafish atlas. Clusters of particular interest are labeled. (**F**) Normalized counts data for wild-type, heterozygous, and homozygous samples for genes with strong downregulation in *ebf3a* mutants and mark specific neuron types. (**G**). Genes that are downregulated in 5 dpf RNA-sequencing data and mark specific neuron types. All analysis code, additional results, terms and genes found by the GO and GSEA analyses, and necessary files for the panels of this figure are available in Supplementary Data S1.

Using Gene Set Enrichment Analysis (GSEA), we highlighted additional pathways disrupted in the *ebf3a* mutants. For the homozygous mutants (Figure 4D, Supplementary Figure S3B), enrichment using the C5 set of molecular signatures overlapped with the GO analysis, with most terms resulting from the upregulation of eye genes. In the heterozygous samples compared to the wild type, however, many more genes involved in neuronal development, circadian behavior, posterior lateral line system development, channel activity, ion transport, and synapse assembly were uncovered by both GSEA and GO (Supplementary Figure S4, A and B). The differentially expressed genes in the heterozygous data overall had smaller fold changes than in the homozygous. This comparison, however, may reflect target genes of *ebf3a* better than the homozygous data that could be confounded by signatures of their altered development.

To predict impacted cell types, we also used GSEA to identify marker genes of cell types derived from scRNA-seq of the developing larvae (Sur et al. 2023). Genes selectively expressed in neuron types from multiple sensory systems were downregulated, including olfactory, visual (retinal ganglion cells), and lateral line subtypes (Figure 4E). Ebf3a is known to be expressed in retinal ganglion cells (RGCs), acting downstream of Atoh7 (Gao et al. 2014). The diencephalon, including the hypothalamus, and hindbrain, including the cerebellar Purkinje cells, emerged from this analysis. Highly specific marker genes of these nominated clusters were downregulated, such as olfactory sensory neurons (Supplementary Figure S5A). Two of the most differentially expressed genes, *npffl* and *gnrh3* (Figure 4F) are in so few cells that they are not obviously linked to specific neuron types in Daniocell, but their expression correlates with each other (Sur et al. 2023), and *gnrh3* is selectively expressed in cells near the olfactory placodes in larvae (Kurrasch et al. 2009; Bassi et al. 2020). Although the strongly reduced gene *gsnb* (Figure 4F) is expressed in multiple cell types, including the lateral line and olfactory system, other more specific markers such as *si:ch211-229d2*.*5* are expressed only in the lateral line and no other cell type (Supplementary Figure S5B), supporting the GSEA results (Figure 4E). Although cerebellar Purkinje cells are not detectable by Parvalbumin staining until 3 dpf (Kani et al. 2010), multiple transcriptional markers of this type are downregulated in our 2 dpf data, including *irx4b, ppargc1a*, and *rnf152* (Figure 4F, Supplementary Figure S5C). The transcription factor *ptf1a*, which marks the progenitors that give rise to Purkinje cells (Kani et al. 2010), was not differentially expressed (Supplementary Data S1). Most differentially expressed genes from RNA-seq collected at the later stage of 5 dpf were also highly specific markers of the lateral line and Purkinje cells (Figure 4G, Supplementary Figure S6), although the samples were less consistent, and no heterozygous were included. Despite the different developmental stage and lower quality of the 5 dpf data, multiple genes were shared between the two datasets (Supplementary Figure S6). Taken together, while molecular signatures of aberrant neuronal development were only prominent in the heterozygous samples, a comparison to scRNA-seq marker genes revealed that multiple sensory systems and specific brain areas likely have fewer or missing differentiated neurons in the homozygous larvae.

## Discussion

Here, we describe the characterization of an *ebf3a* zebrafish mutant. Loss of this *EBF3* ortholog results in substantial changes to brain structure, brain activity, and behavior, as well as lethality at late larval stages. Using GSEA in combination with single-cell marker genes highlighted that the lateral line and cerebellar Purkinje cells (Figure 4) are possibly missing or fewer in number. Sequencing data from both 2 and 5 dpf support this conclusion. Our results align well with one another, as the imaging (Figure 2) indicated a smaller cerebellum, and the altered movement structure (Figure 3) could reflect the impacted cerebellum and hindbrain. While our finding might have been expected, as Purkinje cell numbers are markedly decreased in mouse *Ebf2* null mutants (Croci et al. 2006) and *ebf3a* is expressed in this cell type, no specific data exists on orthologs of *EBF3* and cerebellar development. Our study also highlights other less obvious neuron types for future exploration, such as a hypothalamus subtype (neur.48) that is specific marked by *lmx1al* (Sur et al. 2023).

Some zebrafish phenotypes resemble those found in individuals with *EBF3* mutations. Ataxia and challenges with motor coordination are common. While it is not generally known how a larval zebrafish with ataxia would behave, we did observe changes in the bout structure (Figure 3F). Responses to stimuli were also dampened (Figure 3, I and J), possibly due to abnormal processing of the sensory input or motor coordination of the response. The cerebellum, particularly Purkinje cells, is strongly implicated in many ataxias (Huang and Verbeek 2019). A consistent finding in neuroimaging data is cerebellar hypoplasia (Harms et al. 2017; Tanaka et al. 2017; D’Arrigo et al. 2020), which we also observed in the homozygous mutants (Figure 2, B and F). The reduced response to dark flashes (Figure 3I) and dysregulation of genes involved in eye development (Figure 4, B and D) could arise similarly to ophthalmologic problems such as strabismus. The zebrafish larvae also displayed seizure-like movements, and abnormal EEG results and febrile seizures did occur in some individuals (Sleven et al. 2017).

One limitation of our model is that individuals with *EBF3* mutations are haploinsufficient. From our lethality data, we expect that a complete loss of *EBF3* would also not be compatible with human survival. The necessity of homozygosity to yield strong phenotypes in zebrafish with protein-truncating mutations in neurodevelopmental disorder genes is a common phenomenon (Hoffman et al. 2016; Zoodsma et al. 2022; Weinschutz Mendes et al. 2023; Capps et al. 2024 Jan 20). It is likely that genetic compensation mediated by the premature stop codon is responsible for the limited heterozygous phenotypes, as it could result in upregulation of the *ebf3a* wild-type allele or other transcription factors with a similar sequence that could act in its place (El-Brolosy et al. 2019). While the *ebf3a* level is reduced in the heterozygous samples (Supplementary Figure S4A), rather than upregulated as we have seen previously (Capps et al. 2024 Jan 20), its change is far less than then expected 50% (Supplementary Data S1: −0.14 log2 fold change, p-value = 0.016). We did not observe an upregulation of *ebf3b* or *ebf2*, but *ebf1a* and *ebf1b* were slightly upregulated. In the future, making a new mutant line with the promoter removed, which would not produce truncated RNAs that drive this compensation, could yield stronger heterozygous phenotypes. It is also possible that this compensation could come through genetic mechanisms other than nonsense-mediated decay. The *arxa* gene is downregulated in both heterozygous and homozygous samples (Figure 4C), and *Ebf3* was shown to be a direct target of *Arx* in the developing mouse brain (Fulp et al. 2008). When *Arxa* is reduced, *Ebf3* is upregulated, and it is possible that *arxa* is also a target of *ebf3a* and is itself downregulated to derepress *ebf3a*.

Although the heterozygous mutants had very mild phenotypes (Figure 2G), their transcriptional dysregulation was extensive (Supplementary Figure S4). The levels of target genes known from the literature (Figure 4C) are as strongly increased in heterozygous and homozygous, and the *arxa* was downregulated in both. While the expectation is that the level of target genes would be reduced in correspondence to lower *ebf3a*, those experiments were done using transient microinjection in *Xenopus* (Green and Vetter 2011), which could have a different outcome than for our germline mutant. Pathway analysis uncovered gene sets involved in brain structure and function, which was not the case for homozygous mutants. Sets of interest included circadian behavior, as individuals with *EBF3* mutations have disturbed sleep, and multiple ion channel subunits. While it is surprising that there is significant transcriptional dysregulation with mild observed neural phenotypes (Figure 2G), these gene sets could more directly reflect the transcriptional response to reducing *ebf3a*, rather than the emergent morphological changes in mutants. There are likely missing or disproportionately skewed cell populations in the homozygous mutants and not in the heterozygous siblings. A goal of our work was to uncover downstream target genes of *ebf3a* that could be responsible for the mutant neural phenotypes, and those dysregulated in heterozygous samples and more so in homozygous could be promising for further study.

Several strongly dysregulated genes are possible drivers of the observed phenotypes. Zebrafish mutants in *scn1lab*, which is downregulated in *ebf3a* homozygous mutants, have reduced size in the midbrain-hindbrain boundary region (Weinschutz Mendes et al. 2023). This region overlaps with the area that is ventral to the cerebellum and smaller in *ebf3a* mutants (Figure 2B). Brain activity differs between the two, however, as *scn1ab* has reduced activity throughout the entire brain compared to the strong activity increase in the cerebellum of *ebf3a* mutants. Behaviorally, both display seizure-like movements (Baraban et al. 2013) (Figure 3H), as do *arxa* mutants (Griffin et al. 2021). In the Purkinje cell cluster, *irx4b* is expressed before other marker genes (Supplementary Figure S5C), beginning at 24-34 hpf in Daniocell when the cells are just beginning to be classified. This transcription factor is a possible early driver of Purkinje cell differentiation and has been linked to axonal pathfinding in retina (Jin et al. 2003). The progenitors that give rise to Purkinje cells (Kani et al. 2010) do not appear affected, as *ptf1a* is not differentially expressed. The genes *skor1a* and *skor1b*, which are key regulators of the first step in Purkinje cell differentiation from *ptf1a* progenitors, are also not differentially expressed. Collecting RNA-sequencing data from an earlier stage, such as 36 hpf, could support this hypothesis and reveal other early *ebf3a* target genes.

The phenotypes of *ebf3a* homozygous mutants could be a starting point for drug discovery. Transgenic lines that specifically label the lateral line and Purkinje cells (Tanabe et al. 2010; Fuentes et al. 2016) could be used for an imaging-based drug screen (Wang et al. 2015). Seizure-like movements in zebrafish larvae were used as a screening platform to find a new treatment for Dravet syndrome that is in clinical trials (Baraban et al. 2013; Dinday and Baraban 2015).

Overall, our findings provide a foundation for understanding the role of *ebf3a* in neurodevelopment and offer a promising avenue for developing targeted therapeutic interventions.

## Materials and Methods

### Zebrafish husbandry and mutant generation

Zebrafish (*Danio rerio*) were maintained in the Zebrafish Research Facility at the University of Alabama at Birmingham at 28°C on a 14-hours light / 10-hours dark cycle. All animal studies were performed according to the guidelines by the Institutional Animal Care and Use Committee of the University of Alabama at Birmingham (IACUC protocols 22279, 22155, and 21744).

*ebf3a* and *ebf3b* mutants were generated by microinjection of Cas9 protein and one guide RNA (gRNA) into AB wild-type zebrafish embryos as described previously (Wang et al. 2021). The target site, PAM motif, and identified out of frame mutant are listed in figures 1 and S1.

Experiments were completed using larvae derived from crosses using the F2 generation and later. To genotype mutants, genomic DNA was extracted from whole zebrafish embryos or tail clip from adult zebrafish using alkaline lysis. Then, High-Resolution Melting (HRM) analysis was performed (Thomas et al. 2014): PCR with gene-specific primers (Supplementary Table S2) was completed in black/white 96 well plates (BioRad cat. No. HSP9665) and followed by the generation of melting curves using a Lightscanner HR 96 (Idaho Technology). All experiments were conducted blind, and genotyping occurred after the data was collected.

#### RNA extraction, Real time quantitative PCR (RT-qPCR), and RNA-seq

For RT-qPCR, total RNA was extracted from the anterior half of the body from 10-12 5 dpf larvae per sample at using the Rneasy Mini Kit (Qiagen). cDNA was synthesized using the High-Capacity cDNA Reverse Transcription Kit (Applied Biosystems). The RT-qPCR was performed using SsoAdvanced SYBR Green Supermix (Bio-Rad), primers (Supplementary Table S1), and CFX96 Touch Deep Well Real-Time PCR Detection System (Bio-Rad). RNA for the 2 dpf

RNA-seq samples were collected in the same way, but with 15-20 embryos per pool. The 5 dpf RNA-seq was collected also with 10-12 larvae. The posterior portion of the cuts was used for genotyping. The RNA was isolated from the anterior portion as described and submitted to sequencing by GENEWIZ at Azenta Life Sciences.

#### RNA-sequencing analysis

Paired-end sequencing reads were aligned using STAR aligner (2.7.10a) (Dobin et al. 2013) to the GRCz11 release 104 using the Lawson Lab Zebrafish Transcriptome Annotation version 4.3.2 (Lawson et al. 2020). Using DESeq2 (Love et al. 2014), the raw counts files were normalized using rlog counts method. An example DESeq2 RScript and input files are available at https://github.com/thymelab/BulkRNASeq. The fold-change tables, all downstream analysis code, and complete results from transcriptome analyses are available in Supplementary Data S1 on Zenodo.

Gene Ontology (GO) analysis and Gene Set Enrichment Analysis (GSEA) (Subramanian et al. 2005) were performed using the clusterProfiler_4.10.0 R package, using the enrichGO and GSEA functions, respectively. To perform GSEA using the Daniocell single-cell data (Sur et al. 2023), the single-cell object was downloaded from the Daniocell website. Markers of the annotated clusters were identified using FindAllMarkers from Seurat_5.0.0 package (Satija et al. 2015), and the top 500 markers were used as gene sets for GSEA.

#### Brain activity and morphology

Phosphorylated-ERK (pErk) antibody staining was conducted and analyzed as previously described (Thyme et al. 2019; Capps et al. 2024 Jan 20). Larvae were kept in quiet conditions for a minimum of a half hour before rapid fixation. They were fixed overnight at 4°C in 4% paraformaldehyde (PFA) (Polysciences) diluted in 1X Phosphate Buffered Saline, permeabilized with 0.05% Trypsin-EDTA on ice for 30-35 minutes, stained for 2-3 days with the total ERK antibody (Cell Signaling, #4696) and phospho-Erk antibody (Cell Signaling #4370), and 1-2 days with Alexa Fluor 647 mouse (A21235) and 488 rabbit (A11008) secondary antibodies (Thermo Fisher Scientific). After imaging with a Zeiss 900 upright confocal with 20x 1.0 NA water-dipping objective, larvae were removed from agarose for genotyping. As previously, the confocal stacks were registered to the Z-Brain atlas and processed using MapMAPPING with a false discovery rate (FDR) of 0.05% (Randlett et al. 2015; Capps et al. 2024 Jan 20).

#### Larval behavior

Larval behavior assays were completed using custom-built behavioral systems as previously described (Thyme et al. 2019; Joo et al. 2020). Briefly, on the evening of 4 dpf, the animals are placed in 96-well plates with square wells and sealed with oxygen permeable film. The white light cycle is 9 AM on / 11 PM off. The larvae are exposed to the following stimuli at 5 dpf: light flashes (9:11-9:25), mixed acoustic stimuli (prepulse, strong, and weak, from 9:38-2:59), three blocks of acoustic habituation (3:35-6:35). The night between 5 and 6 dpf: mixed acoustic stimuli (1:02-5:00), light flashes (6:01-6:20). During the day at 6 dpf: three blocks of dark flashes with an hour between them (10:00-3:00), mixed acoustic stimuli and light flashes (4:02-6:00). The baseline framerate is 30 frames-per-second, while the responses to stimuli are analyzed from high-speed (285 frames-per-second) videos of a 1-second duration. All code for behavior analysis is available at https://github.com/thymelab/ZebrafishBehavior. As previously, the significance of data from baseline blocks, such as the duration of the experiment, are calculated with a linear mixed model (Thyme et al. 2019; Joo et al. 2020), while stimulus response data is assessed using the more standard Kruskal-Wallis ANOVA. The ANOVA values for these longer durations, with large variation in the measures across time, are also available in the supplementary files on Zenodo. To generate the bubble plot summary visualization, related measures are combined. The size of the bubble represents the percent of significant measurements in the summarized category, and the color represents the mean of the strictly standardized mean difference (SSMD) of the significant assays in that category. The bubbles are offset and both included (versus a typical heatmap), because both decreased and increased measures can occur in the same summarized block, such as if a change occurs between the 4 and 6 dpf stages. The scripts for merging these measurements and generating the summarized bubble plot graphs are available at https://github.com/thymelab/DownstreamAnalysis.

## Acknowledgments

We thank the UAB fish facility staff, the UAB Department of Pharmacology and Toxicology, the UAB Department of Neurobiology, and the Research Computing team at UAB for supporting this study.

## Author contributions

NCY, SBT, and JMP conceived of the study. NCY and SBT designed the experiments. CLC and MDV collected the behavior and imaging data, with genotyping assistance from NDPD. NDPD, TQT, and JW did the morphology and lethality assessment. NDPD and TQT collected the RT-qPCR and RNA-sequencing data. HRT and JMP generated the *ebf3a* and *ebf3b* mutant lines. SBT and NCY analyzed the data. SBT wrote the manuscript with contributions from NDPD and NCY and feedback from JMP.

## Data availability

All data are available in the main text, the Supplementary Materials, or appropriate databases. Code is available in Supplementary Data S1 or from https://github.com/thymelab. Supplementary Data S1 and processed behavioral and imaging data are available from Zenodo (10.5281/zenodo.13738007). Additional RNA-seq analysis results are in Supplementary Data S1 on Zenodo. Raw behavioral and imaging files that were too large for Zenodo are available upon request. RNA-sequencing data is available from GEO (GSE276705).

