## Supplementary Material for "Disrupted development of sensory systems and the cerebellum in a zebrafish *ebf3a* mutant"

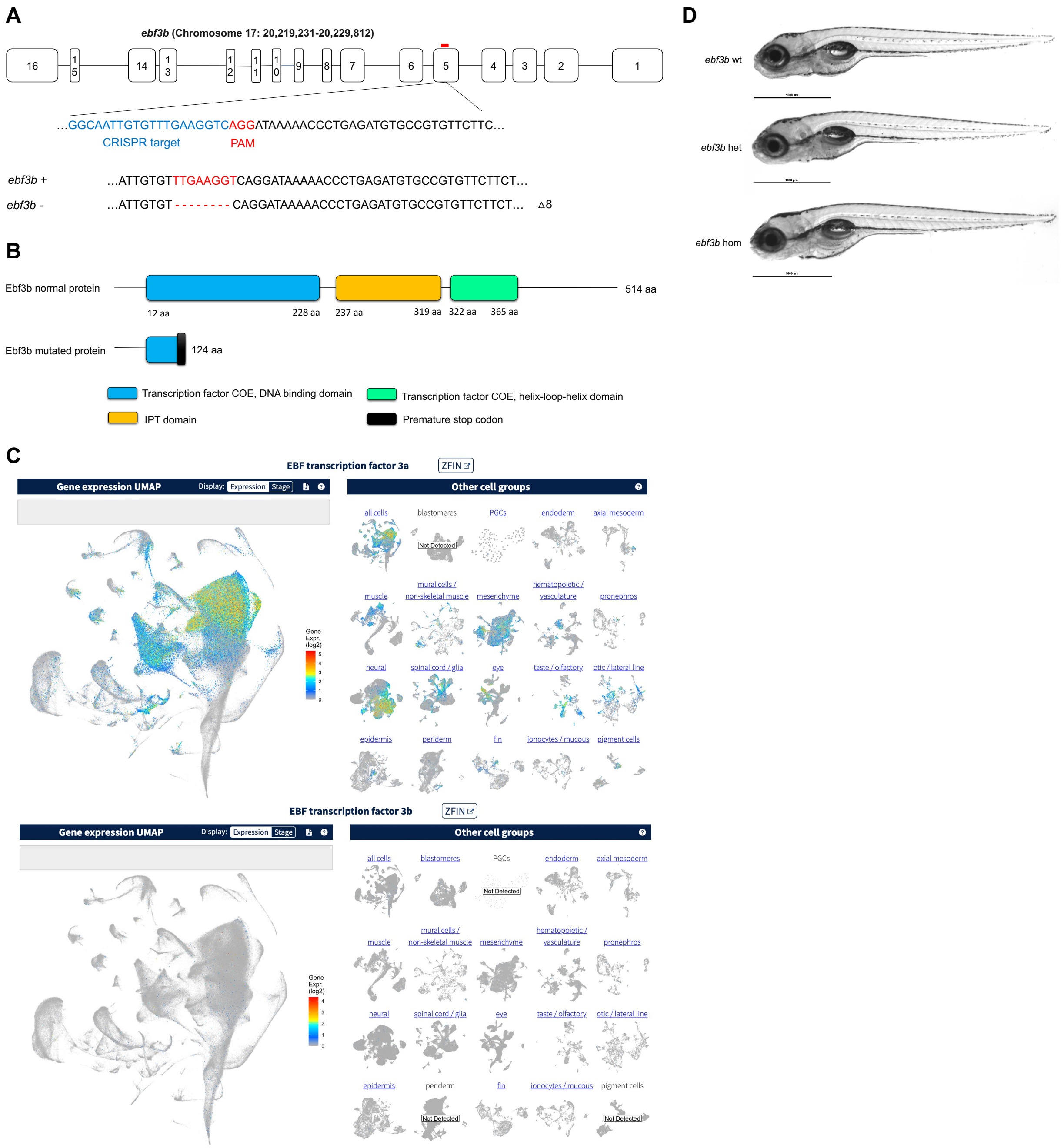


**Figure S1. Generation and developmental characterization of zebrafish *ebf3b* mutants.** (**A**) Schematic of zebrafish *ebf3b* gene transcript and CRISPR gRNA targeting sequence. (**B**) Schematic of zebrafish wild-type and mutant Ebf3b proteins. Domains are annotated based on ensemble Pfam database. (**C**) Expression of *ebf3a* and *ebf3b* in single-cell RNA-sequence data from Daniocell (https://daniocell.nichd.nih.gov/) (Sur et al. 2023). (**D**) Representative photos of *ebf3b* homozygous mutants and respective siblings at 5 dpf.

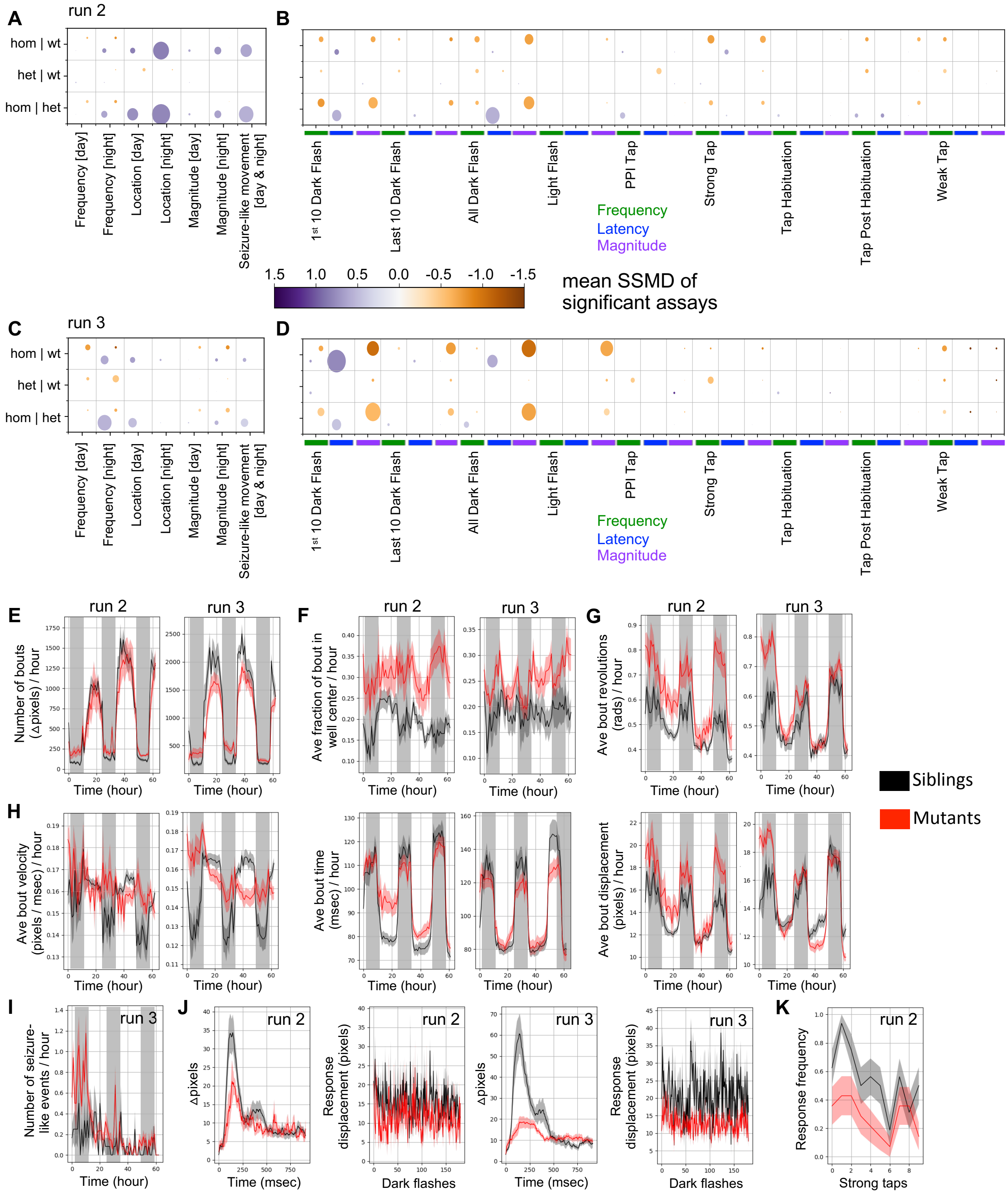


**Figure S2. Behavioral data from two additional independent clutches.** (**A**) Summary of baseline behavioral phenotypes for second run. Run 2 N = 14 homozygous, 16 wild type, and 37 heterozygous. (**B**) Summary of stimulus-driven behavioral phenotypes for second run. (**C**) Summary of baseline behavioral phenotypes for third run. Run 3 N = 34 homozygous, 12 wild type, and 36 heterozygous. (**D**) Summary of stimulus-driven behavioral phenotypes for third run. (**E**) P-values from the entire experiment duration are non-significant for both runs. However, some subsection p-values are significant, such as the daytime of day 3 (day3msdf, binned per 10-

minutes, p-value = 0.001 for both runs). Both these runs indicate there may be an increase in nighttime movement as well as the decrease in daytime movement, suggesting possibly disrupted sleep. (**F**) P-values for the entire experiment duration for run 2 = 0.009. For run 3, however, it was non-significant on the entire experiment duration. The p-value for subsections were significant, such as the morning of day 2 (day2morning, binned per 10-minutes, p-value = 0.002). (**G**) P-values for both runs = 0.001. (**H**) For run 2, the bout velocity for the experiment duration was not significant, but subsections were (day3nightall, binned per 10-minutes, p-value = 0.001). Run 3 p-value for bout velocity over the experiment duration = 0.005. The bout time p-value for run 2 = 0.03 and is non-significant for run 3. The bout displacement p-value = 0.001 for both runs. (**I**) P-value for run 3 = 0.001 (run 2 is in the corresponding main text figure). Displacement Kruskal-Wallis ANOVA p-values = 0.0028 (run 2) and 2.9e-05 (run 3). (**J**) Dark flash response displacement plots (left in pair) and response graphs (right in pair). (**K**) Frequency of responses to strong taps. The block shown is the strong taps completed prior to the first habituation block at 5 dpf (day5dpfhab1pre), with Kruskal-Wallis ANOVA p-value = 0.0067 (run 2). The same section of taps was not significant for run 3, and strong tap responses were generally least affected in this clutch.

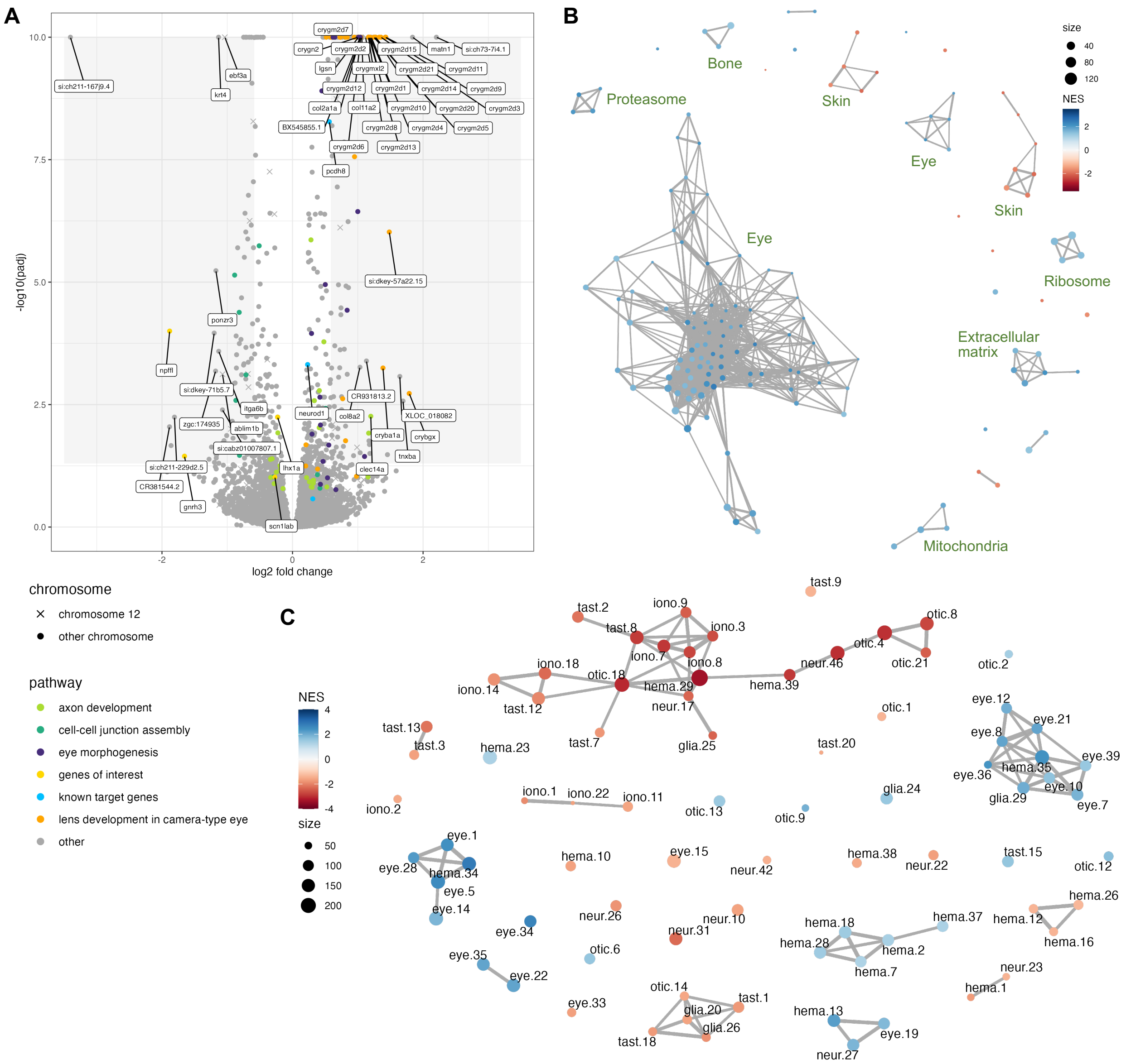


**Figure S3. Analysis of *ebf3a* homozygous mutants versus wild-type siblings 2 dpf RNA-sequencing data.** (**A**) Volcano plot of the 2 dpf RNA-sequencing data for the comparison of homozygous mutants versus wild-type larvae. Genes involved in pathways identified by Gene Ontology (GO) analysis are identified, as are several additional genes of interest. Genes with log2 fold changes of greater than 1.0 and adjusted p-values of less than 0.01 are labeled. (**B**) Network plot of all GSEA C5 molecular signatures, with general groupings labeled. (**C**) GSEA network plot with single-cell types from the Daniocell larval zebrafish atlas.

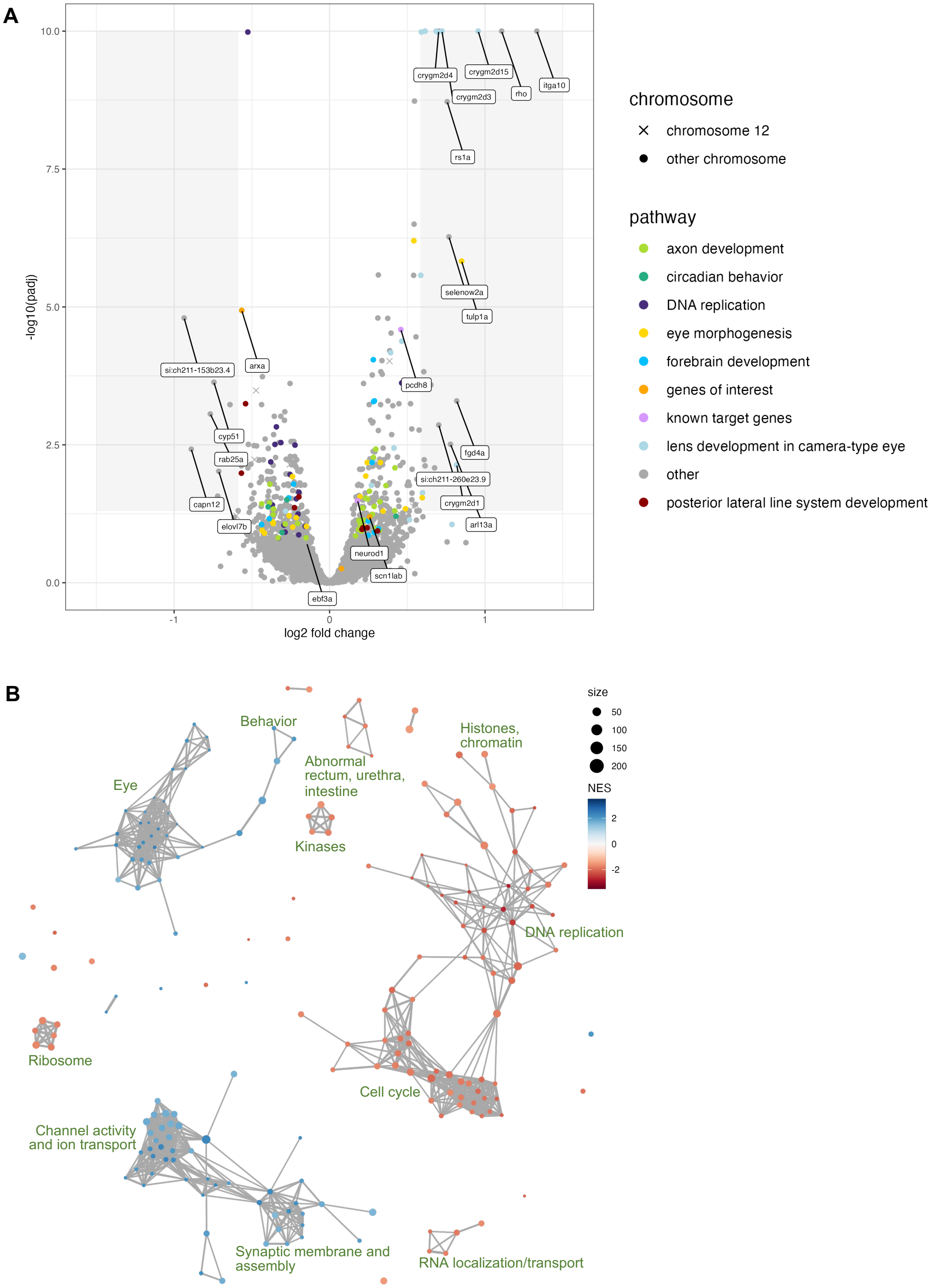


**Figure S4. Analysis of *ebf3a* heterozygous mutants versus wild-type siblings 2 dpf RNA-sequencing data.** (**A**) Volcano plot of the 2 dpf RNA-sequencing data for the comparison of heterozygous mutants versus wild-type larvae. Genes involved in pathways identified by Gene Ontology (GO) analysis are identified, as are several additional genes of interest. Genes with log2 fold changes of greater than 0.7 and adjusted p-values of less than 0.01 are also labeled. (**B**) Network plot of all GSEA C5 molecular signatures, with general groupings labeled.

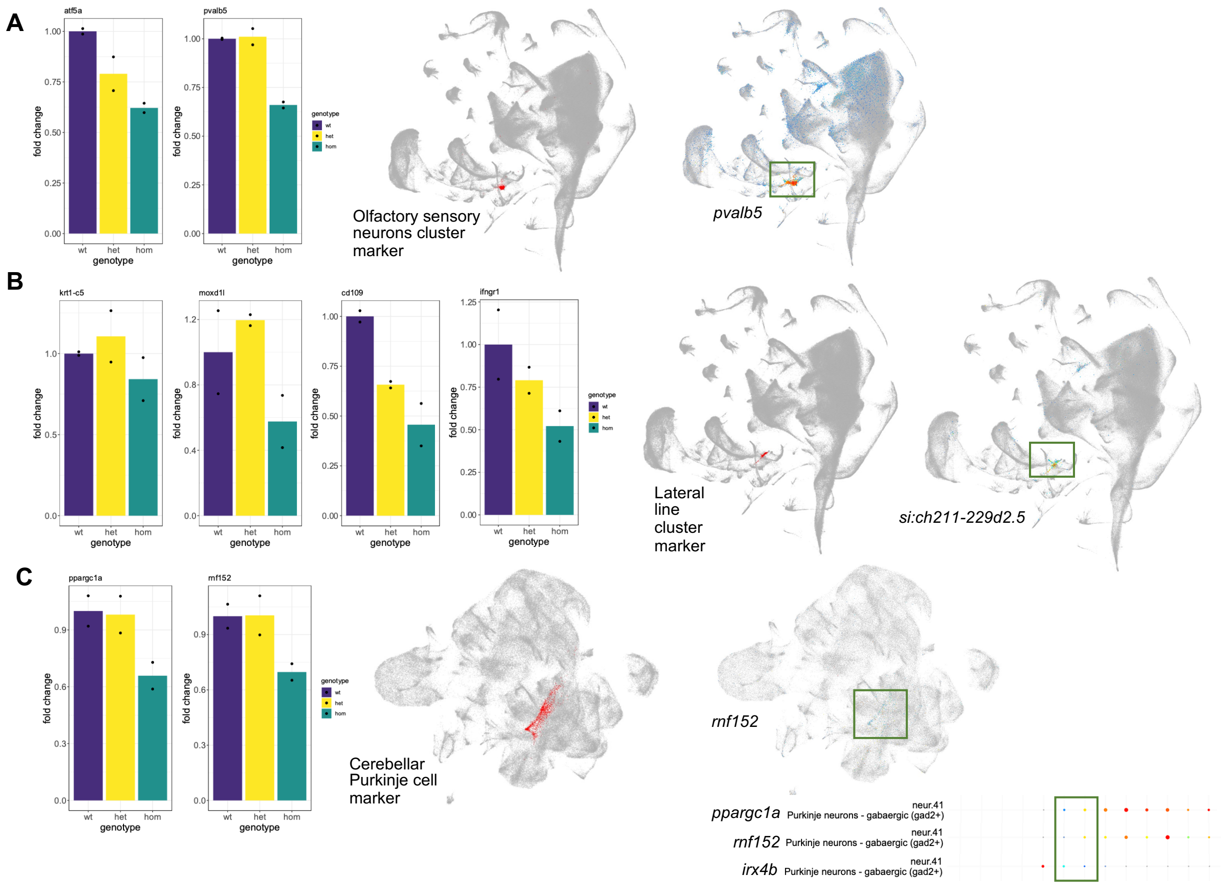


**Figure S5. Examples of downregulated genes that mark specific single-cell clusters in Daniocell.** (**A**) The bar plots are normalized counts data for wild-type, heterozygous, and homozygous samples for genes selected as downregulated and marking olfactory sensory neurons. The Daniocell single-cell maps on the left are the cells associated with the cluster in the whole larval dataset from 0-5 dpf (left) and expression of one marker gene (right). (**B**) Plots are equivalent to **A** but for the lateral line markers. (C) ) Plots are equivalent to **A** but for the lateral cerebellar Purkinje cells. Additionally, the dot plot is from Daniocell and the box represents the marker expression in this cell type most similar to when the samples were collected (24-26 hpf and 48-58 hpf).

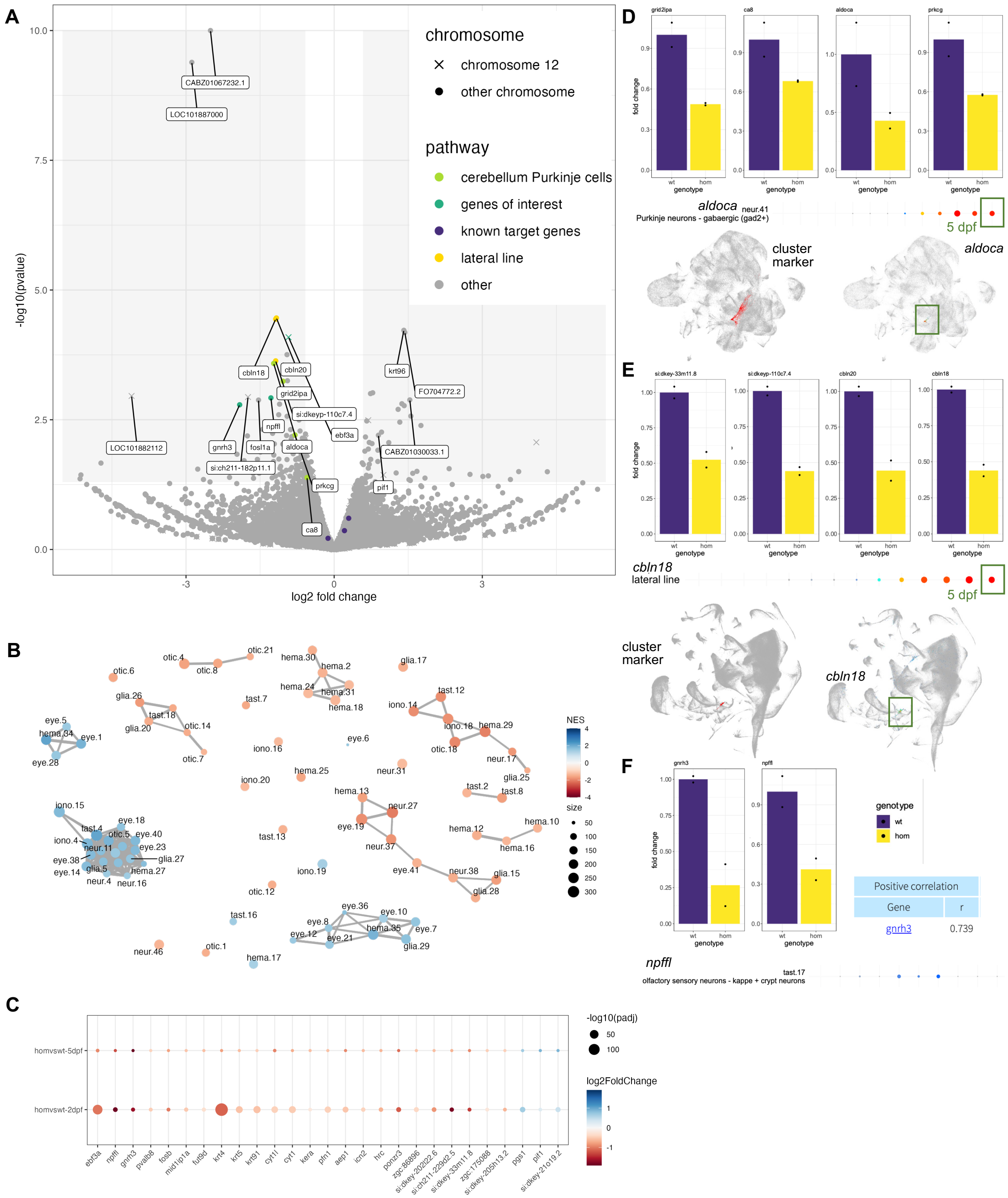


**Figure S6. Analysis of RNA-sequencing data from 5 dpf homozygous *ebf3a* mutants compared to wild type siblings.** (**A**) Volcano plot with labels for genes of interest that specific neuron types. Genes with log2 fold changes of greater than 1.0 and p-values of less than 0.0015 are also labeled. (**B**) GSEA network plot with single-cell types from the Daniocell larval zebrafish atlas. (**C**) Bubble plot of differentially expressed genes shared between the 2 dpf and 5 dpf data. (**D**) The bar plots are normalized counts data for wild-type and homozygous samples for genes selected as downregulated and marking cerebellar Purkinje cells. The Daniocell single-cell maps on the left are the cells associated with the cluster in the whole larval dataset from 0-5 dpf (left) and expression of one marker gene (right). (**E**) Plots are equivalent to **D** but for the lateral line markers. Additionally, the dot plot is from Daniocell and the box represents the marker expression in this cell type most similar to when the samples were collected (5 dpf). (**F**) Plots are equivalent to **D** but for the two very downregulated genes that likely mark a population of olfactory sensory neurons. The two genes correlate with each other.

**Table S1. Survival of *ebf3a* mutants over development.** The asterisk (*) represents clutches where *ebf3b* was also homozygous mutant as the background genotype.

| **Age** | **Total genotyped** | **Wild type** | **Heterozygous** | **Homozygous** |
|---|---|---|---|---|
| 5 dpf | 89 | 17 | 47 | 25 |
| 7 dpf* | 34 | 3 | 21 | 10 |
| 7 dpf* | 41 | 13 | 16 | 12 |
| 7 dpf* | 47 | 10 | 19 | 18 |
| 7 dpf* | 47 | 6 | 28 | 13 |
| 8 dpf | 65 | 18 | 33 | 14 |
| 10 dpf* | 89 | 37 | 47 | 5 |
| 10 dpf* | 84 | 29 | 41 | 14 |
| 11 dpf | 28 | 12 | 12 | 4 |
| 11 dpf | 41 | 13 | 18 | 11 |
| 15 dpf | 9 | 5 | 4 | 0 |
| 15 dpf | 43 | 22 | 21 | 0 |
| 15 dpf | 23 | 14 | 9 | 0 |
| 6 wpf | 18 | 9 | 9 | 0 |
| 2 mpf | 19 | 8 | 11 | 0 |
| 3.5 mpf | 23 | 6 | 17 | 0 |

**Table S2. Primers for genotyping and RT-qPCR.**

| | **Sequence (5′-3′)** |
|---|---|
| *ebf3a* HRM | Forward: CTGGGCAGTGGCATGAAT |
| | Reverse: CGGATGATTTGGCAAACGTA |
| *ebf3b* HRM | Forward: TTGCTTTACATAACAATCTGCTGTT |
| | Reverse: TCATGCGTGAGAAGAACACG |
| *ebf3a* RT-qPCR | Forward: ACAGTCAATGTGGACGGTCA |
| | Reverse: GAAGTTGTCGCCAATGATGA |
| *ebf3b* RT-qPCR | Forward: AGGAAACCCACGAGACACAC |
| | Reverse: CCTTAATGCAGGGGATCTCA |
| *actb1* RT-qPCR (housekeeping) | Forward: CATCCGTAAGGACCTGTATGCCAAC |
| | Reverse: AGGTTGGTCGTTCGTTTGAATCTC |
