## Supplementary Data S1 for "Disrupted development of sensory systems and the cerebellum in a zebrafish *ebf3a* mutant": ebf3-rnaseq-analysis.html

ebf3a bulk RNA-seq analysis


Code 

- Show All Code
- Hide All Code

### ebf3a bulk RNA-seq analysis

###### Summer Thyme, Anna Moyer

#### *2024-08-15*

Make plots for RNAseq data collected for ebf3a gene.

### 1 Setup zone

#### 1.1 Set seed and working directory

#### 1.2 Load packages

```
library("tidyverse") 
library("cowplot") #arranging plots into grids
library("viridis") #viridis color schemes
library("scales") #use to get color schemes for viridis
library("ggrepel") #repels text labels on plots
library("RColorBrewer") #pick colors
library("DT") #interactive and searchable tables of our GSEA results
library("GSEABase") #functions and methods for Gene Set Enrichment Analysis
library("Biobase") #base functions for bioconductor; required by GSEABase
library("GSVA") #Gene Set Variation Analysis, a non-parametric and unsupervised method for estimating variation of gene set enrichment across samples.
library("gprofiler2") #tools for accessing the GO enrichment results using g:Profiler web resources
library("clusterProfiler") # provides a suite of tools for functional enrichment analysis
library("dplyr")
library("readr")
library("stringr")
library("org.Dr.eg.db")  # For zebrafish; change if needed
library("msigdbr") # access to msigdb collections directly within R
library("enrichplot") # great for making the standard GSEA enrichment plots
library("ontologyIndex") #for parsing obo files
library("BaseSet") #for importing gaf file
library("plotly") #make interactive plots
library("lattice") #used for making manhattan plot
library("ggpubr") #calculating correlation coefficient 
library("colorspace") #colors for heatmap
library("ggraph") #library for making network graphs
library("svglite") #export svgs in loop
library("gplots") #the heatmap2 function in this package is a primary tool for making heatmaps
library("venn")
library("eulerr")
```

### 2 Function zone

#### 2.1 Add chromosome information to DEG csv

```
process_DEGs <- function(input_csv, annotation_tsv, output_prefix, chrnum) {
  # Import data frame with the previously calculated DEGs
  DEG <- read.csv(input_csv)[,2:9]
  # Limit padj to 1e-10
  DEG <- DEG %>% 
    mutate(padjbound = ifelse(padj < 1e-10, 1e-10, padj))
  # Limit pvalue to 1e-10
  DEG <- DEG %>% 
    mutate(pvaluebound = ifelse(pvalue < 1e-10, 1e-10, pvalue))
  # Import annotations for chromosome and TSS
  chromosomeinfo <- readr::read_tsv(file = annotation_tsv)
  # Keep chromosome and gene name
  chromosomeinfo <- chromosomeinfo %>% select(chrom, name2, txStart)
  # Make empty columns to hold chromosome and txStart
  DEG$chrom <- NA
  DEG$txStart <- NA
  # Add chromosome and txStart information to data frame
  for (x in 1:nrow(DEG)) {
    gene <- DEG$LLgeneAbbrev[x]
    chromosomeinfosub <- chromosomeinfo %>% filter(name2 == gene)
    if (nrow(chromosomeinfosub) > 0) {
      chr <- chromosomeinfosub$chrom[1]
      txStart <- chromosomeinfosub$txStart[1]
      DEG$chrom[x] <- chr
      DEG$txStart[x] <- txStart
    }
  }
  # Add a column with chromosome number
  # FUTURE: this would change or need to be more flexible if using a different species with different numbers of chromosomes
  DEG <- DEG %>% 
    mutate(chromosomenumber = str_replace_all(chrom, c(
      "chr1" = "1", "chr2" = "2", "chr3" = "3", "chr4" = "4", 
      "chr5" = "5", "chr6" = "6", "chr7" = "7", "chr8" = "8", 
      "chr9" = "9", "chr10" = "10", "chr11" = "11", "chr12" = "12", 
      "chr13" = "13", "chr14" = "14", "chr15" = "15", "chr16" = "16", 
      "chr17" = "17", "chr18" = "18", "chr19" = "19", "chr20" = "20", 
      "chr21" = "21", "chr22" = "22", "chr23" = "23", "chr24" = "24", 
      "chr25" = "25"
    )))
  # Remove duplicate rows
  DEG <- distinct(DEG)
  # Export CSV
  write.csv(DEG, file = paste0(output_prefix,"_DEG_chr.csv"), row.names = FALSE)
  return(DEG)
}
```

#### 2.2 Determining relationship of DEGs with mutant chromosome

##### 2.2.1 Plotting DEGs by chromosome with Manhattan plot

Are differentially expressed genes located at a particular place in
the genome? The manhattan plot function is from: https://genome.sph.umich.edu/wiki/Code\_Sample:\_Generating\_Manhattan\_Plots\_in\_R

```
#function for making manhattan plots
manhattan_plot_graph<-function(chr, pos, pvalue, 
    sig.level=NA, annotate=NULL, ann.default=list(),
    should.thin=T, thin.pos.places=2, thin.logp.places=2, 
    xlab="Chromosome", ylab=expression(-log[10](p-value)),
    col=c("gray","darkgray"), panel.extra=NULL, pch=20, cex=0.8,...) {
    if (length(chr)==0) stop("chromosome vector is empty")
    if (length(pos)==0) stop("position vector is empty")
    if (length(pvalue)==0) stop("pvalue vector is empty")
    #make sure we have an ordered factor
    if(!is.ordered(chr)) {
        chr <- ordered(chr)
    } else {
        chr <- chr[,drop=T]
    }
    #make sure positions are in kbp
    if (any(pos>1e6)) pos<-pos/1e6;
    #calculate absolute genomic position
    #from relative chromosomal positions
    posmin <- tapply(pos,chr, min);
    posmax <- tapply(pos,chr, max);
    posshift <- head(c(0,cumsum(posmax)),-1);
    names(posshift) <- levels(chr)
    genpos <- pos + posshift[chr];
    getGenPos<-function(cchr, cpos) {
        p<-posshift[as.character(cchr)]+cpos
        return(p)
    }
    #parse annotations
    grp <- NULL
    ann.settings <- list()
    label.default<-list(x="peak",y="peak",adj=NULL, pos=3, offset=0.5, 
        col=NULL, fontface=NULL, fontsize=NULL, show=F)
    parse.label<-function(rawval, groupname) {
        r<-list(text=groupname)
        if(is.logical(rawval)) {
            if(!rawval) {r$show <- F}
        } else if (is.character(rawval) || is.expression(rawval)) {
            if(nchar(rawval)>=1) {
                r$text <- rawval
            }
        } else if (is.list(rawval)) {
            r <- modifyList(r, rawval)
        }
        return(r)
    }
    if(!is.null(annotate)) {
        if (is.list(annotate)) {
            grp <- annotate[[1]]
        } else {
            grp <- annotate
        } 
        if (!is.factor(grp)) {
            grp <- factor(grp)
        }
    } else {
        grp <- factor(rep(1, times=length(pvalue)))
    }
    ann.settings<-vector("list", length(levels(grp)))
    ann.settings[[1]]<-list(pch=pch, col=col, cex=cex, fill=col, label=label.default)
    if (length(ann.settings)>1) { 
        lcols<-trellis.par.get("superpose.symbol")$col 
        lfills<-trellis.par.get("superpose.symbol")$fill
        for(i in 2:length(levels(grp))) {
            ann.settings[[i]]<-list(pch=pch, 
                col=lcols[(i-2) %% length(lcols) +1 ], 
                fill=lfills[(i-2) %% length(lfills) +1 ], 
                cex=cex, label=label.default);
            ann.settings[[i]]$label$show <- T
        }
        names(ann.settings)<-levels(grp)
    }
    for(i in 1:length(ann.settings)) {
        if (i>1) {ann.settings[[i]] <- modifyList(ann.settings[[i]], ann.default)}
        ann.settings[[i]]$label <- modifyList(ann.settings[[i]]$label, 
            parse.label(ann.settings[[i]]$label, levels(grp)[i]))
    }
    if(is.list(annotate) && length(annotate)>1) {
        user.cols <- 2:length(annotate)
        ann.cols <- c()
        if(!is.null(names(annotate[-1])) && all(names(annotate[-1])!="")) {
            ann.cols<-match(names(annotate)[-1], names(ann.settings))
        } else {
            ann.cols<-user.cols-1
        }
        for(i in seq_along(user.cols)) {
            if(!is.null(annotate[[user.cols[i]]]$label)) {
                annotate[[user.cols[i]]]$label<-parse.label(annotate[[user.cols[i]]]$label, 
                    levels(grp)[ann.cols[i]])
            }
            ann.settings[[ann.cols[i]]]<-modifyList(ann.settings[[ann.cols[i]]], 
                annotate[[user.cols[i]]])
        }
    }
    rm(annotate)
    #reduce number of points plotted
    if(should.thin) {
        thinned <- unique(data.frame(
            logp=round(-log10(pvalue),thin.logp.places), 
            pos=round(genpos,thin.pos.places), 
            chr=chr,
            grp=grp)
        )
        logp <- thinned$logp
        genpos <- thinned$pos
        chr <- thinned$chr
        grp <- thinned$grp
        rm(thinned)
    } else {
        logp <- -log10(pvalue)
    }
    rm(pos, pvalue)
    gc()
    #custom axis to print chromosome names
    axis.chr <- function(side,...) {
        if(side=="bottom") {
            panel.axis(side=side, outside=T,
                at=((posmax+posmin)/2+posshift),
                labels=levels(chr), 
                ticks=F, rot=0,
                check.overlap=F
            )
        } else if (side=="top" || side=="right") {
            panel.axis(side=side, draw.labels=F, ticks=F);
        }
        else {
            axis.default(side=side,...);
        }
     }
    #make sure the y-lim covers the range (plus a bit more to look nice)
    prepanel.chr<-function(x,y,...) { 
        A<-list();
        maxy<-ceiling(max(y, ifelse(!is.na(sig.level), -log10(sig.level), 0)))+.5;
        A$ylim=c(0,maxy);
        A;
    }

    xyplot(logp~genpos, chr=chr, groups=grp,
        axis=axis.chr, ann.settings=ann.settings, 
        prepanel=prepanel.chr, scales=list(axs="i"),
        panel=function(x, y, ..., getgenpos) {
            if(!is.na(sig.level)) {
                #add significance line (if requested)
                panel.abline(h=-log10(sig.level), lty=2);
            }
            panel.superpose(x, y, ..., getgenpos=getgenpos);
            if(!is.null(panel.extra)) {
                panel.extra(x,y, getgenpos, ...)
            }
        },
        panel.groups = function(x,y,..., subscripts, group.number) {
            A<-list(...)
            #allow for different annotation settings
            gs <- ann.settings[[group.number]]
            A$col.symbol <- gs$col[(as.numeric(chr[subscripts])-1) %% length(gs$col) + 1]    
            A$cex <- gs$cex[(as.numeric(chr[subscripts])-1) %% length(gs$cex) + 1]
            A$pch <- gs$pch[(as.numeric(chr[subscripts])-1) %% length(gs$pch) + 1]
            A$fill <- gs$fill[(as.numeric(chr[subscripts])-1) %% length(gs$fill) + 1]
            A$x <- x
            A$y <- y
            do.call("panel.xyplot", A)
            #draw labels (if requested)
            if(gs$label$show) {
                gt<-gs$label
                names(gt)[which(names(gt)=="text")]<-"labels"
                gt$show<-NULL
                if(is.character(gt$x) | is.character(gt$y)) {
                    peak = which.max(y)
                    center = mean(range(x))
                    if (is.character(gt$x)) {
                        if(gt$x=="peak") {gt$x<-x[peak]}
                        if(gt$x=="center") {gt$x<-center}
                    }
                    if (is.character(gt$y)) {
                        if(gt$y=="peak") {gt$y<-y[peak]}
                    }
                }
                if(is.list(gt$x)) {
                    gt$x<-A$getgenpos(gt$x[[1]],gt$x[[2]])
                }
                do.call("panel.text", gt)
            }
        },
        xlab=xlab, ylab=ylab, 
        panel.extra=panel.extra, getgenpos=getGenPos, ...
    );
}

# main function that is called from below, the plot itself is above function manhattan_plot_graph
manhattan_plot<-function(DEG, output_prefix, chrnum){
  #select columns to include in manhattan plot
  myTopHits.df <- DEG %>% dplyr::select(chromosomenumber, txStart, pvaluebound)
  #filter to only include chromosomes 1-25
  myTopHits.df <- myTopHits.df %>% dplyr::filter(chromosomenumber %in% 1:25)
  #make chromosomes plotted in order
  myTopHits.df$chromosomenumber <- factor(myTopHits.df$chromosomenumber, levels = 1:25)
  #omit NA values
  myTopHits.df <- na.omit(myTopHits.df)
  #make colors
  manhattancol <- rep(c("lightgrey", "darkgrey"), length.out = 25)
  manhattancol[chrnum] <- "#21908CFF"
  # Set up the PNG device with the desired dimensions
  png(filename = paste0(output_prefix, "_manhattan_plot.png"), width = 1000, height = 500)
  #can filter at this step if desired
  manhattan <- myTopHits.df 
  #make manhattan plot
  manhattan_plot_graph(manhattan$chromosomenumber, manhattan$txStart, manhattan$pvaluebound, should.thin=F, col=manhattancol)
  # Close the PNG device to save the plot
  dev.off()
}
```

##### 2.2.2 Check for enrichment of one chromosome in DEGs

```
chr_gsea <- function(output_prefix, annotation_tsv, DEG, chrnum, mutgene){
  # Import annotations for chromosome and TSS
  chromosomeinfo <- readr::read_tsv(file = annotation_tsv)
  # Keep chromosome and gene name
  chromosomeinfo <- chromosomeinfo %>% select(chrom, name2)
  #change column names
  colnames(chromosomeinfo) <- c("chromosome", "genename")
  #select only chromosomes 1-25
  chromosomeinfo <- chromosomeinfo %>% dplyr::filter(chromosome %in% paste("chr", 1:25, sep=""))
  #keep only distinct rows
  chromosomeinfo <- distinct(chromosomeinfo)  
  
  # Perform GSEA using clusterProfiler
  #keep only the columns we need for GSEA
  mydata.df.sub <- dplyr::select(DEG, LLgeneAbbrev, log2FoldChange)
  #sort by abs(foldchange)
  mydata.df.sub$log2FoldChange <- abs(mydata.df.sub$log2FoldChange)
  # construct a named vector
  mydata.gsea <- mydata.df.sub$log2FoldChange
  names(mydata.gsea) <- as.character(mydata.df.sub$LLgeneAbbrev)
  mydata.gsea <- sort(mydata.gsea, decreasing = TRUE)
  # run GSEA using the 'GSEA' function from clusterProfiler
  myGSEA.res <- GSEA(mydata.gsea, TERM2GENE=chromosomeinfo, verbose=FALSE, scoreType="pos", maxGSSize=2000, pvalueCutoff = 1)
  myGSEA.df <- as_tibble(myGSEA.res@result)
  write.csv(myGSEA.df, file=paste0(output_prefix,"_chrGSEA.csv"))
  # view results as an interactive table
  #datatable(myGSEA.df, 
  #          extensions = c('KeyTable', "FixedHeader"), 
  #          options = list(keys = TRUE, searchHighlight = TRUE, pageLength = 10, lengthMenu = c("10", "25", "50", "100"))) %>%
  #  formatRound(columns=c(2:10), digits=2)
  
  # create enrichment plots using the enrichplot package
  enrichment_plot <- gseaplot2(
        myGSEA.res, 
        geneSetID = c(1), #can choose multiple signatures to overlay in this plot
        pvalue_table = FALSE, #can set this to FALSE for a cleaner plot
        title = myGSEA.res$Description[1]) #can also turn off this title
  # Capture the plot
  g <- grid::grid.grabExpr(grid::grid.draw(enrichment_plot))
  # Save the enrichment plot
  ggsave(filename = paste0(output_prefix, "_enrichment_plot.png"), 
         plot = g, 
         width = 10, 
         height = 5, 
         dpi = 300)
  # Plot fold change of genes on mutant chromosome in chromosomal order
  #filter to only include those with logfc > 1
  myTopHits.df <- DEG %>% dplyr::filter(abs(log2FoldChange) > 1)
  #get names of genes with logfc > 1
  genestoplot <- unique(myTopHits.df$LLgeneAbbrev)
  #filter to only include all data ????
  myTopHits.df <- DEG # This seems not necessary when not filtering
  #filter to only include genes on chr 13
  myTopHits.df <- myTopHits.df %>% dplyr::filter(chrom %in% c(paste0("chr",chrnum)))
  #change back to fold change
  myTopHits.df <- myTopHits.df %>% mutate(foldchange = 2^log2FoldChange)
  #change the order of the genes
  myTopHits.df <- myTopHits.df %>% arrange(txStart)
  #filter to include only those with log2fc >1
  myTopHits.df <- myTopHits.df %>% dplyr::filter(LLgeneAbbrev %in% genestoplot)
  #make genes into a factor
  myTopHits.df$LLgeneAbbrev <- factor(myTopHits.df$LLgeneAbbrev, levels=unique(myTopHits.df$LLgeneAbbrev))

  chrplot <- ggplot(myTopHits.df %>% dplyr::filter(chrom == paste0("chr",chrnum)), aes(y=log2FoldChange, x=LLgeneAbbrev, fill=factor(ifelse(LLgeneAbbrev == mutgene, "Highlighted", "Normal")))) + 
    geom_bar(position="dodge", stat="summary", ) +
    ylab("log2 fold change") +
    theme_bw() +
    theme(axis.text.x = element_text(angle = 45, hjust=1), legend.position = "none", axis.title.x=element_blank()) +
    scale_fill_manual(values = c("#21908CFF","gray35"))
  ggsave(filename = paste0(output_prefix, "_chrplot.png"), plot = chrplot, width = 13, height = 7, dpi = 300)
  # Plot fold change of genes on mutant chromosome by TSS
  #can filter at this step
  myTopHits.df <- DEG
  #filter to only include genes on mutant chr
  myTopHits.df <- myTopHits.df %>% dplyr::filter(chrom == paste0("chr",chrnum))
  #change the order of the genes
  myTopHits.df <- myTopHits.df %>% arrange(txStart)
  chrTSSplot <- ggplot(myTopHits.df, aes(y=log2FoldChange, x=txStart, color=factor(ifelse(LLgeneAbbrev == mutgene, "Highlighted", "Normal")))) + 
      geom_bar(stat="summary" ) +
      ylab("log2 fold change") +
      xlab("TSS base position") +
      theme_bw() +
      theme(axis.text.x = element_text(angle = 45, hjust=1), legend.position = "none") +
      scale_color_manual(values = c("#21908CFF","gray35")) +
      scale_x_continuous(n.breaks = 20, labels = scales::comma, expand = c(0, 0))
  ggsave(filename = paste0(output_prefix, "_chrTSSplot.png"), plot = chrTSSplot, width = 13, height = 7, dpi = 300)
}
```

#### 2.3 Analysis of DEGs

##### 2.3.1 GO analysis

```
go_analysis <- function(output_prefix, DEG, chrnum){
  # Get genes with pvalue < 0.01 and exclude same chromosome as gene
  myTopHits.GO <- DEG %>% filter(pvalue < 0.01 & chromosomenumber != chrnum)
  write.csv(myTopHits.GO, paste0(output_prefix, "_genesforGO.csv"))
  justgenes <- dplyr::pull(myTopHits.GO, LLgeneAbbrev)
  # Perform GO enrichment analysis
  enrich_results <- enrichGO(
    gene = justgenes,
    OrgDb = org.Dr.eg.db,
    keyType = "SYMBOL",  # Adjust if using a different keyType
    ont = "BP",
    pAdjustMethod = "BH",
    qvalueCutoff = 0.05
  )
  # Print and visualize results
  results_df <- as.data.frame(enrich_results)
  write.csv(results_df, file = paste0(output_prefix,"_GO_clusterprofiler_results.csv"), row.names = FALSE)

  return(results_df)
}
```

##### 2.3.2 GSEA with C5

```
c5_gsea <- function(DEGs, output_prefix, chrnum) {
  #we want the zebrafish signatures
  dr_gsea <- msigdbr(species = "Danio rerio") #gets all collections/signatures with zebrafish
  #look at the categories and subcategories of signatures available
  dr_gsea %>% 
    dplyr::distinct(gs_cat, gs_subcat) %>% 
    dplyr::arrange(gs_cat, gs_subcat)
  # choose a specific msigdb collection/subcollection
  dr_gsea_c5 <- msigdbr(species = "Danio rerio", category = "C5") %>% # choose  msigdb collection of interest
  dplyr::select(gs_name, gene_symbol) #just get the columns corresponding to signature name and gene symbols of genes in each signature 
  #filter to exclude same chromosome genes
  mydata.df.sub <- DEGs %>% filter(chromosomenumber != chrnum)
  # pull out data that we need
  mydata.df.sub <- dplyr::select(mydata.df.sub, LLgeneAbbrev, log2FoldChange)
  #get rid of duplicates
  mydata.df.sub <- mydata.df.sub %>% dplyr::filter(LLgeneAbbrev %in% mydata.df.sub$LLgeneAbbrev[!duplicated(mydata.df.sub$LLgeneAbbrev)])
  # construct a named vector
  mydata.gsea <- mydata.df.sub$log2FoldChange
  names(mydata.gsea) <- as.character(mydata.df.sub$LLgeneAbbrev)
  mydata.gsea <- sort(mydata.gsea, decreasing = TRUE)
  # run GSEA with C5
  myGSEA.res <- GSEA(mydata.gsea, TERM2GENE=dr_gsea_c5, verbose=FALSE, seed=TRUE)
  #convert to DF
  myGSEA.df <- as_tibble(myGSEA.res)
  #look at top terms
  datatable(myGSEA.df, 
            extensions = c('KeyTable', "FixedHeader"), 
            options = list(keys = TRUE, searchHighlight = TRUE, pageLength = 10, lengthMenu = c("10", "25", "50", "100"))) %>%
    formatRound(columns=c(2:10), digits=2)
  #export
  write.csv(myGSEA.df, paste0(output_prefix,"_GSEA-c5.csv"))
  return(myGSEA.res)
}
```

##### 2.3.3 Use single cell markers from 5 dpf zebrafish brain to perform GSEA

```
scGSEA <- function(output_prefix, DEG, chrnum, hmapFCfilter = 0.5){
  #import single cell markers
  singlecellmarkers <- read.csv("zf5dpf_markersforGSEA.csv")
  #select relevant columns
  singlecellmarkers <- singlecellmarkers %>% dplyr::select(cluster.description, gene)
  #filter to exclude same chr genes
  mydata.df.sub <- DEG %>% filter(chromosomenumber != chrnum)
  # Pull out just the columns corresponding to gene symbols and LogFC
  mydata.df.sub <- dplyr::select(mydata.df.sub, LLgeneAbbrev, log2FoldChange)
  #get rid of duplicates
  mydata.df.sub <- mydata.df.sub %>% dplyr::filter(LLgeneAbbrev %in% mydata.df.sub$LLgeneAbbrev[!duplicated(mydata.df.sub$LLgeneAbbrev)])
  # construct a named vector
  mydata.gsea <- mydata.df.sub$log2FoldChange
  names(mydata.gsea) <- as.character(mydata.df.sub$LLgeneAbbrev)
  mydata.gsea <- sort(mydata.gsea, decreasing = TRUE)
  # run GSEA with single cell markers
  myGSEA.res <- GSEA(mydata.gsea, TERM2GENE=singlecellmarkers, verbose=FALSE, seed=TRUE, minGSSize = 80)
  myGSEA.df <- as_tibble(myGSEA.res@result)
  #export
  write.csv(myGSEA.df, file=paste0(output_prefix,"_singlecellGSEA.csv"))
  #import
  #myGSEA.df <- read.csv(file=paste0(output_prefix,"_singlecellGSEA.csv"))[,2:12]
  datatable(myGSEA.df, 
          extensions = c('KeyTable', "FixedHeader"), 
          options = list(keys = TRUE, searchHighlight = TRUE, pageLength = 10, lengthMenu = c("10", "25", "50", "100"))) %>%
    formatRound(columns=c(2:10), digits=2)
  myheatcolors3 <- brewer.pal(name="RdBu", n=11)
  #make network plot
  myGSEA.res <- pairwise_termsim(myGSEA.res)
  #print(myGSEA.res)
  #print(length(myGSEA.res))
  network <- emapplot(myGSEA.res, color="NES", categorySize="p.adjust", showCategory = nrow(myGSEA.res)) #58??? All the showCategory points need to be fixed
  #get data of out network plot
  network <- ggplot_build(network)
  networkdata <- network$plot$data
  #make network plot
  singlecell_network <- ggraph(networkdata) + 
    geom_edge_link(alpha=.8, aes_(width=~I(width)), colour='darkgrey') + 
    geom_node_point(aes(colour = color, size=size)) +
    geom_node_text(aes(label=name), repel=TRUE) + 
    theme_void() +
    scale_color_gradientn(colors = myheatcolors3, limit=c(-4,4), name="NES")
  ggsave(filename = paste0(output_prefix, "_scRNA5dpf_network.png"), plot = singlecell_network, width = 10, height = 6, dpi = 300)
  #singlecell_network
  #export 1000x600
#make heatmap
  p3 <- heatplot(myGSEA.res, foldChange=mydata.gsea, showCategory = nrow(myGSEA.res)) + ggplot2::coord_flip()
  #get data of out heatmap
  p3 <- ggplot_build(p3)
  heatmapdata <- p3$plot$data
  #filter to only include fold changes > 0.5
  genestoinclude <- heatmapdata %>% dplyr::filter(foldChange > hmapFCfilter | foldChange < -hmapFCfilter)
  heatmapdata <- heatmapdata %>% dplyr::filter(Gene %in% genestoinclude$Gene)
  #use pivot wider to make untidy table
  heatmapdata <- heatmapdata %>% pivot_wider(names_from = categoryID, values_from  = foldChange, values_fill=NA) 
  #make matrix
  heatmapmatrix <- as.matrix(heatmapdata[,2:length(heatmapdata)])
  rownames(heatmapmatrix) <- heatmapdata$Gene
  #turn matrix into 1s and 0s and cluster
  heatmapmatrix<-ifelse(abs(heatmapmatrix) > 0,1,0)
  heatmapmatrix[is.na(heatmapmatrix)] = 0
 #cluster rows by pearson correlation
  hc <- hclust(as.dist(1-cor(heatmapmatrix, method="spearman")), method="average") #cluster columns by spearman correlation
  #cluster your selected genes
  #hr <- hclust(as.dist(1-cor(t(heatmapmatrix), method="pearson")), method="complete") #cluster rows by pearson correlation
  #turn heatmap data back into tidy table
  heatmapdata <- heatmapdata %>% pivot_longer(!Gene, names_to = "categoryID", values_to  = "foldChange") 
  #change order of data to be plotted
  heatmapdata$categoryID <- factor(heatmapdata$categoryID, levels = hc$labels)
  heatmapdata$Gene <- factor(heatmapdata$Gene, levels = rev(unique(heatmapdata$Gene)))
  #heatmapdata$Gene <- factor(heatmapdata$Gene, levels = hr$labels)
  #rename dataframe columns
  colnames(heatmapdata) <- c("Gene", "categoryID", "log2foldchange")
  singlecellheatmap <- ggplot(heatmapdata, aes(x = Gene, y = categoryID, fill = log2foldchange)) + 
    geom_tile(color="black") + 
    theme_classic() + 
    scale_fill_continuous_divergingx(palette = 'RdBu', mid = 0, na.value="white") +
    theme(axis.text.x = element_text(angle = 45, vjust = 1, hjust = 1), axis.title.x=element_blank(), axis.title.y=element_blank()) 
  flipped_singlecellheatmap <- singlecellheatmap + ggplot2::coord_flip()
  ggsave(filename = paste0(output_prefix, "_scRNA5dpf_heatmap.png"), plot = flipped_singlecellheatmap, width = 20, height = 35, dpi = 300)
  #export 1200x2000
}
```

##### 2.3.4 Use single cell markers from Danio Cell to perform GSEA

```
scGSEA_danio <- function(output_prefix, DEG, chrnum, filter_string="", hmapFCfilter = 0.5){
  #import single cell markers
  # PATH MUST BE REPLACED FOR THE MARKER SET
  singlecellmarkers <- read.csv("/Users/sthyme/daniocell/markerlist-by500P_combo_neu-eye-ot-tas-gli-hem-ion.csv")
  #select relevant columns
  singlecellmarkers <- singlecellmarkers %>% dplyr::select(cluster, gene)
  #filter to exclude same chr genes
  mydata.df.sub <- DEG %>% filter(chromosomenumber != chrnum)
  # Pull out just the columns corresponding to gene symbols and LogFC
  mydata.df.sub <- dplyr::select(mydata.df.sub, LLgeneAbbrev, log2FoldChange)
  #get rid of duplicates
  mydata.df.sub <- mydata.df.sub %>% dplyr::filter(LLgeneAbbrev %in% mydata.df.sub$LLgeneAbbrev[!duplicated(mydata.df.sub$LLgeneAbbrev)])
  # construct a named vector
  mydata.gsea <- mydata.df.sub$log2FoldChange
  names(mydata.gsea) <- as.character(mydata.df.sub$LLgeneAbbrev)
  mydata.gsea <- sort(mydata.gsea, decreasing = TRUE)
  # run GSEA with single cell markers
  myGSEA.res <- GSEA(mydata.gsea, TERM2GENE=singlecellmarkers, verbose=FALSE, seed=TRUE, minGSSize = 80)
  myGSEA.df <- as_tibble(myGSEA.res@result)
  #export
  write.csv(myGSEA.df, file=paste0(output_prefix,"_singlecellGSEA-danio-byp.csv"))
  #import
  #myGSEA.df <- read.csv(file=paste0(output_prefix,"_singlecellGSEA-danio-byp.csv"))[,2:12]
  datatable(myGSEA.df, 
          extensions = c('KeyTable', "FixedHeader"), 
          options = list(keys = TRUE, searchHighlight = TRUE, pageLength = 10, lengthMenu = c("10", "25", "50", "100"))) %>%
    formatRound(columns=c(2:10), digits=2)
  myheatcolors3 <- brewer.pal(name="RdBu", n=11)
  #make network plot
  myGSEA.res <- pairwise_termsim(myGSEA.res)
  network <- emapplot(myGSEA.res, color="NES", categorySize="p.adjust", showCategory = nrow(myGSEA.res)) #58??? All the showCategory points need to be fixed
  #get data of out network plot
  network <- ggplot_build(network)
  networkdata <- network$plot$data
  #make network plot
  singlecell_network <- ggraph(networkdata) + 
    geom_edge_link(alpha=.8, aes_(width=~I(width)), colour='darkgrey') + 
    geom_node_point(aes(colour = color, size=size)) +
    geom_node_text(aes(label=name), repel=TRUE) + 
    theme_void() +
    scale_color_gradientn(colors = myheatcolors3, limit=c(-4,4), name="NES")
  ggsave(filename = paste0(output_prefix, "_scRNA-danio-byp_network.png"), plot = singlecell_network, width = 10, height = 6, dpi = 300)
  #make heatmap
  p3 <- heatplot(myGSEA.res, foldChange=mydata.gsea, showCategory = nrow(myGSEA.res)) + ggplot2::coord_flip()
  #get data of out heatmap
  p3 <- ggplot_build(p3)
  heatmapdata <- p3$plot$data
  # Define the string to filter by
  if (filter_string != "") {
    # Filter to include only the categories containing the given string
    heatmapdata <- heatmapdata %>%
      dplyr::filter(str_detect(categoryID, filter_string))
  }
  #filter to only include fold changes > 0.5
  genestoinclude <- heatmapdata %>% dplyr::filter(foldChange > hmapFCfilter | foldChange < -hmapFCfilter)
  heatmapdata <- heatmapdata %>% dplyr::filter(Gene %in% genestoinclude$Gene)
  #use pivot wider to make untidy table
  heatmapdata <- heatmapdata %>% pivot_wider(names_from = categoryID, values_from  = foldChange, values_fill=NA) 
  #make matrix
  heatmapmatrix <- as.matrix(heatmapdata[,2:length(heatmapdata)])
  rownames(heatmapmatrix) <- heatmapdata$Gene
  #turn matrix into 1s and 0s and cluster
  heatmapmatrix<-ifelse(abs(heatmapmatrix) > 0,1,0)
  heatmapmatrix[is.na(heatmapmatrix)] = 0
 #cluster rows by pearson correlation
  hc <- hclust(as.dist(1-cor(heatmapmatrix, method="spearman")), method="average") #cluster columns by spearman correlation
  #cluster your selected genes
  #hr <- hclust(as.dist(1-cor(t(heatmapmatrix), method="pearson")), method="complete") #cluster rows by pearson correlation
  #turn heatmap data back into tidy table
  heatmapdata <- heatmapdata %>% pivot_longer(!Gene, names_to = "categoryID", values_to  = "foldChange") 
  #change order of data to be plotted
  heatmapdata$categoryID <- factor(heatmapdata$categoryID, levels = hc$labels)
  heatmapdata$Gene <- factor(heatmapdata$Gene, levels = rev(unique(heatmapdata$Gene)))
  #heatmapdata$Gene <- factor(heatmapdata$Gene, levels = hr$labels)
  #rename dataframe columns
  colnames(heatmapdata) <- c("Gene", "categoryID", "log2foldchange")
  singlecellheatmap <- ggplot(heatmapdata, aes(x = Gene, y = categoryID, fill = log2foldchange)) + 
    geom_tile(color="black") + 
    theme_classic() + 
    scale_fill_continuous_divergingx(palette = 'RdBu', mid = 0, na.value="white") +
    theme(axis.text.x = element_text(angle = 45, vjust = 1, hjust = 1), axis.title.x=element_blank(), axis.title.y=element_blank()) 
  flipped_singlecellheatmap <- singlecellheatmap + ggplot2::coord_flip()
  ggsave(filename = paste0(output_prefix, filter_string, "_scRNA-danio-byp_heatmap.png"), plot = flipped_singlecellheatmap, width = 20, height = 35, dpi = 300)
}
```

##### 2.3.5 Use ZFA anatomy to perform GSEA

```
anatGSEA <- function(output_prefix, DEG, chrnum, hmapFCfilter = 0.2){
  CNStermsGSEA <- read.csv("CNStermsGSEA.csv")[,2:3]
  headtermsGSEA <- read.csv("headtermsGSEA.csv")[,2:3]
  fullsetGSEA <- read.csv("fullsetGSEA.csv")[,2:3]
  # Pull out just the columns corresponding to gene symbols and LogFC for at least one pairwise comparison for the enrichment analysis
  GSEAgenes <- DEG %>% dplyr::filter(chromosomenumber != chrnum)
  mydata.df.sub <- dplyr::select(GSEAgenes, LLgeneAbbrev, log2FoldChange)
  #get rid of duplicates
  mydata.df.sub <- mydata.df.sub %>% dplyr::filter(LLgeneAbbrev %in% mydata.df.sub$LLgeneAbbrev[!duplicated(mydata.df.sub$LLgeneAbbrev)])
  # construct a named vector
  mydata.gsea <- mydata.df.sub$log2FoldChange
  names(mydata.gsea) <- as.character(mydata.df.sub$LLgeneAbbrev)
  mydata.gsea <- sort(mydata.gsea, decreasing = TRUE)

  #options for doing GSEA using ZFA
  #CNStermsGSEA
  #headtermsGSEA
  #fullsetGSEA

  # run GSEA with CNS terms 
  myGSEA.res <- GSEA(mydata.gsea, TERM2GENE=CNStermsGSEA, verbose=FALSE, seed=TRUE, minGSSize = 80)
  myGSEA.df <- as_tibble(myGSEA.res@result)

  #export
  write.csv(myGSEA.df, file=paste0(output_prefix,"_CNS_GSEA.csv"))
  #import
  #myGSEA.df <- read.csv(file=paste0(output_prefix,"_CNS_GSEA.csv"))[,2:12]

  # view results as an interactive table
  #datatable(myGSEA.df, 
  #          extensions = c('KeyTable', "FixedHeader"), 
  #          options = list(keys = TRUE, searchHighlight = TRUE, pageLength = 10, lengthMenu = c("10", "25", "50", "100"))) %>%
  #  formatRound(columns=c(2:10), digits=2)

  #make network plot
  myGSEA.res <- pairwise_termsim(myGSEA.res)
  network <- emapplot(myGSEA.res, color="NES", categorySize="p.adjust", showCategory = nrow(myGSEA.res))
  #get data of out network plot
  network <- ggplot_build(network)
  networkdata <- network$plot$data
  #get color
  myheatcolors3 <- brewer.pal(name="RdBu", n=11)
  #make network plot
  anat_network <- ggraph(networkdata) + 
    geom_edge_link(alpha=.8, aes_(width=~I(width)), colour='darkgrey') + 
    geom_node_point(aes(colour = color, size=size)) +
    geom_node_text(aes(label=name), repel=TRUE) + 
    theme_void() +
    scale_color_gradientn(colors = myheatcolors3, limit=c(-1.5,1.5), name="NES")
  ggsave(filename = paste0(output_prefix, "_anatzfa_network.png"), plot = anat_network, width = 10, height = 6, dpi = 300)
  
  #make heatmap
  p3 <- heatplot(myGSEA.res, foldChange=mydata.gsea, showCategory = nrow(myGSEA.res)) + ggplot2::coord_flip()
  #get data of out heatmap
  p3 <- ggplot_build(p3)
  heatmapdata <- p3$plot$data
  #filter to only include fold changes > 0.5
  genestoinclude <- heatmapdata %>% dplyr::filter(foldChange > hmapFCfilter | foldChange < -hmapFCfilter)
  heatmapdata <- heatmapdata %>% dplyr::filter(Gene %in% genestoinclude$Gene)
  #use pivot wider to make untidy table
  heatmapdata <- heatmapdata %>% pivot_wider(names_from = categoryID, values_from  = foldChange, values_fill=NA) 
  #make matrix
  heatmapmatrix <- as.matrix(heatmapdata[,2:length(heatmapdata)])
  rownames(heatmapmatrix) <- heatmapdata$Gene
  #turn matrix into 1s and 0s and cluster
  heatmapmatrix<-ifelse(abs(heatmapmatrix) > 0,1,0)
  heatmapmatrix[is.na(heatmapmatrix)] = 0
  #cluster rows by pearson correlation
  #hc <- hclust(as.dist(1-cor(heatmapmatrix, method="spearman")), method="average") #cluster columns by spearman correlation
  #turn heatmap data back into tidy table
  heatmapdata <- heatmapdata %>% pivot_longer(!Gene, names_to = "categoryID", values_to  = "foldChange") 
  #change order of data to be plotted
  heatmapdata$Gene <- factor(heatmapdata$Gene, levels = rev(unique(heatmapdata$Gene)))
  #rename dataframe columns
  colnames(heatmapdata) <- c("Gene", "categoryID", "log2foldchange")
  anatheatmap <- ggplot(heatmapdata, aes(x = Gene, y = categoryID, fill = log2foldchange)) + 
    geom_tile(color="black") + 
    theme_classic() + 
    scale_fill_continuous_divergingx(palette = 'RdBu', mid = 0, na.value="white") +
    theme(axis.text.x = element_text(angle = 45, vjust = 1, hjust = 1), axis.title.x=element_blank(), axis.title.y=element_blank())
  flipped_anatheatmap <- anatheatmap + ggplot2::coord_flip()
  ggsave(filename = paste0(output_prefix, "_anatzfa_heatmap.png"), plot = flipped_anatheatmap, width = 12, height = 20, dpi = 300)
}
```

#### 2.4 Make plots for C5 GSEA

```
gsea_plot_sub <- function(output_prefix, myGSEA.res.filter, DEG, chrnum, nolabel=FALSE, hmapFCfilter){
    #print(myGSEA.res.filter)
    myGSEA.res.filter <- pairwise_termsim(myGSEA.res.filter)
    #valid_terms <- intersect(as.character(myGSEA.res.filter$Description), rownames(myGSEA.res.filter@termsim))
    #myGSEA.res.filter <- myGSEA.res.filter[myGSEA.res.filter$Description %in% valid_terms, ]
    #print(myGSEA.res.filter)
    network <- emapplot(myGSEA.res.filter, color="NES", categorySize="p.adjust", showCategory=200)##nrow(myGSEA.res.filter))
    #get data of out network plot
    network <- ggplot_build(network)
    networkdata <- network$plot$data
    #get color
    myheatcolors3 <- brewer.pal(name="RdBu", n=11) # this should perhaps not be within the loop and repeated
    #make plot
    if(nolabel==TRUE){
      networkplot <- ggraph(networkdata) + 
        geom_edge_link(alpha=.8, aes_(width=~I(width)), colour='darkgrey') + 
        geom_node_point(aes(colour = color, size=size)) +
        theme_void() +
        scale_color_gradientn(colors = myheatcolors3, limit=c(-3.5,3.5), name="NES")
    } else {
        networkplot <- ggraph(networkdata) + 
        geom_edge_link(alpha=.8, aes_(width=~I(width)), colour='darkgrey') + 
        geom_node_point(aes(colour = color, size=size)) +
        geom_node_text(aes(label=name), repel=TRUE) + 
        theme_void() +
        scale_color_gradientn(colors = myheatcolors3, limit=c(-3.5,3.5), name="NES")
    }
    ggsave(filename = paste0(output_prefix, "_GSEA-c5_network.png"), plot = networkplot, width = 15, height = 15, dpi = 300)
    ggsave(filename = paste0(output_prefix, "_GSEA-c5-large_network.png"), plot = networkplot, width = 30, height = 30, dpi = 300)
    
    # add heatmap
    mydata.df.sub <- DEG %>% filter(chromosomenumber != chrnum)
    # Pull out just the columns corresponding to gene symbols and LogFC
    mydata.df.sub <- dplyr::select(mydata.df.sub, LLgeneAbbrev, log2FoldChange)
    #get rid of duplicates
    mydata.df.sub <- mydata.df.sub %>% dplyr::filter(LLgeneAbbrev %in% mydata.df.sub$LLgeneAbbrev[!duplicated(mydata.df.sub$LLgeneAbbrev)])
    # construct a named vector
    mydata.gsea <- mydata.df.sub$log2FoldChange
    names(mydata.gsea) <- as.character(mydata.df.sub$LLgeneAbbrev)
    mydata.gsea <- sort(mydata.gsea, decreasing = TRUE)
    p3 <- heatplot(myGSEA.res.filter, foldChange=mydata.gsea, showCategory=nrow(myGSEA.res.filter)) + ggplot2::coord_flip() # CHECK IF LENGTH IS RIGHT
    p3 <- ggplot_build(p3)
    heatmapdata <- p3$plot$data
    #filter to only include fold changes > 0.5
    genestoinclude <- heatmapdata %>% dplyr::filter(foldChange > hmapFCfilter | foldChange < -hmapFCfilter)
    heatmapdata <- heatmapdata %>% dplyr::filter(Gene %in% genestoinclude$Gene)
    #choose which clusters to keep in heatmap
    #heatmapdata <- heatmapdata %>% dplyr::filter(categoryID %in% networkterms[1:100]) # I DONT KNOW WHAT THIS IS FOR
    #use pivot wider to make untidy table
    heatmapdata <- heatmapdata %>% pivot_wider(names_from = categoryID, values_from  = foldChange, values_fill=NA)
    #make matrix
    heatmapmatrix <- as.matrix(heatmapdata[,2:length(heatmapdata)])
    rownames(heatmapmatrix) <- heatmapdata$Gene
    #turn matrix into 1s and 0s and cluster
    heatmapmatrix<-ifelse(abs(heatmapmatrix) > 0,1,0)
    #cluster your selected genes
    heatmapmatrix[is.na(heatmapmatrix)] = 0
    #heatmapmatrix[is.infinite(heatmapmatrix)] <- max(heatmapmatrix[!is.infinite(heatmapmatrix)], na.rm = TRUE)
    # Define a small random value to add
    # BELOW IS A POSSIBLE SOLUTION IF THE HEATMAP KEEPS SAYING THERE ARE NAN OR INF VALUES BECAUSE VALUES ARE ALL THE SAME IN A COLUMN
    ##set.seed(42)  # Set seed for reproducibility
    ##random_small_values <- matrix(runif(length(heatmapmatrix), min = 0, max = 1e-8), 
    ##                           nrow = nrow(heatmapmatrix), 
    ##                           ncol = ncol(heatmapmatrix))
    # Add the random small values to the original matrix
    ##heatmapmatrix <- heatmapmatrix + random_small_values
    hc <- hclust(as.dist(1-cor(heatmapmatrix, method="spearman")), method="average") #cluster columns by spearman correlation
    #turn heatmap data back into tidy table
    heatmapdata <- heatmapdata %>% pivot_longer(!Gene, names_to = "categoryID", values_to  = "foldChange") 
    #change order of data to be plotted
    heatmapdata$categoryID <- factor(heatmapdata$categoryID, levels = hc$labels)
    heatmapdata$Gene <- factor(heatmapdata$Gene, levels = rev(unique(heatmapdata$Gene)))
    #rename dataframe columns
    colnames(heatmapdata) <- c("Gene", "categoryID", "log2foldchange")
    c5_heatmap <- ggplot(heatmapdata, aes(x = Gene, y = categoryID, fill = log2foldchange)) + 
      geom_tile(color="black") + 
      theme_classic() + 
      scale_fill_continuous_divergingx(palette = 'RdBu', mid = 0, na.value="white") +
      theme(axis.text.x = element_text(angle = 45, vjust = 1, hjust = 1), axis.title.x=element_blank(), axis.title.y=element_blank())
  flipped_c5_heatmap <- c5_heatmap + ggplot2::coord_flip()
  ggsave(filename = paste0(output_prefix, "_GSEA-c5_heatmap.png"), plot = flipped_c5_heatmap, width = 12, height = 20, dpi = 300)
}

gsea_plot <- function(output_prefix, networkterms, myGSEA.res, DEG, chrnum, hmapFCfilter = 0.5) {
  # make plots with all terms
  myGSEA.res.nofilter <- myGSEA.res
  gsea_plot_sub(paste0(output_prefix,"all-nolabel_"), myGSEA.res.nofilter, DEG, chrnum, TRUE, hmapFCfilter)
  gsea_plot_sub(paste0(output_prefix,"all-label_"), myGSEA.res.nofilter, DEG, chrnum, FALSE, hmapFCfilter)
  for (name in names(gsea_terms)) {
    sublist <- gsea_terms[[name]]
    myGSEA.res.filter <- myGSEA.res
    myGSEA.res.filter@result <- myGSEA.res.filter@result %>% dplyr::filter(ID %in% sublist)
    gsea_plot_sub(paste0(output_prefix, name, "_"), myGSEA.res.filter, DEG, chrnum, FALSE, hmapFCfilter)
  }
}
```

#### 2.5 Gene extraction from cluster profiler file

```
extract_genes_for_terms <- function(results_df, terms_of_interest) {
  # Check if the input terms are in the results_df
  if (!all(terms_of_interest %in% results_df$ID)) {
    stop("Some terms in 'terms_of_interest' are not found in 'results_df'")
  }
  # Extract the gene lists for the specified terms
  gene_lists <- results_df$geneID[results_df$ID %in% terms_of_interest]
  #print(gene_lists)
  # Create a named list to store genes for each term
  genes_for_terms <- list()
  for (term in terms_of_interest) {
    # Get the index of the term
    index <- which(results_df$ID == term)
    # Extract gene list for the term
    gene_list <- results_df$geneID[index]
    description <- results_df$Description[index]
    genes_for_terms[[description]] <- unlist(strsplit(gene_list, "/"))  # Split by "/"
  }
  return(genes_for_terms)
}
```

#### 2.6 Volcano plot with multiple customizations possible

```
# In the future, may need to add additional customizations such as for the y-limits etc
create_volcano_plot <- function(output_prefix, myTopHits.df, gene_sets, genestolabel, chrnum, color_vector, pabound = 0.01, log2fcbound = 1.0, xmaxb = 3.5, xlimb = 3.7, nudgex = 0.1, padj = TRUE) { # set padj to FALSE if you want to use pvalue instead
  # Ensure the column names are correctly named
  if (!"chrom" %in% colnames(myTopHits.df)) {
    stop("Input dataframe must contain a 'chrom' column")
  }
  # Add column about whether chr is same as input gene
  myTopHits.df <- myTopHits.df %>% dplyr::mutate(chrom = replace_na(chrom, "none"))
  myTopHits.df <- myTopHits.df %>% mutate(chromosome = ifelse(chrom == paste0("chr",chrnum), paste0("chromosome ",chrnum), "other chromosome"))
  # Label the genes with color
  labeled_genes <- lapply(names(gene_sets), function(category) {
    df_subset <- myTopHits.df %>% dplyr::filter(LLgeneAbbrev %in% gene_sets[[category]])
    df_subset$category <- category
    return(df_subset)
  })
  myTopHits.labels <- do.call(rbind, labeled_genes)
  # Filter all genes based on thresholds
  if (padj == TRUE) {
    myTopHits.filtered <- myTopHits.df %>%
      dplyr::filter(abs(log2FoldChange) > log2fcbound & padjbound < pabound)
  }
  else {
    myTopHits.filtered <- myTopHits.df %>%
      dplyr::filter(abs(log2FoldChange) > log2fcbound & pvaluebound < pabound)
  }
  myTopHits.labels.all <- dplyr::bind_rows(
    myTopHits.filtered %>% dplyr::filter(LLgeneAbbrev %in% myTopHits.df$LLgeneAbbrev),
    myTopHits.df %>% dplyr::filter(LLgeneAbbrev %in% genestolabel)
  ) %>% dplyr::distinct()
  # Make all points other
  myTopHits.df <- myTopHits.df %>% dplyr::mutate(mutation = "other")
  # Make the plot
  volcano <- ggplot() +
    annotate("rect", xmin = log2(1.5), xmax = xmaxb, ymin = -log10(0.05), ymax = 10, alpha = .15, fill = "grey") +
    annotate("rect", xmin = -log2(1.5), xmax = -xmaxb, ymin = -log10(0.05), ymax = 10, alpha = .15, fill = "grey") +
  #  geom_point(data = myTopHits.df, aes(y = -log10(padjbound), x = log2FoldChange, shape = chromosome, color = mutation), size = 2) +
  #  geom_point(data = myTopHits.labels, aes(y = -log10(padjbound), x = log2FoldChange, color = category, shape = chromosome), size = 2, show.legend = TRUE) +
    theme_bw() +
    coord_cartesian(xlim = c(-xlimb, xlimb), ylim = c(-0.5, 10.5), expand = FALSE) +
   # ylab("-log10(padj)") + 
    xlab("log2 fold change")
    # Add conditional layers
    if (padj == TRUE) {
      volcano <- volcano +
        ylab("-log10(padj)") + 
        geom_point(data = myTopHits.df, aes(y = -log10(padjbound), x = log2FoldChange, shape = chromosome, color = mutation), size = 2) +
        geom_point(data = myTopHits.labels, aes(y = -log10(padjbound), x = log2FoldChange, color = category, shape = chromosome), size = 2, show.legend = TRUE) +
        geom_label_repel(data = myTopHits.labels.all, aes(x = log2FoldChange, y = -log10(padjbound), label = LLgeneAbbrev), 
                         force = 2, nudge_y = -1, nudge_x = nudgex, size = 2.5, max.overlaps = Inf, show.legend = FALSE, color = "black")
    } else {
      volcano <- volcano +
        ylab("-log10(pvalue)") + 
        geom_point(data = myTopHits.df, aes(y = -log10(pvaluebound), x = log2FoldChange, shape = chromosome, color = mutation), size = 2) +
        geom_point(data = myTopHits.labels, aes(y = -log10(pvaluebound), x = log2FoldChange, color = category, shape = chromosome), size = 2, show.legend = TRUE) +
        geom_label_repel(data = myTopHits.labels.all, aes(x = log2FoldChange, y = -log10(pvaluebound), label = LLgeneAbbrev), 
                         force = 2, nudge_y = -1, nudge_x = nudgex, size = 2.5, max.overlaps = Inf, show.legend = FALSE, color = "black")
    }
  # Add scales
    volcano <- volcano +
      scale_color_manual(values = color_vector, name = "pathway") +
      scale_shape_manual(values = c(4, 16))
  
#  volcano <- ggplot() +
#    annotate("rect", xmin = log2(1.5), xmax = xmaxb, ymin = -log10(0.05), ymax = 10, alpha = .15, fill = "grey") +
#    annotate("rect", xmin = -log2(1.5), xmax = -xmaxb, ymin = -log10(0.05), ymax = 10, alpha = .15, fill = "grey") +
#    geom_point(data = myTopHits.df, aes(y = -log10(padjbound), x = log2FoldChange, shape = chromosome, color = mutation), size = 2) +
    #geom_point(data = myTopHits.df, aes(y = -log10(pvaluebound), x = log2FoldChange, shape = chromosome, color = mutation), size = 2) +
#    geom_point(data = myTopHits.labels, aes(y = -log10(padjbound), x = log2FoldChange, color = category, shape = chromosome), size = 2, show.legend = T) +
#    #geom_point(data = myTopHits.labels, aes(y = -log10(pvaluebound), x = log2FoldChange, color = category, shape = chromosome), size = 2, show.legend = T) +
#    theme_bw() +
#    coord_cartesian(xlim = c(-xlimb, xlimb), ylim = c(-0.5, 10.5), expand = FALSE) +
#    ylab("-log10(padj)") + 
#    xlab("log2 fold change") +
    
 #      geom_label_repel(data = myTopHits.labels.all, aes(x = log2FoldChange, y = -log10(padjbound), label = LLgeneAbbrev), 
#                      force = 2, nudge_y = -1, nudge_x = 0.1, size = 2.5, max.overlaps = Inf, show.legend = FALSE, color = "black") +
#                      theme_bw() +
#                      scale_color_manual(values = c("#AADC32FF", "#27AD81FF", "#472D7BFF", "gold", "darkgrey", "deepskyblue", "orange",  # HAVE TO MOVE THE GRAY FOR THE "OTHER" POINTS
 #                                  "#D697FF", "lightblue", "darkgrey", "darkred"), name = "pathway") +
#                      scale_shape_manual(values = c(4, 16))
  ggsave(filename = paste0(output_prefix, "_volcano.png"), plot = volcano, width = 10, height = 8, dpi = 300)
  #return(volcano)
}
```

##### 2.6.1 Make a heatmap from an input list of genes

```
gene_heatmap <- function(output_prefix, genecounts, mySelectedGenes){
  myheatcolors3 <- brewer.pal(name="RdBu", n=11)
  #filter replicate data with list of selected genes
  mySelectedGenes_exp <- genecounts %>% filter(LLgeneAbbrev %in% mySelectedGenes)
  #reorder data table based on myselectedgenes
  mySelectedGenes_exp <- mySelectedGenes_exp[match(mySelectedGenes, mySelectedGenes_exp$LLgeneAbbrev),]
  #turn replicate data into a matrix
  mySelectedGenes.matrix <- as.matrix(mySelectedGenes_exp[,4:ncol(mySelectedGenes_exp)])
  #add row names (genes) to data matrix
  rownames(mySelectedGenes.matrix) <- mySelectedGenes_exp$LLgeneAbbrev
  mySelectedGenes.matrix <- mySelectedGenes.matrix[!rowSums(is.na(mySelectedGenes.matrix)),]
  mySelectedGenes.matrix <- mySelectedGenes.matrix[rowSums(mySelectedGenes.matrix[])>0,]
  #you may (or may not) want to cluster your selected genes
  #TURNING IT OFF FOR NOW, COULD MAKE IT AN ARGUMENT
  #hr <- hclust(as.dist(1-cor(t(mySelectedGenes.matrix), method="pearson")), method="complete") #cluster rows by pearson correlation
#  hc <- hclust(as.dist(1-cor(mySelectedGenes.matrix, method="spearman")), method="average") #cluster columns by spearman correlation
  #make heatmap
  # Calculate the appropriate margins based on label lengths
  png(filename = paste0(output_prefix,"heatmap.png"), width = 1800, height = 900)
  # Automatically adjust margins based on the size of labels
  # Calculate the length of the longest row and column labels
  max_row_label_length <- max(nchar(rownames(mySelectedGenes.matrix)))
  max_col_label_length <- max(nchar(colnames(mySelectedGenes.matrix)))
  # Set margins dynamically based on label length
  par(mar = c(max_col_label_length * 0.5, max_row_label_length * 0.5, 4, 2))
  # Create the heatmap with dynamic label size
  heatmap.2(mySelectedGenes.matrix, 
          Rowv = NA,#as.dendrogram(hr), 
          Colv = NA, 
          col = myheatcolors3, 
          scale = "row", 
          density.info = "none", 
          trace = "none",
          cexRow = 1 - (max_row_label_length / 100),   # Dynamically adjust row label size
          cexCol = 1 - (max_col_label_length / 100))   # Dynamically adjust column label size
  dev.off()
  }
```

##### 2.6.2 Make bar plots from an input list of genes

```
gene_barplots <- function(output_prefix, genecounts, geneofinterest, controlave){
  #make new columns with mean counts of het samples
  genecountslogfc <- genecounts
  genecountslogfc$controlave <- rowMeans(genecountslogfc[,controlave])
  #divide columns by mean counts of the control sample
  genecountslogfc <- genecountslogfc %>% mutate(across(colnames(genecountslogfc)[4:ncol(genecountslogfc)], function(x) x/controlave))
  #pull out data for a select gene
  generepdata <- genecountslogfc %>% dplyr::filter(LLgeneAbbrev %in% geneofinterest)
  #get rid of unnecessary columns
  generepdata <- generepdata[,c(1,4:(ncol(genecountslogfc)-1))]
  #pivot longer to make tidy
  generepdata <- generepdata %>% pivot_longer(!LLgeneAbbrev, names_to = "sample", values_to = "foldchange")
  #add condition information based on sampleName
  generepdata$genotype <- generepdata$sample
  #substitute condition name based on replicate
  generepdata <- generepdata %>% mutate(genotype = str_replace_all(genotype, c("^het.*"="het", "^wt.*"="wt", "^hom.*"="hom")))
  #order data on plot
  generepdata$genotype <- factor(generepdata$genotype, levels=c("wt", "het", "hom"))
  #set colors
  colors <- c("#472D7BFF", "#FDE725FF", "#21908CFF") 
  #make an empty list
  plot_list = list()
  #make all the plots
  for (z in 1:length(geneofinterest)) {
    genedata2dpf <- generepdata %>% dplyr::filter(LLgeneAbbrev == geneofinterest[z])
    plot <- ggplot(genedata2dpf, aes(fill=genotype, y=foldchange, x=genotype)) + 
      geom_bar(position="dodge", stat="summary") +
      geom_point(position = position_dodge(width = .9)) + 
      labs(title = paste(geneofinterest[z])) +
      ylab("fold change") +
      theme_bw() +
      scale_fill_manual(values = colors) +
      theme(
        axis.text = element_text(size = 14),  # Adjust axis text size
        axis.title = element_text(size = 16)  # Adjust axis title size
      )
    plot_list[[z]] <- plot
  }
  # Save plots to svg Makes a separate file for each plot.
  for (i in 1:length(geneofinterest)) {
      file_name = paste(output_prefix,"_foldchange_", geneofinterest[i], ".svg", sep="")
      svglite(file_name, width = 4, height = 6)
      print(plot_list[[i]])
      dev.off()
  }
}
```

#### 2.7 Shared analysis of more than one dataset

##### 2.7.1 Find shared genes, make euler plot

```
shared_euler <- function(deglist, output_prefix, diagram_type = "euler", pvalue_cutoff = 0.05, logfc_cutoff = 0.1, logfc_condition = "abs") { # condition can be "up", "down", or "abs"
  # List to store filtered gene sets
  filtered_deglist <- list()
  # Loop over each gene set in the list
  for (i in seq_along(deglist)) {
    gene_set <- deglist[[i]]
    # Determine the filter condition for log2FoldChange
    if (logfc_condition == "abs") {
      filtered_genes <- gene_set %>%
        filter(pvalue < pvalue_cutoff, abs(log2FoldChange) > logfc_cutoff) %>%
        pull(LLgeneAbbrev)
        filtered_deglist[[i]] <- filtered_genes
    } else if (logfc_condition == "up") {
      filtered_genes <- gene_set %>%
        filter(pvalue < pvalue_cutoff, log2FoldChange > logfc_cutoff) %>%
        pull(LLgeneAbbrev)
        filtered_deglist[[i]] <- filtered_genes
    } else if (logfc_condition == "down") {
      filtered_genes <- gene_set %>%
        filter(pvalue < pvalue_cutoff, log2FoldChange < logfc_cutoff) %>%
        pull(LLgeneAbbrev)
        filtered_deglist[[i]] <- filtered_genes
    } else {
      stop("Invalid logfc_condition. Choose 'abs', '>', or '<'.")
    }
  }
    # Filter by pvalue and log2FoldChange
  #  filtered_genes <- gene_set %>%
  #    filter(pvalue < pvalue_cutoff, abs(log2FoldChange) > logfc_cutoff) %>%
  #    pull(LLgeneAbbrev)
  #  # Store filtered genes in the list
  #  filtered_deglist[[i]] <- filtered_genes
  #}
  
  # Naming the sets for diagram
  names(filtered_deglist) <- names(deglist)
  combined_names <- paste(names(filtered_deglist), collapse = "_")
  
  # Compute pairwise intersections
  pairwise_combinations <- combn(names(filtered_deglist), 2, simplify = FALSE)
  pairwise_intersections <- sapply(pairwise_combinations, function(pair) {
    intersect(filtered_deglist[[pair[1]]], filtered_deglist[[pair[2]]])
  }, simplify = FALSE)

  # Create a data frame for pairwise intersections
  pairwise_intersections_df <- data.frame(
    Set1 = sapply(pairwise_combinations, `[`, 1),
    Set2 = sapply(pairwise_combinations, `[`, 2),
    Intersection = sapply(pairwise_intersections, paste, collapse = ", "), # Convert list to comma-separated string
    stringsAsFactors = FALSE
  )
  
  # Compute intersection of all sets
  all_sets_intersection <- Reduce(intersect, filtered_deglist)

  # Create a data frame for all sets intersection
  all_sets_intersection_df <- data.frame(
    Set1 = NA,  # Placeholder as there's no single pair for all sets intersection
    Set2 = NA,  # Placeholder
    Intersection = paste(all_sets_intersection, collapse = ", "), # Convert list to comma-separated string
    stringsAsFactors = FALSE
  )

  # Combine pairwise intersections and all sets intersection
  intersecting_data_df <- rbind(pairwise_intersections_df, all_sets_intersection_df)

  # Save intersecting data to CSV
  write.csv(intersecting_data_df, file = paste0(output_prefix, "intersecting_data.csv"), row.names = FALSE)

  if (length(filtered_deglist) > 0) {
    intersected_genes <- Reduce(intersect, filtered_deglist)
    # Creating a data frame to store the intersected genes (if needed)
    intersecting_data <- data.frame(Gene = intersected_genes, Set = rep(combined_names, length(intersected_genes)))
    write.csv(intersecting_data, file = paste0(output_prefix, combined_names, "_intersection.csv"), row.names = FALSE)
  } else {
    intersected_genes <- character(0)
  }
  # Create Venn or Euler diagram
  if (diagram_type == "venn") {
    png(filename = paste0(output_prefix, combined_names, "_venn_plot.png"), width = 1000, height = 1000)
    venn::venn(filtered_deglist)
    dev.off()
  } else if (diagram_type == "euler") {
    euler_data <- eulerr::euler(setNames(filtered_deglist, names(filtered_deglist)))
    print(euler_data)
    plot(euler_data, quantities=TRUE)
    #euler_plot <- ggplot2::ggplot(euler_data) + eulerr::geom_euler()
    #ggsave(filename = paste0(output_prefix, combined_names, "_euler_plot.png"), plot = euler_plot, width = 10, height = 10, units = "in")
    #print(euler_data)
    #png(filename = paste0(output_prefix, combined_names, "_euler_plot.png"), width = 1000, height = 1000)
    #eulerr::plot(euler_data)
    #euler::plot(euler_data)
    #euler::plot(eulerr::euler(filtered_deglist))
    #dev.off()
  } else {
    stop("Invalid diagram_type. Choose 'venn' or 'euler'.")
  }
}
```

##### 2.7.2 Bubble plot of shared genes

```
shared_bubble <-function(multiDEG, upgenes, downgenes){
  sharedgenes <- c(downgenes, upgenes)
  #filter to only include genes of interest
  bubbleDEG <- multiDEG %>% dplyr::filter(LLgeneAbbrev %in% sharedgenes)
  #find middle of foldchange (to center color scale)
  limit <- max(abs(bubbleDEG$log2FoldChange)) * c(-1, 1)
  #get color
  myheatcolors3 <- brewer.pal(name="RdBu", n=11)
  #order genes
  bubbleDEG$LLgeneAbbrev <- factor(bubbleDEG$LLgeneAbbrev, levels = sharedgenes)
  # create 'bubble plot' to summarize y signatures across x phenotypes
  bubbleplot_shared <- ggplot(bubbleDEG, aes(x=LLgeneAbbrev, y=mutation)) + 
    geom_point(aes(size=-log10(padj), color = log2FoldChange)) +
    scale_color_gradientn(colors = myheatcolors3, limit=limit) +
    theme_bw() +
    theme(axis.text.x = element_text(angle = 45, hjust = 1),
          axis.title.x = element_blank(),
          axis.title.y = element_blank())#+
    #annotate("rect", xmin = 0, xmax = 9.5, ymin = 0.25, ymax = 0.5, 
    #        alpha=1, fill="#472D7BFF") +
    #annotate("rect", xmin = 9.5, xmax = 33, ymin = 0.25, ymax = 0.5, 
    #        alpha=1, fill="#21908CFF")
  ggsave("shared_bubble.png", plot = bubbleplot_shared, width = 10, height = 7, dpi = 300)
  bubbleplot_shared 
}
```

##### 2.7.3 Scatterplot of shared genes

```
# FUNCTION NEEDS WORK - IT IS NOT COLORING WHAT IT SHOULD RIGHT NOW
deg_scatter <- function(multiDEG, genestolabel, chrnum1, chrnum2) {
  # Subset to only include comparison, logfc, gene name, and pvalue
  scatterdata <- multiDEG %>%
    dplyr::select(log2FoldChange, pvaluebound, chrom, mutation, LLgeneAbbrev) %>%
    dplyr::mutate(chrom = replace_na(chrom, "none")) %>%
    mutate(chromosome = ifelse(chrom == paste0("chr", chrnum1), paste0("chromosome ", chrnum1),
                               ifelse(chrom == paste0("chr", chrnum2), paste0("chromosome ", chrnum2), "other chromosome")))
  # Find duplicates
  duplicategenes <- dplyr::filter(scatterdata %>%
    distinct() %>%
    group_by(LLgeneAbbrev, mutation) %>%
    dplyr::count(), n != 1)$LLgeneAbbrev
  # Remove duplicate rows
  scatterdata <- scatterdata %>% dplyr::filter(!LLgeneAbbrev %in% duplicategenes)
  # Pivot wider to make untidy table
  scatterdata <- scatterdata %>%
    distinct() %>%
    pivot_wider(names_from = mutation, values_from = c(log2FoldChange, pvaluebound), values_fill = NA)
  # Extract column names for log2FoldChange and pvaluebound
  log2FoldChange_cols <- grep("^log2FoldChange_", names(scatterdata), value = TRUE)
  pvaluebound_cols <- grep("^pvaluebound_", names(scatterdata), value = TRUE)
  # Ensure we have at least two columns for plotting
  if (length(log2FoldChange_cols) < 2) {
    stop("Not enough log2FoldChange columns to plot.")
  }
  # Extract mutation names for labels
  mutation_names <- sub("^log2FoldChange_", "", log2FoldChange_cols)
  x_label <- paste("log2 fold change", mutation_names[1])
  y_label <- paste("log2 fold change", mutation_names[2])
  # Rename columns for easier plotting (using the first two as examples)
  scatterdata <- scatterdata %>%
    dplyr::rename(
      log2FoldChange_x = !!log2FoldChange_cols[1],
      log2FoldChange_y = !!log2FoldChange_cols[2]
    )
  # Add row names
  row.names(scatterdata) <- scatterdata$LLgeneAbbrev
  # Subset to only include labeled genes
  myTopHits.labels <- scatterdata %>% dplyr::filter(LLgeneAbbrev %in% genestolabel)
  # Change order that points are plotted
  scatterdata <- scatterdata %>%
    arrange(match(chromosome, c("other chromosome", paste0("chromosome ", chrnum1), paste0("chromosome ", chrnum2))), desc(chromosome))
  # Plotting
  scatterplot <- ggplot() +
    geom_point(data = scatterdata, aes(x = log2FoldChange_x, y = log2FoldChange_y, color = chromosome, text = paste("Symbol:", LLgeneAbbrev))) +
    geom_hline(yintercept = 0, linetype = 'dotted') +
    geom_vline(xintercept = 0, linetype = 'dotted') +
    theme_bw() +
    scale_color_manual(values = c("darkgrey", "#AADC32FF", "#27AD81FF", "#472D7BFF", "darkgrey"), name = "chromosome") +
    geom_label_repel(data = myTopHits.labels, aes(x = log2FoldChange_x, y = log2FoldChange_y, label = LLgeneAbbrev), force = 1, nudge_y = .5, size = 2.5, max.overlaps = Inf, show.legend = FALSE, color = "black") +
    ylab(paste0("log2 fold change ",y_label)) +
    xlab(paste0("log2 fold change ",x_label))

  filename <- paste0("scatterplot_", mutation_names[1], "_vs_", mutation_names[2], ".png")
  ggsave(filename, plot = scatterplot, width = 10, height = 7, dpi = 300)
}
```

### 3 Running zone

#### 3.1 Processing area, this is where we input the specific data etc

##### 3.1.1 First steps and GSEAs for ebf3a hom vs hetandwt

```
# User input for specific chromosome where gene is located and names of data
refseq_genes = "/Users/sthyme/ebf3_rna_analysis/ncbi_refseqgenes"
chrnum = 12
output_prefix0 = "ebf3a-2dpf-homvshetandwt_"
deg_file = "ebf3a-2dpf-homvshetandwt_allresults_wt_hom-with-normalized.csv"
csv = read.csv("ebf3a-2dpf-homvshetandwt_normalized_reads_gene_list.csv") #[,2:10] # IMPORTANT, THIS 8 MUST CHANGE DEPENDING ON HOW MANY SAMPLES
genecounts0 <- csv[, 2:ncol(csv)]
mutgene = "ebf3a"
# add chromosome information
DEG0 <- process_DEGs(deg_file, refseq_genes, output_prefix0, chrnum)
# make manhattan plot and gsea to check if DEGs are on same chromosome as gene itself
# don't need now
#manhattan_plot(DEG0, output_prefix0, chrnum) # one of these two plots is not always working as expected, although it has (the one that is a Zoom of just a few genes)
#chr_gsea(output_prefix0, refseq_genes, DEG0, chrnum, mutgene)
# deg analysis
GO_df <- go_analysis(output_prefix0, DEG0, chrnum)
c5gsea0 <- c5_gsea(DEG0, output_prefix0, chrnum)
scGSEA_danio(output_prefix0, DEG0, chrnum) # can change the last argument to do any of the daniocell clusters or leave out to do all (but heatmap is hard to read)
scGSEA_danio(output_prefix0, DEG0, chrnum, "hema") # can change the last argument to do any of the daniocell clusters or leave out to do all (but heatmap is hard to read)
scGSEA_danio(output_prefix0, DEG0, chrnum, "neur") # can change the last argument to do any of the daniocell clusters or leave out to do all (but heatmap is hard to read)
scGSEA_danio(output_prefix0, DEG0, chrnum, "eye") # can change the last argument to do any of the daniocell clusters or leave out to do all (but heatmap is hard to read)
scGSEA_danio(output_prefix0, DEG0, chrnum, "iono") # can change the last argument to do any of the daniocell clusters or leave out to do all (but heatmap is hard to read)
scGSEA_danio(output_prefix0, DEG0, chrnum, "otic") # can change the last argument to do any of the daniocell clusters or leave out to do all (but heatmap is hard to read)
scGSEA_danio(output_prefix0, DEG0, chrnum, "glia") # can change the last argument to do any of the daniocell clusters or leave out to do all (but heatmap is hard to read)
scGSEA_danio(output_prefix0, DEG0, chrnum, "tast") # can change the last argument to do any of the daniocell clusters or leave out to do all (but heatmap is hard to read)
scGSEA(output_prefix0, DEG0, chrnum)
#anatGSEA(output_prefix0, DEG0, chrnum) # keeps just giving epiphysis and pineal complex when it's not working well, so skip
```

##### 3.1.2 User input for specific genes and go terms etc for ebf3a hom vs hetandwt

```
# List of terms you are interested in
terms_of_interest <- c("GO:0002088", "GO:0007043", "GO:0006869", "GO:0016050", "GO:0048592")  # Replace with your actual GO terms
genestolabel <- c("ebf3a","neurod1", "lhx1a", "npffl", "gnrh3", "pcdh8","scn1lab")
# Extract genes for the specified terms
genes_for_terms <- extract_genes_for_terms(GO_df, terms_of_interest)
# add custom terms
genes_for_terms[["known target genes"]] <- c("pcdh8", "neurod1", "prph")
genes_for_terms[["genes of interest"]] <- c("npffl", "gnrh3", "scn1lab", "lhx1a")

color_vector = c("#AADC32FF", "#27AD81FF", "#472D7BFF", "gold", "deepskyblue", "orange", 
                                  "darkgrey", "#D697FF", "lightblue", "darkred")
create_volcano_plot(output_prefix0, DEG0, genes_for_terms, genestolabel, chrnum, color_vector)#, 0.01, 1.0, 3.5, 3.7, 0.1, TRUE)

# can include if you want, but code won't fail if you just want to plot all instead
networkterms_nucleic <- c(
  "GOBP_NCRNA_METABOLIC_PROCESS",
  "GOMF_CATALYTIC_ACTIVITY_ACTING_ON_A_NUCLEIC_ACID",
  "GOBP_NCRNA_PROCESSING",
  "GOBP_RIBONUCLEOPROTEIN_COMPLEX_BIOGENESIS",
  "GOBP_RIBOSOME_BIOGENESIS",
  "GOMF_CATALYTIC_ACTIVITY_ACTING_ON_RNA",
  "GOCC_PEPTIDASE_COMPLEX",
  "GOCC_PRERIBOSOME",
  "GOBP_RRNA_METABOLIC_PROCESS",
  "GOCC_NUCLEOID",
  "GOCC_RIBOSOMAL_SUBUNIT",
  "GOCC_CATALYTIC_STEP_2_SPLICEOSOME")
gsea_terms = list()
#gsea_terms <- list(nucleic = networkterms_nucleic)
gsea_plot(output_prefix0, gsea_terms, c5gsea0, DEG0, chrnum)

mySelectedGenesa <- unique(c("ebf3a", "pcdh8", "npffl", "gnrh3", "arxa", "neurod1", "prph", "vsig8a", "gpc1a", "gpc3", "dlgap2a", "cdc25d", "fam69ab", "irx4b", "dgkzb", "prkcg", "ppargc1a", "osbp2", "rnf152", "cadm2b", "pvalb8", "ifngr1", "krt1-c5", "gsnb", "moxd1l", "cd109"))
#gene_heatmap(output_prefix0, genecounts0, mySelectedGenesa)
controlave = c(6,7) # column ids for wt (it is by one from the excel file because we select columns 2-end rather than including column 1 when read in)
gene_barplots(output_prefix0, genecounts0, mySelectedGenesa, controlave)
```

##### 3.1.3 First steps and GSEAs for ebf3a hom vs wt

```
# User input for specific chromosome where gene is located and names of data
refseq_genes = "/Users/sthyme/ebf3_rna_analysis/ncbi_refseqgenes"
chrnum = 12
output_prefix1 = "ebf3a-2dpf-homvswt_"
deg_file = "ebf3a-2dpf-homvswt_allresults_wt_hom-with-normalized.csv"
csv = read.csv("ebf3a-2dpf-homvswt_normalized_reads_gene_list.csv")#[,2:8] # IMPORTANT, THIS 8 MUST CHANGE DEPENDING ON HOW MANY SAMPLES
genecounts1 <- csv[, 2:ncol(csv)]
mutgene = "ebf3a"
# add chromosome information
DEG1 <- process_DEGs(deg_file, refseq_genes, output_prefix1, chrnum)
# make manhattan plot and gsea to check if DEGs are on same chromosome as gene itself
# don't need now
#manhattan_plot(DEG1, output_prefix1, chrnum)
#chr_gsea(output_prefix1, refseq_genes, DEG1, chrnum, mutgene)
# deg analysis
GO_df <- go_analysis(output_prefix1, DEG1, chrnum)
c5gsea1 <- c5_gsea(DEG1, output_prefix1, chrnum)
# full heatmap is failing for some reason, there must be an issue in one of the columns, but the "neur" only one is fine
#scGSEA_danio(output_prefix1, DEG1, chrnum) # can change the last argument to do any of the daniocell clusters or leave out to do all (but heatmap is hard to read)
scGSEA_danio(output_prefix1, DEG1, chrnum, "neur") # can change the last argument to do any of the daniocell clusters or leave out to do all (but heatmap is hard to read)
scGSEA(output_prefix1, DEG1, chrnum)
#anatGSEA(output_prefix1, DEG1, chrnum) # keeps just giving epiphysis and pineal complex
```

##### 3.1.4 User input for specific genes and go terms etc for ebf3a hom vs wt

```
# List of terms you are interested in
terms_of_interest <- c("GO:0002088", "GO:0007043", "GO:0061564", "GO:0048592")  # Replace with your actual GO terms
genestolabel <- c("ebf3a","neurod1", "lhx1a", "npffl", "gnrh3", "pcdh8","scn1lab")
# Extract genes for the specified terms
genes_for_terms <- extract_genes_for_terms(GO_df, terms_of_interest)
# add custom terms
genes_for_terms[["known target genes"]] <- c("pcdh8", "neurod1", "prph")
genes_for_terms[["genes of interest"]] <- c("npffl", "gnrh3", "scn1lab", "lhx1a")

color_vector = c("#AADC32FF", "#27AD81FF", "#472D7BFF", "gold", "deepskyblue", "orange", 
                                  "darkgrey", "#D697FF", "lightblue", "darkred")
create_volcano_plot(output_prefix1, DEG1, genes_for_terms, genestolabel, chrnum, color_vector)#, 0.01, 1.0, 3.5, 3.7, 0.1, TRUE)

# can include if you want, but code won't fail if you just want to plot all instead
networkterms_placeholder <- c(
  "GOBP_SENSORY_PERCEPTION_OF_LIGHT_STIMULUS",
  "GOBP_SENSORY_PERCEPTION",
  "HP_ABNORMAL_VISUAL_ELECTROPHYSIOLOGY",
  "HP_ABNORMAL_DARK_ADAPTED_ELECTRORETINOGRAM",
  "GOCC_PROTEASOME_COMPLEX",
  "GOMF_CATALYTIC_ACTIVITY_ACTING_ON_RNA",
  "HP_ABNORMALITY_IRIS_MORPHOLOGY",
  "GOCC_DESMOSOME",
  "GOCC_INNER_MITOCHONDRIAL_MEMBRANE_PROTEIN_COMPLEX")

gsea_terms = list()
#gsea_terms <- list(placeholder = networkterms_placeholder)
gsea_plot(output_prefix1, gsea_terms, c5gsea1, DEG1, chrnum)

#mySelectedGenesa <- unique(c("ebf3a", "pcdh8", "npffl", "gnrh3", "arxa", "neurod1", "prph", "vsig8a", "gpc1a", "gpc3", "dlgap2a", "cdc25d", "fam69ab", "irx4b", "dgkzb", "prkcg", "ppargc1a", "osbp2", "rnf152", "cadm2b", "pvalb8", "ifngr1", "krt1-c5", "gsnb", "moxd1l", "cd109"))
#gene_heatmap(output_prefix0, genecounts0, mySelectedGenesa)
#controlave = c(6,7) # column ids for wt (it is by one from the excel file because we select columns 2-end rather than including column 1 when read in)
#gene_barplots(output_prefix0, genecounts0, mySelectedGenesa, controlave)
```

##### 3.1.5 First steps and GSEAs for ebf3a het vs wt

```
# User input for specific chromosome where gene is located and names of data
refseq_genes = "/Users/sthyme/ebf3_rna_analysis/ncbi_refseqgenes"
chrnum = 12
output_prefix2 = "ebf3a-2dpf-hetvswt_"
deg_file = "ebf3a-2dpf-hetvswt_allresults_wt_hom-with-normalized.csv"
csv = read.csv("ebf3a-2dpf-hetvswt_normalized_reads_gene_list.csv")#[,2:8] # IMPORTANT, THIS 8 MUST CHANGE DEPENDING ON HOW MANY SAMPLES
genecounts2 <- csv[, 2:ncol(csv)]
mutgene = "ebf3a"
# add chromosome information
DEG2 <- process_DEGs(deg_file, refseq_genes, output_prefix2, chrnum)
# make manhattan plot and gsea to check if DEGs are on same chromosome as gene itself
# don't need now
#manhattan_plot(DEG2, output_prefix2, chrnum)
#chr_gsea(output_prefix2, refseq_genes, DEG2, chrnum, mutgene)
# deg analysis
GO_df <- go_analysis(output_prefix2, DEG2, chrnum)
c5gsea2 <- c5_gsea(DEG2, output_prefix2, chrnum)
scGSEA_danio(output_prefix2, DEG2, chrnum) # can change the last argument to do any of the daniocell clusters or leave out to do all (but heatmap is hard to read)
scGSEA_danio(output_prefix2, DEG2, chrnum, "neur") # can change the last argument to do any of the daniocell clusters or leave out to do all (but heatmap is hard to read)
scGSEA(output_prefix2, DEG2, chrnum)
#anatGSEA(output_prefix2, DEG2, chrnum) # keeps just giving epiphysis and pineal complex
```

##### 3.1.6 User input for specific genes and go terms etc for ebf3a het vs wt

```
# List of terms you are interested in
terms_of_interest <- c("GO:0002088", "GO:0006260", "GO:0061564", "GO:0048592", "GO:0030900", "GO:0048512", "GO:0048915")  # Replace with your actual GO terms
genestolabel <- c("ebf3a","neurod1", "lhx1a", "npffl", "gnrh3", "pcdh8","scn1lab", "arxa")
# Extract genes for the specified terms
genes_for_terms <- extract_genes_for_terms(GO_df, terms_of_interest)
# add custom terms
genes_for_terms[["known target genes"]] <- c("pcdh8", "neurod1", "prph")
genes_for_terms[["genes of interest"]] <- c("npffl", "gnrh3", "scn1lab", "lhx1a", "arxa")

color_vector = c("#AADC32FF", "#27AD81FF", "#472D7BFF", "gold", "deepskyblue", "orange", 
                                  "#D697FF", "lightblue", "darkgrey", "darkred")
create_volcano_plot(output_prefix2, DEG2, genes_for_terms, genestolabel, chrnum, color_vector, 0.01, 0.7, 1.5, 1.7, 0.1, TRUE)

networkterms_placeholder <- c(
  "GOBP_CELL_CYCLE_DNA_REPLICATION",
  "GOCC_SYNAPTIC_MEMBRANE",
  "GOCC_ION_CHANNEL_COMPLEX",
  "GOCC_CATION_CHANNEL_COMPLEX",
  "GOBP_ADULT_BEHAVIOR",
  "GOMF_CATALYTIC_ACTIVITY_ACTING_ON_RNA",
  "GOCC_NEURON_TO_NEURON_SYNAPSE",
  "GOMF_KINASE_BINDING",
  "GOCC_DESMOSOME")

gsea_terms = list()
#gsea_terms <- list(placeholder = networkterms_placeholder)
# was failing for het vs wt because there were too many terms allowed - the package has a limit of 200
gsea_plot(output_prefix2, gsea_terms, c5gsea2, DEG2, chrnum)

#mySelectedGenesa <- unique(c("ebf3a", "pcdh8", "npffl", "gnrh3", "arxa", "neurod1", "prph", "vsig8a", "gpc1a", "gpc3", "dlgap2a", "cdc25d", "fam69ab", "irx4b", "dgkzb", "prkcg", "ppargc1a", "osbp2", "rnf152", "cadm2b", "pvalb8", "ifngr1", "krt1-c5", "gsnb", "moxd1l", "cd109"))
#gene_heatmap(output_prefix0, genecounts0, mySelectedGenesa)
#controlave = c(6,7) # column ids for wt (it is by one from the excel file because we select columns 2-end rather than including column 1 when read in)
#gene_barplots(output_prefix0, genecounts0, mySelectedGenesa, controlave)
```

##### 3.1.7 First steps and GSEAs for ebf3a hom vs het

```
# User input for specific chromosome where gene is located and names of data
refseq_genes = "/Users/sthyme/ebf3_rna_analysis/ncbi_refseqgenes"
chrnum = 12
output_prefix3 = "ebf3a-2dpf-homvshet_"
deg_file = "ebf3a-2dpf-homvshet_allresults_wt_hom-with-normalized.csv"
csv = read.csv("ebf3a-2dpf-homvshet_normalized_reads_gene_list.csv")
genecounts3 <- csv[, 2:ncol(csv)]
mutgene = "ebf3a"
# add chromosome information
DEG3 <- process_DEGs(deg_file, refseq_genes, output_prefix3, chrnum)
# make manhattan plot and gsea to check if DEGs are on same chromosome as gene itself
# don't need now
#manhattan_plot(DEG3, output_prefix3, chrnum)
#chr_gsea(output_prefix3, refseq_genes, DEG3, chrnum, mutgene)
# deg analysis
GO_df <- go_analysis(output_prefix3, DEG3, chrnum)
c5gsea3 <- c5_gsea(DEG3, output_prefix3, chrnum)
# heatmaps can be buggy sometimes
scGSEA_danio(output_prefix3, DEG3, chrnum) # can change the last argument to do any of the daniocell clusters or leave out to do all (but heatmap is hard to read)
#scGSEA(output_prefix3, DEG3, chrnum)
#anatGSEA(output_prefix3, DEG3, chrnum) # keeps just giving epiphysis and pineal complex
# skipping the volcano and GSEA term plots for hom vs het, which would be below. It's redundant with hom vs wt and hom vs hetandwt
# it does give some interesting clusters when running Bushra's scGSEA with 5 dpf head clusters, but I think it is due to misregulation of pathways in the het, or they would show up in hom vs wt
```

##### 3.1.8 First steps and GSEAs for 5 dpf ebf3a hom vs wt

```
# User input for specific chromosome where gene is located and names of data
refseq_genes = "/Users/sthyme/ebf3_rna_analysis/ncbi_refseqgenes"
chrnum = 12
output_prefix4 = "ebf3a-5dpf-homvswt_"
deg_file = "ebf3a-5dpf-homvswt_allresults_wt_hom-with-normalized.csv"
genecounts4 = read.csv("ebf3a-5dpf-homvswt_normalized_reads_gene_list.csv")[,2:8] # IMPORTANT, THIS 8 MUST CHANGE DEPENDING ON HOW MANY SAMPLES
mutgene = "ebf3a"
# add chromosome information
DEG4 <- process_DEGs(deg_file, refseq_genes, output_prefix4, chrnum)
# make manhattan plot and gsea to check if DEGs are on same chromosome as gene itself
# don't need now
#manhattan_plot(DEG4, output_prefix4, chrnum)
#chr_gsea(output_prefix4, refseq_genes, DEG4, chrnum, mutgene)
# deg analysis
GO_df <- go_analysis(output_prefix4, DEG4, chrnum)
c5gsea4 <- c5_gsea(DEG4, output_prefix4, chrnum)
# heatmaps can be buggy sometimes
scGSEA_danio(output_prefix4, DEG4, chrnum) # can change the last argument to do any of the daniocell clusters or leave out to do all (but heatmap is hard to read)
scGSEA(output_prefix4, DEG4, chrnum)
#anatGSEA(output_prefix4, DEG2, chrnum) # keeps just giving epiphysis and pineal complex
```

##### 3.1.9 User input for specific genes and go terms etc for 5 dpf ebf3a hom vs wt

```
# List of terms you are interested in
#terms_of_interest <- c("GO:0002088", "GO:0006260", "GO:0061564", "GO:0048592", "GO:0030900", "GO:0048512", "GO:0048915")  # Replace with your actual GO terms
genestolabel <- c("ebf3a","npffl", "gnrh3", "pcdh8", "cbln18", "prkcg", "ca8", "aldoca", "pif1")
# Extract genes for the specified terms
genes_for_terms = list()
#genes_for_terms <- extract_genes_for_terms(GO_df, terms_of_interest)
# add custom terms
genes_for_terms[["known target genes"]] <- c("pcdh8", "neurod1", "prph")
genes_for_terms[["genes of interest"]] <- c("ebf3a", "npffl", "gnrh3")
genes_for_terms[["lateral line"]] <- c("cbln18", "cbln20", "si:dkeyp-110c7.4", "si:dkey-33m11.8")
genes_for_terms[["cerebellum Purkinje cells"]] <- c("aldoca", "ca8", "grid2ipa", "prkcg")

color_vector = c("#AADC32FF", "#27AD81FF", "#472D7BFF", "gold", "darkgrey", "deepskyblue", "orange", 
                                  "#D697FF", "lightblue", "darkred")
create_volcano_plot(output_prefix4, DEG4, genes_for_terms, genestolabel, chrnum, color_vector, 0.0015, 1.0, 5.5, 5.7, 0.1, FALSE)

#gsea_terms <- list(placeholder = networkterms_placeholder)
#gsea_plot(output_prefix4, gsea_terms, c5gsea2, DEG4, chrnum)

mySelectedGenesll<-c("cbln18", "cbln20", "si:dkeyp-110c7.4", "si:dkey-33m11.8")
mySelectedGenescb<-c("aldoca", "ca8", "grid2ipa", "prkcg")
mySelectedGenes<-c("npffl", "gnrh3")
mySelectedGenesall<-c("cbln18", "cbln20", "si:dkeyp-110c7.4", "si:dkey-33m11.8","aldoca", "ca8", "grid2ipa", "prkcg","npffl", "gnrh3")
#mySelectedGenesa <- unique(c("ebf3a", "pcdh8", "npffl", "gnrh3", "arxa", "neurod1", "prph", "vsig8a", "gpc1a", "gpc3", "dlgap2a", "cdc25d", "fam69ab", "irx4b", "dgkzb", "prkcg", "ppargc1a", "osbp2", "rnf152", "cadm2b", "pvalb8", "ifngr1", "krt1-c5", "gsnb", "moxd1l", "cd109"))
gene_heatmap(paste0(output_prefix4,"_lateralline_"), genecounts4, mySelectedGenesll)
```

```
## quartz_off_screen 
##                 2
```

```
gene_heatmap(paste0(output_prefix4,"_cerebellumpurkinje_"), genecounts4, mySelectedGenescb)
```

```
## quartz_off_screen 
##                 2
```

```
gene_heatmap(paste0(output_prefix4,"_npfflandgnrh3_"), genecounts4, mySelectedGenes)
```

```
## quartz_off_screen 
##                 2
```

```
gene_heatmap(paste0(output_prefix4,"_all_"), genecounts4, mySelectedGenesall)
```

```
## quartz_off_screen 
##                 2
```

```
controlave = c(6,7) # column ids for wt (it is by one from the excel file because we select columns 2-end rather than including column 1 when read in)
gene_barplots(paste0(output_prefix4,"_lateralline_"), genecounts4, mySelectedGenesll, controlave)
gene_barplots(paste0(output_prefix4,"_cerebellumpurkinje_"), genecounts4, mySelectedGenescb, controlave)
gene_barplots(paste0(output_prefix4,"_npfflandgnrh3_"), genecounts4, mySelectedGenes, controlave)
```

##### 3.1.10 Section for working with multiple genes together

```
# This section is for working with multiple gene sets together
#add columns to old and new data about whether data is old or new
DEG1$mutation <- "homvswt-2dpf"
DEG2$mutation <- "hetvswt-dpf"
DEG3$mutation <- "homvshet-dpf"
#combine two mutations
multiDEG <- rbind(DEG1, DEG2, DEG3)
deglist <- list(homvswt = DEG1, hetvswt = DEG2, homvshet = DEG3)
shared_euler(deglist, "ebf3a_allcombos_upgenes", diagram_type = "euler", 0.05, 0.2, "up")
```

```
##                          original  fitted residuals regionError
## homvswt                       495 494.796     0.204       0.002
## hetvswt                       348 347.764     0.236       0.001
## homvshet                      217 216.587     0.413       0.001
## homvswt&hetvswt               156 157.003    -1.003       0.000
## homvswt&homvshet              143 144.110    -1.110       0.000
## hetvswt&homvshet                0   8.602    -8.602       0.006
## homvswt&hetvswt&homvshet       34  30.017     3.983       0.003
## 
## diagError: 0.006 
## stress:    0
```

```
shared_euler(deglist, "ebf3a_allcombos_downgenes",diagram_type = "euler", 0.05, -0.2, "down")
```

```
##                          original  fitted residuals regionError
## homvswt                       488 487.999     0.001           0
## hetvswt                       370 369.999     0.001           0
## homvshet                      242 241.998     0.002           0
## homvswt&hetvswt               162 162.006    -0.006           0
## homvswt&homvshet              159 159.007    -0.007           0
## hetvswt&homvshet                0   0.390    -0.390           0
## homvswt&hetvswt&homvshet        8   7.936     0.064           0
## 
## diagError: 0 
## stress:    0
```

```
shared_euler(deglist, "ebf3a_allcombos_allgenes",diagram_type = "euler", 0.05, -0.2, "abs")
```

```
##                          original   fitted residuals regionError
## homvswt                      1104 1108.300    -4.300       0.022
## hetvswt                       769  773.909    -4.909       0.016
## homvshet                      262  277.336   -15.336       0.010
## homvswt&hetvswt               468  450.543    17.457       0.003
## homvswt&homvshet              339  313.852    25.148       0.002
## hetvswt&homvshet              234   13.750   220.250       0.068
## homvswt&hetvswt&homvshet       59  112.182   -53.182       0.019
## 
## diagError: 0.068 
## stress:    0.023
```

```
DEG4$mutation <- "homvswt-5dpf"
multiDEGb <- rbind(DEG1, DEG4)
deglistb <- list(homvswt2 = DEG1, homvswt5 = DEG4)
shared_euler(deglistb, "ebf3a_5dpfvs2dpf_allgenes", diagram_type = "euler", 0.05, 0.2, "abs")
```

```
##                   original fitted residuals regionError
## homvswt2              1611   1611         0           0
## homvswt5               180    180         0           0
## homvswt2&homvswt5       34     34         0           0
## 
## diagError: 0 
## stress:    0
```

```
upgenes = c("pgs1", "pif1")
downgenes = c("ebf3a", "krt91", "cyt1l", "pfn1", "aep1","krt4", "cyt1", "krt5", "pvalb8", "npffl", "icn2", "hrc", "gnrh3", "ponzr3", "fosb", "mid1ip1a", "fut9d")
# FUTURE: add arguments for shape, since the look of this plot is impacted by the designated shape
shared_bubble(multiDEGb, upgenes, downgenes)
```

```
# this plot is not refined or checked over
# only leaving this here for now as a placeholder, since the code works
genestolabel <- unique(c("ebf3a", "krt91", "cyt1l", "pfn1", "aep1","krt4", "cyt1", "krt5", "pvalb8", "npffl", "icn2", "hrc", "gnrh3", "ponzr3", "fosb", "mid1ip1a", "fut9d"))
chrnum1 = 12
chrnum2 = 12
# chrnum1 and 2 should be in the order as the degs are bound
deg_scatter(multiDEGb, genestolabel, chrnum1, chrnum2)
```

### 4 Session info

Packages and versions necessary to reproduce the results in this
report.

```
sessionInfo()
```

```
## R version 4.4.1 (2024-06-14)
## Platform: x86_64-apple-darwin20
## Running under: macOS Sonoma 14.1
## 
## Matrix products: default
## BLAS:   /Library/Frameworks/R.framework/Versions/4.4-x86_64/Resources/lib/libRblas.0.dylib 
## LAPACK: /Library/Frameworks/R.framework/Versions/4.4-x86_64/Resources/lib/libRlapack.dylib;  LAPACK version 3.12.0
## 
## locale:
## [1] en_US.UTF-8/en_US.UTF-8/en_US.UTF-8/C/en_US.UTF-8/en_US.UTF-8
## 
## time zone: America/New_York
## tzcode source: internal
## 
## attached base packages:
## [1] stats4    stats     graphics  grDevices utils     datasets  methods  
## [8] base     
## 
## other attached packages:
##  [1] eulerr_7.0.2           venn_1.12              gplots_3.1.3.1        
##  [4] svglite_2.1.3          ggraph_2.2.1           colorspace_2.1-1      
##  [7] ggpubr_0.6.0           lattice_0.22-6         plotly_4.10.4         
## [10] BaseSet_0.9.0          ontologyIndex_2.12     enrichplot_1.24.2     
## [13] msigdbr_7.5.1          org.Dr.eg.db_3.19.1    clusterProfiler_4.12.2
## [16] gprofiler2_0.2.3       GSVA_1.52.3            GSEABase_1.66.0       
## [19] graph_1.82.0           annotate_1.82.0        XML_3.99-0.17         
## [22] AnnotationDbi_1.66.0   IRanges_2.38.1         S4Vectors_0.42.1      
## [25] Biobase_2.64.0         BiocGenerics_0.50.0    DT_0.33               
## [28] RColorBrewer_1.1-3     ggrepel_0.9.5          scales_1.3.0          
## [31] viridis_0.6.5          viridisLite_0.4.2      cowplot_1.1.3         
## [34] lubridate_1.9.3        forcats_1.0.0          stringr_1.5.1         
## [37] dplyr_1.1.4            purrr_1.0.2            readr_2.1.5           
## [40] tidyr_1.3.1            tibble_3.2.1           ggplot2_3.5.1         
## [43] tidyverse_2.0.0        knitr_1.48             tinytex_0.52          
## [46] rmarkdown_2.27        
## 
## loaded via a namespace (and not attached):
##   [1] splines_4.4.1               bitops_1.0-8               
##   [3] ggplotify_0.1.2             polyclip_1.10-7            
##   [5] lifecycle_1.0.4             rstatix_0.7.2              
##   [7] vroom_1.6.5                 MASS_7.3-61                
##   [9] crosstalk_1.2.1             backports_1.5.0            
##  [11] magrittr_2.0.3              sass_0.4.9                 
##  [13] jquerylib_0.1.4             yaml_2.3.10                
##  [15] DBI_1.2.3                   abind_1.4-5                
##  [17] zlibbioc_1.50.0             GenomicRanges_1.56.1       
##  [19] yulab.utils_0.1.5           tweenr_2.0.3               
##  [21] GenomeInfoDbData_1.2.12     irlba_2.3.5.1              
##  [23] tidytree_0.4.6              codetools_0.2-20           
##  [25] DelayedArray_0.30.1         DOSE_3.30.2                
##  [27] ggforce_0.4.2               tidyselect_1.2.1           
##  [29] aplot_0.2.3                 UCSC.utils_1.0.0           
##  [31] farver_2.1.2                ScaledMatrix_1.12.0        
##  [33] matrixStats_1.3.0           jsonlite_1.8.8             
##  [35] tidygraph_1.3.1             systemfonts_1.1.0          
##  [37] polylabelr_0.2.0            ggnewscale_0.5.0           
##  [39] tools_4.4.1                 ragg_1.3.2                 
##  [41] treeio_1.28.0               Rcpp_1.0.13                
##  [43] glue_1.7.0                  gridExtra_2.3              
##  [45] SparseArray_1.4.8           admisc_0.35                
##  [47] xfun_0.46                   qvalue_2.36.0              
##  [49] MatrixGenerics_1.16.0       GenomeInfoDb_1.40.1        
##  [51] HDF5Array_1.32.0            withr_3.0.1                
##  [53] fastmap_1.2.0               rhdf5filters_1.16.0        
##  [55] fansi_1.0.6                 caTools_1.18.2             
##  [57] digest_0.6.36               rsvd_1.0.5                 
##  [59] timechange_0.3.0            R6_2.5.1                   
##  [61] gridGraphics_0.5-1          textshaping_0.4.0          
##  [63] GO.db_3.19.1                gtools_3.9.5               
##  [65] RSQLite_2.3.7               utf8_1.2.4                 
##  [67] generics_0.1.3              data.table_1.15.4          
##  [69] graphlayouts_1.1.1          httr_1.4.7                 
##  [71] htmlwidgets_1.6.4           S4Arrays_1.4.1             
##  [73] scatterpie_0.2.3            pkgconfig_2.0.3            
##  [75] gtable_0.3.5                blob_1.2.4                 
##  [77] SingleCellExperiment_1.26.0 XVector_0.44.0             
##  [79] shadowtext_0.1.4            htmltools_0.5.8.1          
##  [81] carData_3.0-5               fgsea_1.30.0               
##  [83] png_0.1-8                   SpatialExperiment_1.14.0   
##  [85] ggfun_0.1.5                 rstudioapi_0.16.0          
##  [87] tzdb_0.4.0                  reshape2_1.4.4             
##  [89] rjson_0.2.21                nlme_3.1-165               
##  [91] cachem_1.1.0                rhdf5_2.48.0               
##  [93] KernSmooth_2.23-24          parallel_4.4.1             
##  [95] HDO.db_0.99.1               pillar_1.9.0               
##  [97] grid_4.4.1                  vctrs_0.6.5                
##  [99] car_3.1-2                   BiocSingular_1.20.0        
## [101] beachmat_2.20.0             xtable_1.8-4               
## [103] evaluate_0.24.0             magick_2.8.4               
## [105] cli_3.6.3                   compiler_4.4.1             
## [107] rlang_1.1.4                 crayon_1.5.3               
## [109] ggsignif_0.6.4              labeling_0.4.3             
## [111] plyr_1.8.9                  fs_1.6.4                   
## [113] stringi_1.8.4               BiocParallel_1.38.0        
## [115] babelgene_22.9              munsell_0.5.1              
## [117] Biostrings_2.72.1           lazyeval_0.2.2             
## [119] GOSemSim_2.30.0             Matrix_1.7-0               
## [121] hms_1.1.3                   patchwork_1.2.0            
## [123] sparseMatrixStats_1.16.0    bit64_4.0.5                
## [125] Rhdf5lib_1.26.0             KEGGREST_1.44.1            
## [127] highr_0.11                  SummarizedExperiment_1.34.0
## [129] broom_1.0.6                 igraph_2.0.3               
## [131] memoise_2.0.1               bslib_0.8.0                
## [133] ggtree_3.12.0               fastmatch_1.1-4            
## [135] bit_4.0.5                   ape_5.8                    
## [137] gson_0.1.0
```
