## Supplementary figures and images for "Disrupted development of sensory systems and the cerebellum in a zebrafish *ebf3a* mutant"

### ebf3a-2dpf-homvshetandwt_eye_scRNA-danio-byp_heatmap.png

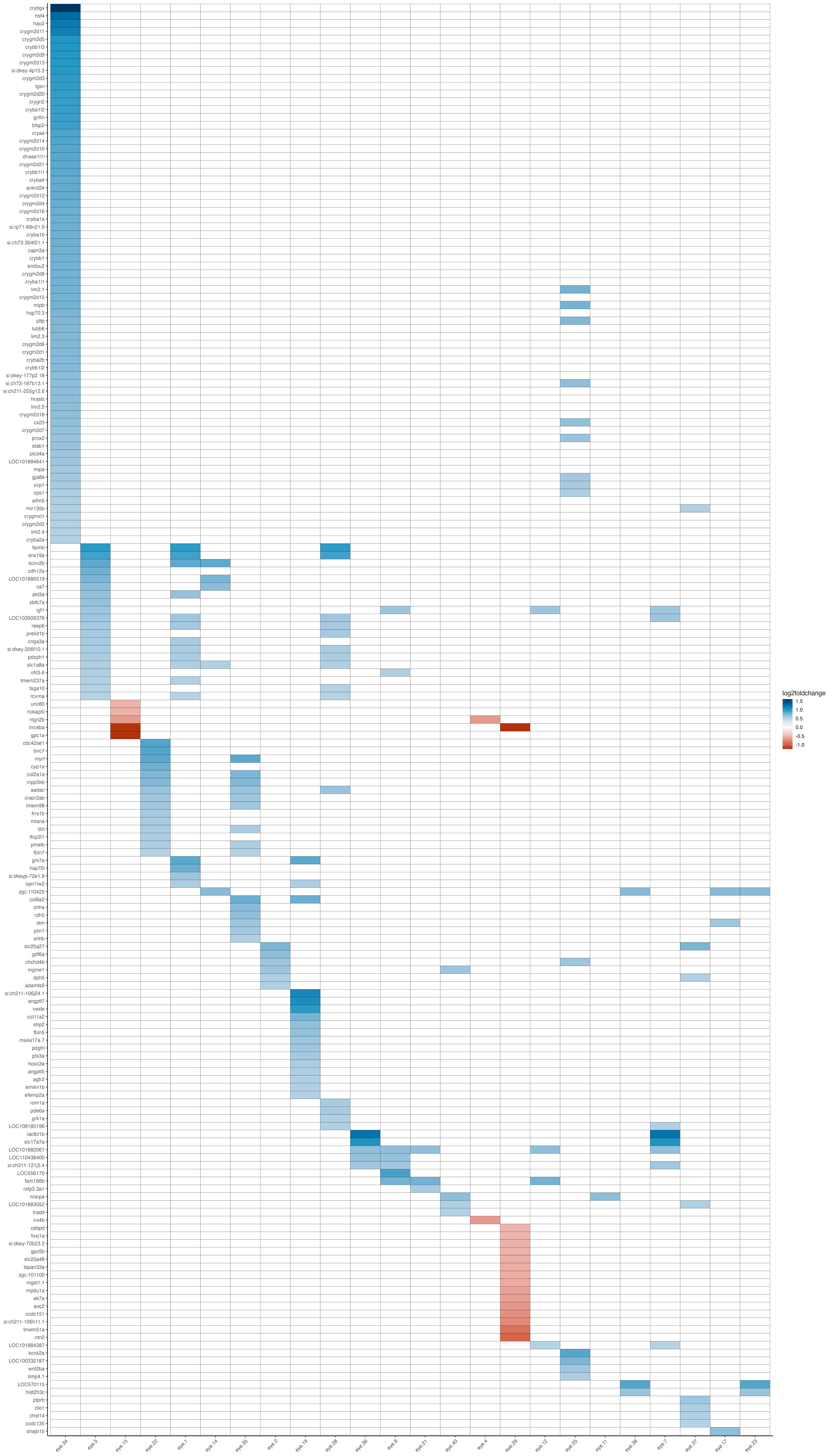

### ebf3a-2dpf-homvshetandwt_glia_scRNA-danio-byp_heatmap.png

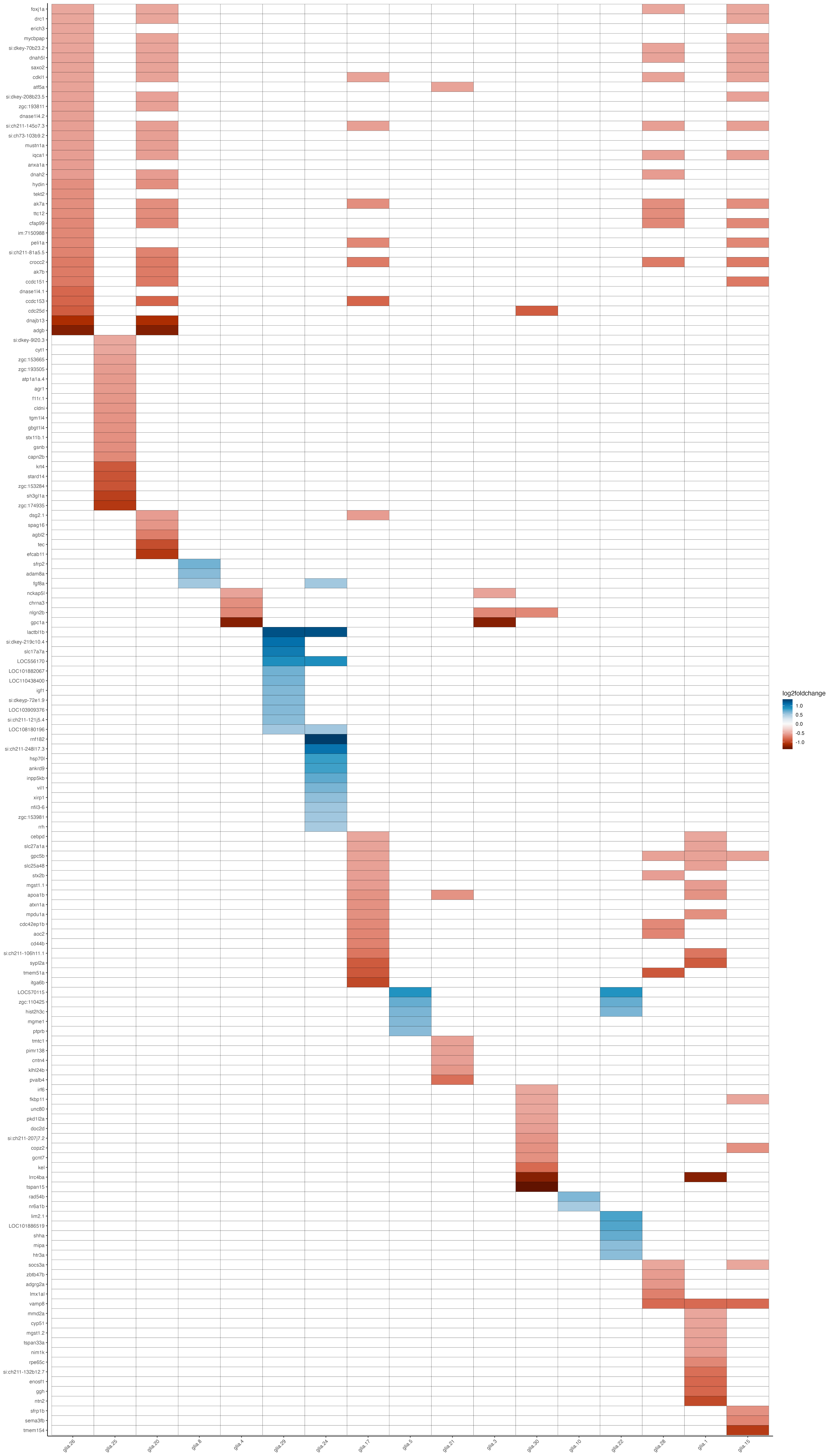

### ebf3a-2dpf-homvshetandwt_hema_scRNA-danio-byp_heatmap.png

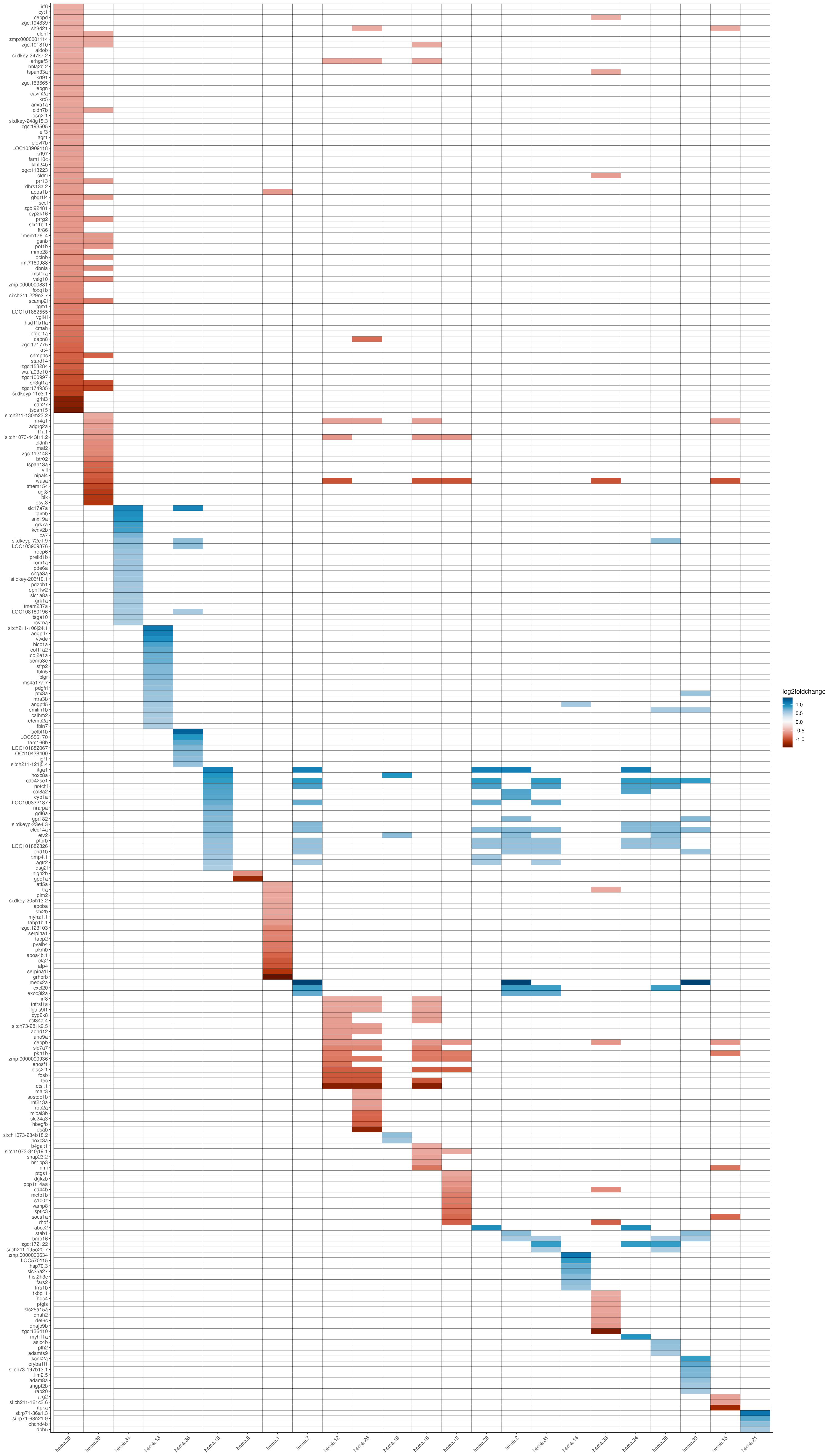

### ebf3a-2dpf-homvshetandwt_iono_scRNA-danio-byp_heatmap.png

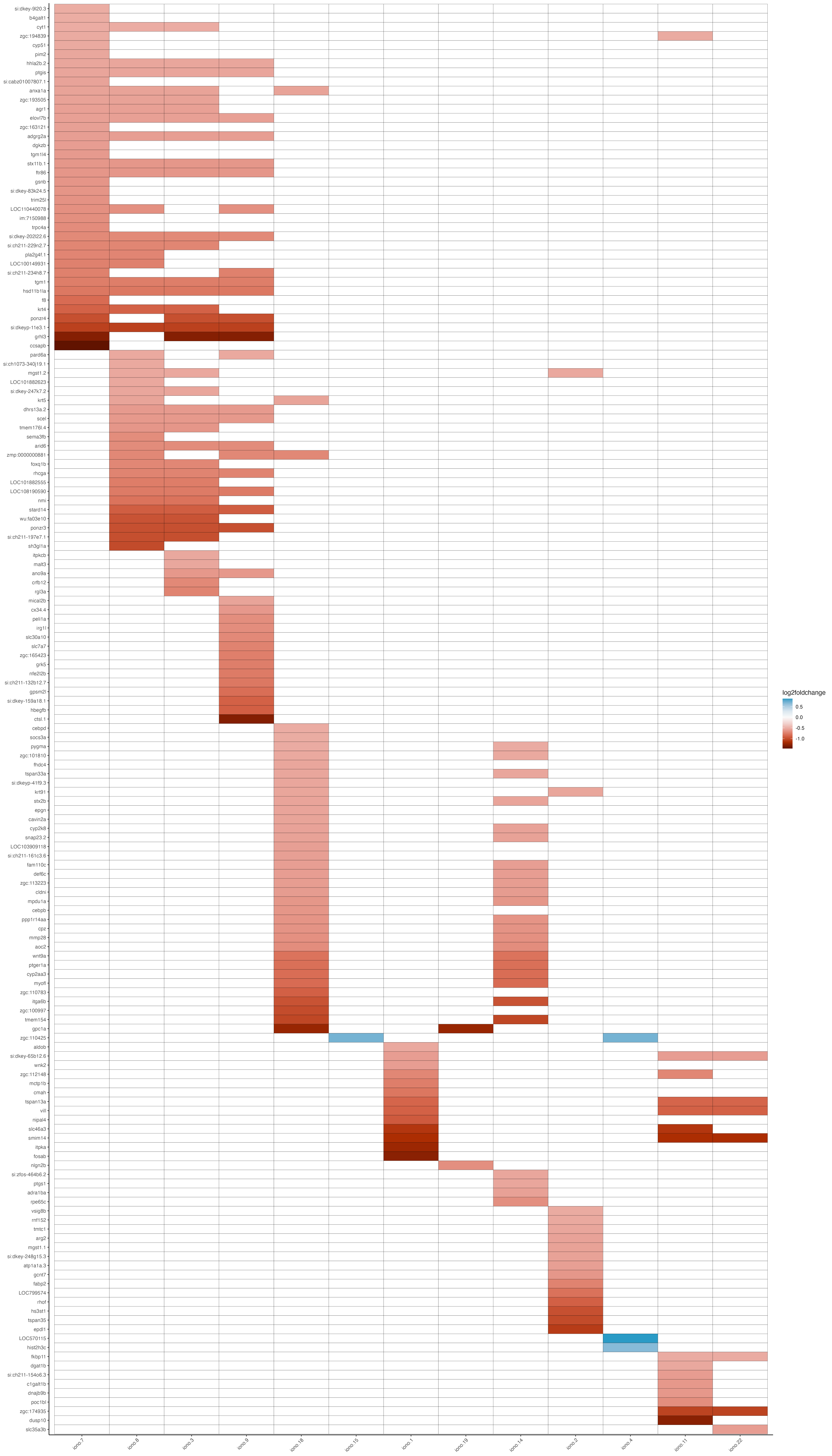

### ebf3a-2dpf-homvshetandwt_neur_scRNA-danio-byp_heatmap.png

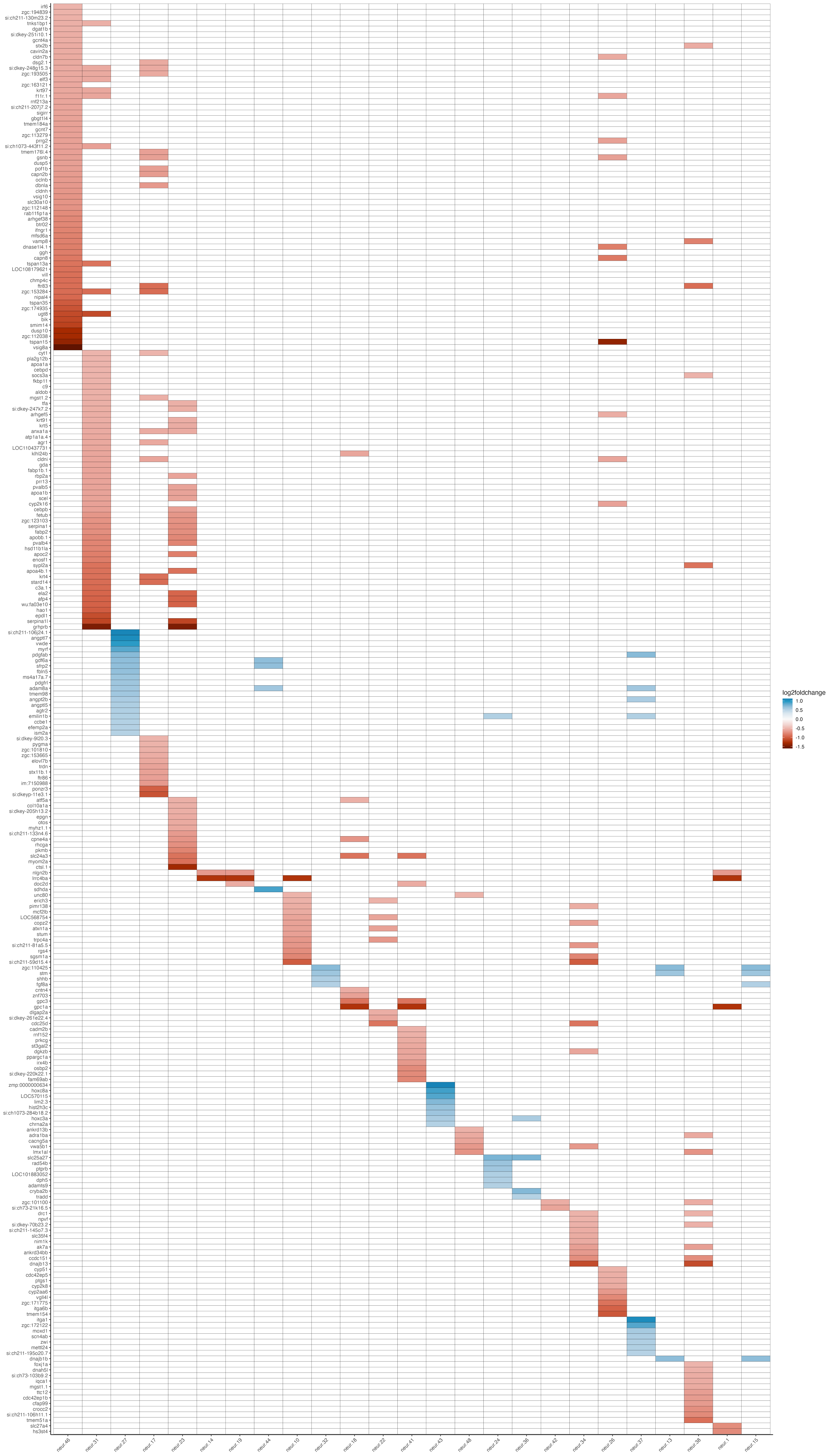

### ebf3a-2dpf-homvshetandwt_otic_scRNA-danio-byp_heatmap.png

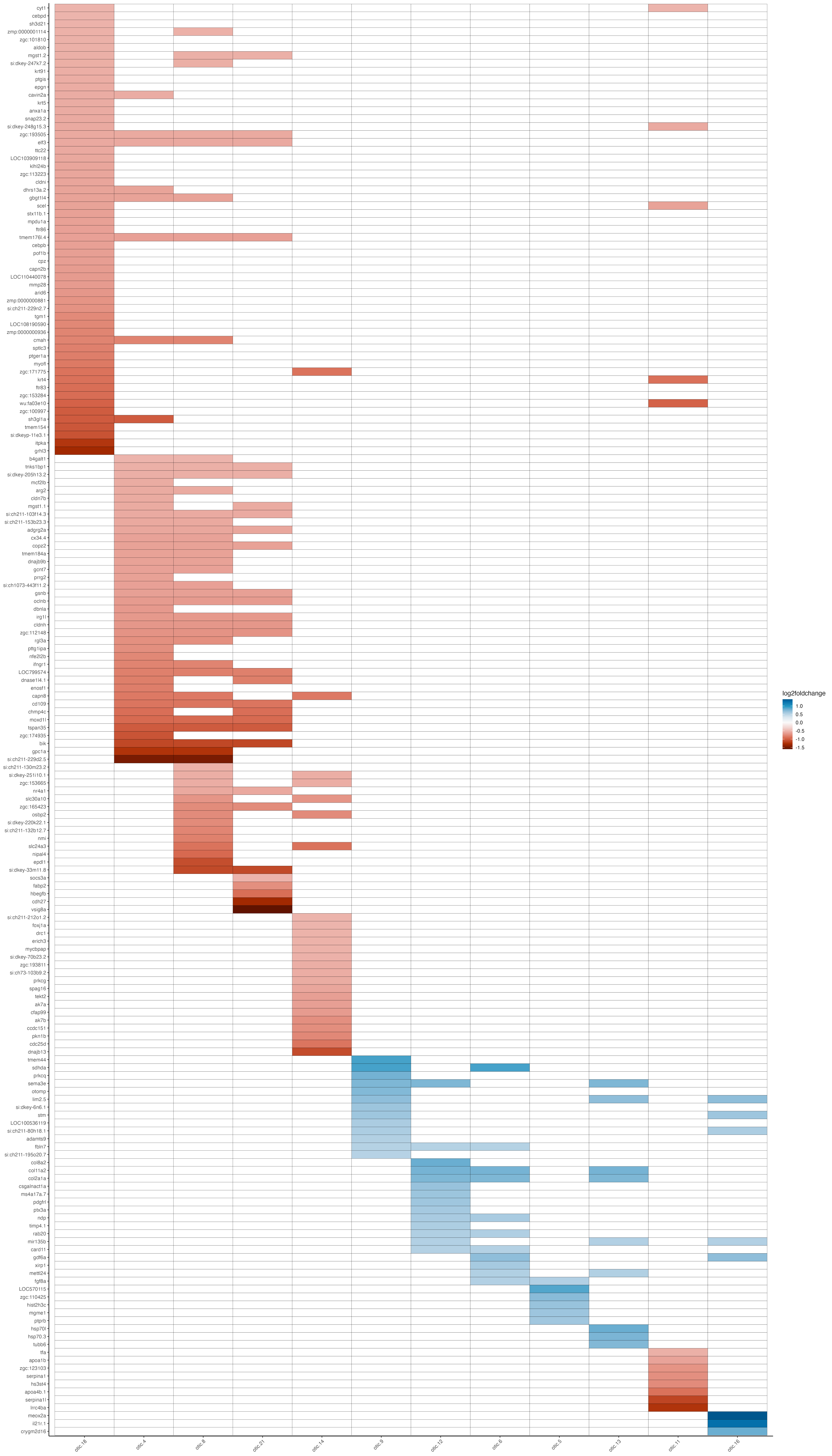

### ebf3a-2dpf-homvshetandwt_tast_scRNA-danio-byp_heatmap.png

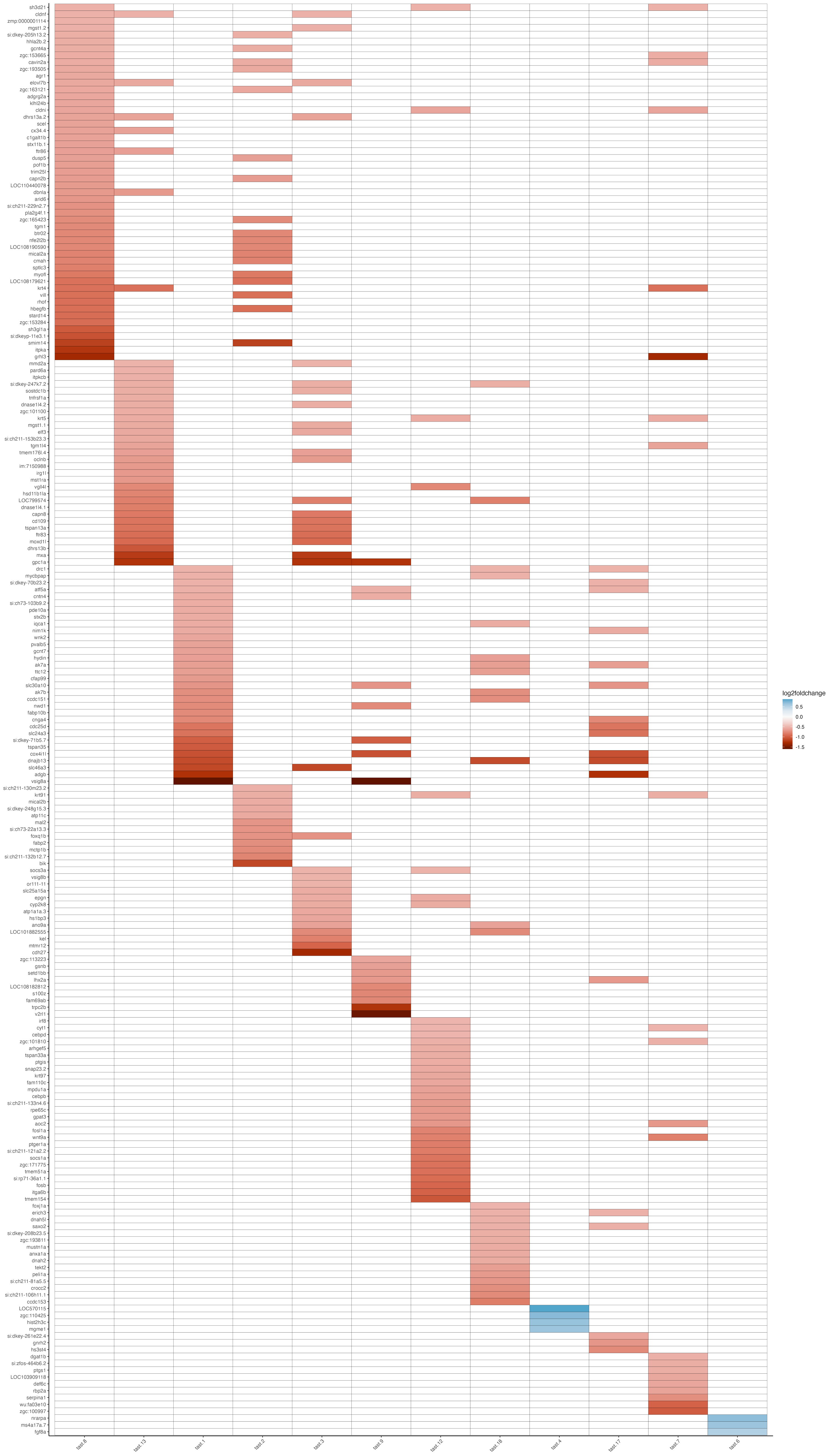
